# GIFT – A Global Inventory of Floras and Traits for macroecology and biogeography

**DOI:** 10.1101/535005

**Authors:** Patrick Weigelt, Christian König, Holger Kreft

**Affiliations:** Biodiversity, Macroecology & Biogeography, University of Goettingen, Büsgenweg 1, 37077 Göttingen, Germany

**Keywords:** Angiosperms, Functional biogeography, Functional traits, Plant checklists, Species composition, Vascular plants

## Abstract

To understand how traits and evolutionary history shape the geographic distribution of plant life on Earth, we need to integrate high-quality and global-scale distribution data with functional and phylogenetic information. Large-scale distribution data for plants are, however, often restricted to either certain taxonomic groups or geographic regions. For example, range maps only exist for a small subset of all plant species and digitally available point-occurrence information is strongly biased both geographically and taxonomically. An alternative, currently rarely used resource for macroecological and botanical research are regional Floras and checklists, which contain highly curated information about the species composition of a clearly defined area, and which together virtually cover the entire global land surface. Here we report on our recent efforts to mobilize this information for macroecological and biogeographical analyses in the GIFT database, the Global Inventory of Floras and Traits. GIFT integrates plant distributions, functional traits, phylogenetic information, and region-level geographic, environmental and socioeconomic data. GIFT currently holds species lists for 2,893 regions across the whole globe including ~315,000 taxonomically standardized species names (i.e. c. 80% of all known land plant species) and ~3 million species-by-region occurrences. In addition, GIFT contains information about the floristic status (native, endemic, alien and naturalized) and takes advantage of the wealth of trait information in the regional Floras, complemented by data from global trait databases. Based on a hierarchical and taxonomical derivation scheme, GIFT holds information for 83 functional traits and more than 2.3 million trait-by-species combinations and achieves unprecedented coverage in categorical traits such as woodiness (~233,000 spp.) or growth form (~213,000 spp.). Here we present the structure, content and automated workflows of GIFT and a corresponding web-interface (http://gift.uni-goettingen.de) as proof of concept for the feasibility and potential of mobilizing aggregated biodiversity data for global macroecological and biogeographical research.

## Introduction

Worldwide, about 382,000 vascular plant species form the basis of our terrestrial biosphere and provide key ecosystem services to humanity (Willis, 2017). Despite the long history of botanical exploration of our planet, the global distribution is only known for a subset of all plant species at comparatively coarse spatial grains (e.g. WCSP, 2012). In contrast to smaller and better known taxa like birds and mammals (BirdLife International, 2018; IUCN, 2018), high-quality species-level range maps or atlas data of plants are only available for selected groups (e.g. conifers in Farjon & Filer, 2013; cacti in Barthlott *et al.*, 2015) or regions (e.g. Europe in Tutin *et al.*, 1964–1980; woody plants of China in Fang *et al.*, 2011). Many research questions at the forefront of biogeography and macroecology, however, require a detailed knowledge of global plant distributions and, additionally, of species-level functional traits and phylogenetic relationships (e.g. Morueta-Holme *et al.*, 2013; Weigelt *et al.*, 2015; König *et al.*, 2017).

Several national and international initiatives focus on mobilizing and aggregating plant distribution information. For instance, the Global Biodiversity Information Facility (GBIF, 2018), provides access to ~214 million point occurrences of vascular plant species. These records are invaluable for plant ecology and conservation-related research, as they provide information about key aspects of species identity, time and place (Powney & Isaac, 2015). However, taxonomic, geographical and temporal biases (Hortal *et al.*, 2015; Meyer *et al.*, 2016), as well as limited curation and standardization efforts and the lack of important meta-information, like, for example, the floristic status at a given location (native, non-native, naturalized, etc.), limit their usefulness for macroecological research. An alternative source of information are Floras and checklists which, in contrast, present highly curated accounts of the plant species known to occur in a certain region. Floras and checklists are often based on decades to centuries of exploration and regional botanical work, and have profited from the expertise of generations of botanists. They aim at providing (near-)complete floristic inventories for a given region and thus provide information on species presences and their floristic status, and additionally allow for the inference of local species absences (Lobo *et al.*, 2010; Jetz *et al.*, 2012). So far, extensive compilations of plant checklists exist only for certain geographic regions (e.g. Ulloa Ulloa *et al.*, 2017), taxonomic groups (e.g. Flann, 2009; WCSP, 2012), functional types (e.g. BGCI, 2017), or, for example, naturalized alien plants (van Kleunen *et al.*, 2015; Pyšek *et al.*, 2017).

In light of the increasing availability of biodiversity data, it is a major challenge to integrate various data types and to link data from different ecological domains representing species distributions, functional traits, phylogenetic relationships or environmental characteristics for analyses and cross-validation (König *et al.*, in prep.). Initiatives that integrate different types of distribution data with additional biotic or abiotic information are currently most comprehensive for particular geographic regions (e.g. BIEN for the Americas; Enquist *et al.*, 2016) or other taxa (e.g. Map of Life for vertebrates; Jetz *et al.*, 2012). However, the wealth of aggregated information in regional Floras and checklists (Frodin, 2001) allows for a near-global characterization of plant distributions. In combination with functional traits from the botanical literature or large trait databases (e.g. RBG Kew, 2008; Kattge *et al.*, 2011b) and ever-growing species-level phylogenies (e.g. Smith & Brown, 2018), this represents a promising basis for macroecological and biogeographic research.

Here, we present GIFT, the Global Inventory of Floras and Traits database, a new resource designed to integrate species distribution data and species-level functional traits of plants from regional Floras and checklists with phylogenetic information and geographic, environmental and socio-economic characteristics (**Fig. 1**). As such, the database architecture, workflows and data of GIFT facilitate a wide array of macroecological and biogeographical analyses and may help to extent and validate other plant distribution and trait data resources. The general concepts outlined here may serve as a role model for aggregated species checklist and trait databases for other major taxonomic groups.

**Fig. 1.**
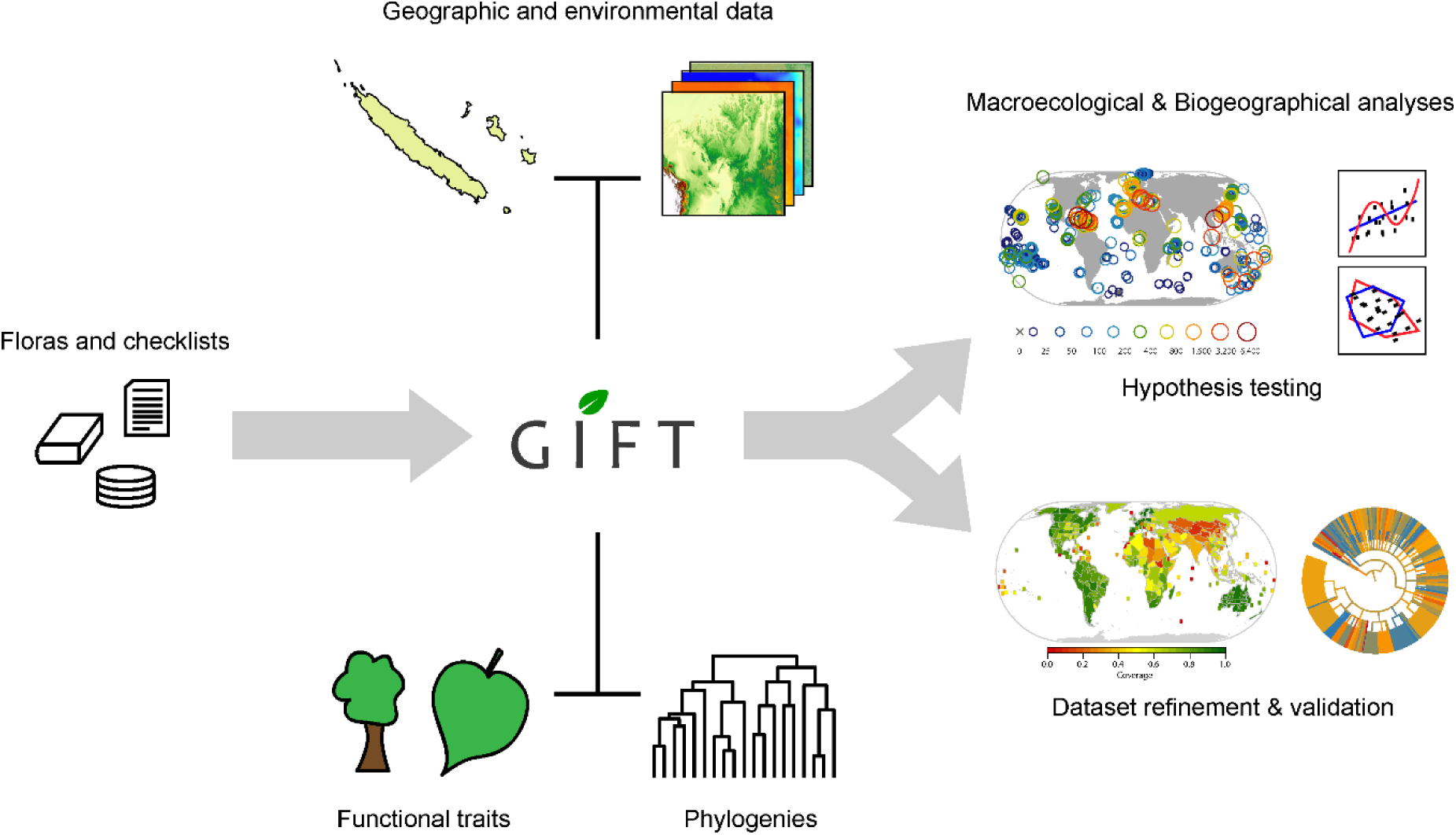
Conceptual framework of the Global Inventory of Floras and Traits database (GIFT). The core information in GIFT are species occurrences in geographic regions (islands, political units, protected areas, biogeographical regions) based on Floras and checklist. At the level of the geographical regions, this information is linked to physical geographic, bioclimatic and socioeconomic properties. At the level of the species, functional traits, taxonomic placement and phylogenetic relationships are linked. This integration of species distribution data in the form of full regional inventories and regional and species characteristics allows for a wide variety of macroecological and biogeographical analyses of taxonomic, phylogenetic and functional diversity as well as for the refinement and validation of other plant distribution and trait datasets.

## Content and structure of GIFT

### Overview

Regional Floras and checklists are a rich source of information on species distributions that often also contain detailed descriptions of species traits and other information such as conservation status and human uses (Palmer & Richardson, 2012). In the past, botanical knowledge was recorded primarily in printed books (Frodin, 2001), which are labor-intensive to convert into structured data. These resources, however, are increasingly being made digitally available (e.g. Zuloaga *et al.*, 2004; Acevedo-Rodríguez & Strong, 2007) and modern regional inventory projects are set up as digital databases right from the start (e.g. Brach & Song, 2006; Jardim Botânico do Rio de Janeiro, 2016). In GIFT, we make use of this wealth of information, and collate and mobilize plant species lists and trait information from published and unpublished Floras, catalogs, checklists and online databases into a single integrated and curated global database.

The original checklist data in GIFT consist of species names from the literature, their occurrences in the regional species lists and original trait information (yellow boxes in **Fig. 2**). All this information is linked to meta-data on the included literature references, species lists, traits and geographic entities (white boxes in **Fig. 2**). Semi-automated workflows allow a fast and reliable integration of new datasets and provide extensive curated and derived information (blue boxes in **Fig. 2**): (1) taxonomic match-up with taxonomic resources and name standardization (section ‘Species names and taxonomic standardization’), (2) taxon placement according to a taxonomic backbone and phylogeny (section ‘Taxonomic backbone and phylogeny’), (3) trait standardization and hierarchical and taxonomic trait derivation (section ‘Functional traits’), (4) calculation of regional summary statistics like species richness or trait coverage (section ‘Geographic regions’) and (5) extraction and computation of geographic, environmental and socioeconomic regional characteristics (section ‘Geographic regions’). Based on this generic database framework, GIFT can be queried for complete species checklists of a certain taxonomic or functional group and floristic status (e.g. ‘native angiosperms’ or ‘naturalized trees’). Alternatively, it is possible to extract distributional and functional information for a set of species, or to extract environmental information, species numbers and mean trait values for a set of regions. GIFT is currently stored on a MySQL 5.5.43 database server. Workflows for preparing, importing, processing, extracting and visualizing data are written in the R statistical programming language (R Core Team, 2018).

**Fig. 2.**
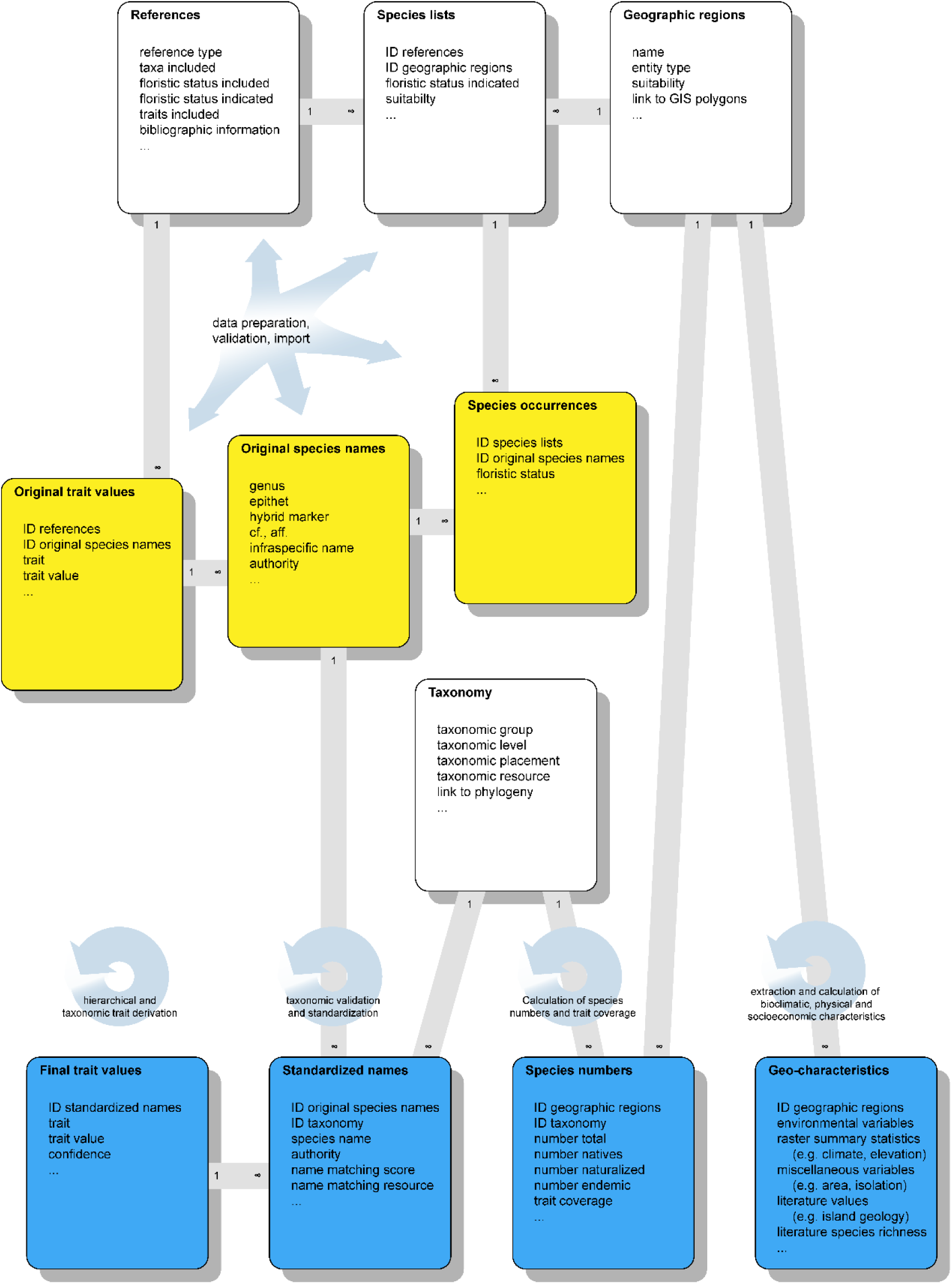
Simplified structure of the Global Inventory of Floras and Traits database (GIFT). Meta-data on literature references, species lists and geographic regions builds the backbone of the database (top row). A reference can include several species lists (e.g. for different sub-regions) and a geographic region can be covered by several lists and references. Species lists vary in taxonomic and floristic scope (e.g. all native and naturalized angiosperms) and in the information content (floristic status, functional traits). Primary occurrence information, species names and functional trait data from the literature resources build the main block of original data (yellow). Automated workflows link those to taxonomically standardized working names, to a higher taxonomy and phylogeny of vascular plants and produce derived resources for analyses (blue). Grey bars indicate links among tables in a simplified way (most tables shown here represent several tables in the database).

### Checklists

GIFT currently contains 3,826 species lists referring to 2,893 different geographic regions which are based on 429 original checklist data sources (**Fig. 2**). Compared to other plant diversity databases which either focus on other types of distribution data (point occurrences, vegetation plots, range maps) or trait data (individual measurements), this represents a complementary resource of unprecedented extent. A full and up-to-date list of all data citations and their bibliographic references is available in **Appendix S 1** and at the GIFT website (http://gift.uni-goettingen.de), and publications based on GIFT are requested to cite the checklist resources the analyses are based upon (e.g in Weigelt *et al.*, 2016). Checklists and inventories stem from publically available sources as well as from unpublished sources with restricted access (4.2%). Meta-data on references and species lists further specify the type of the reference as provided (Flora, checklist, catalogue, identification key, survey, etc.), the taxonomic and floristic scope of the reference (e.g. all native and naturalized angiosperms), whether the species’ floristic status is indicated and which functional traits are reported.

The actual distribution information is kept in a separate table that links the taxonomic names to the species lists they occur in and via those to geographic regions (**Fig. 2**, ‘species occurrences’). For each species occurrence, we indicate, if known, whether this occurrence is native or not. For native species, we further indicate if species are endemic to the geographic entity of the species list or to the geographic entity of the entire reference, if indicated in the literature source. For non-native species, we additionally indicate whether they are naturalized or not (Richardson *et al.*, 2000). We also indicate whether the occurrence and the different kinds of floristic status information are questionable or doubtful according to the literature source as binary variables. Via the species names, occurrences are linked to species-level functional traits as well as to the taxonomic and phylogenetic backbone. Via the geographic regions, species and traits are linked to regional geographical characteristics (**Fig. 2**). Routines to export checklists from GIFT and their meta-data as Darwin Core (Wieczorek *et al.*, 2012; Wieczorek *et al.*, 2014) and Humboldt Core (Guralnick *et al.*, 2018) archives, respectively, are currently being developed to allow easy integration with other kinds if distribution data (e.g. point occurrences from GBIF).

### Species names and taxonomic standardization

All species names enter the database in their original form including infraspecific information and author names where available. Species names derived from heterogeneous resources, referring to various geographic regions and published over a timespan of about one hundred years, inevitably vary in the taxonomic concepts applied (Jansen & Dengler, 2010). To compare species identities across different resources, we therefore submit all non-hybrid species names to a semi-automated taxonomic standardization and validation procedure based on taxonomic information provided by The Plant List 1.1 (TPL; The Plant List, 2013) and additional resources available via iPlant’s Taxonomic Name Resolution Service (TNRS; Boyle *et al.*, 2013). Names of hybrid species are currently not standardized due to heterogeneous formats of their scientific names. The original names are nonetheless stored for further processing if needed. This procedure was exclusively developed to meet the needs of the GIFT database and has already been applied and described in Meyer et al. (2016).

First, all genus names not occurring in TPL are checked manually and spelling mistakes are corrected based on literature and online resources (e.g. Mabberley, 2008; IPNI, 2012). Entries that cannot be assigned to an established genus name at all (valid or not) are excluded from further steps. Second, all species names are compared automatically to all taxonomic names available for a particular genus in TPL based on pairwise orthographic distances (generalized Levenshtein distance; Levenshtein, 1966) between species epithets, infraspecific names, author names and the entire species names. We use both the absolute orthographic distance, which is the number of changes needed to transform one character string into the other (Levenshtein, 1966), and the relative orthographic distance, which relates the absolute orthographic distance to the length of the longer input string. Based on the orthographic distances of an original species name to all congeneric species listed in TPL, we determine the final working name hierarchically: First, we choose the best-matching species epithet. If multiple epithets match equally well, we choose those with best-matching infraspecific names (if infraspecific name available and if absolute orthographic distance < 4 and relative orthographic distance < 0.3), and then those with best-matching author names (if author names available and relative orthographic distance < 0.5). The specific matching thresholds at each step were derived seeking a balance between the number of names that cannot be matched and the number of names that are matched to the wrong species. Synonyms are linked to their accepted species names as suggested by TPL. If several names match equally well and lead to different accepted binomial species names, we first remove illegitimate and invalid names, then synonyms and then accepted names with poorer overall orthographic distance. In addition, all names are resolved using the TNRS application programming interface (API), which returns similar statistics on the name matching and the status of the matched names like the above-described approach using TPL. For choosing standardized binomial working names we give priority to TPL over TNRS, because of the possibility of adjusting our TPL name matching approach. If a name does not match any name via TPL or TNRS with a relative orthographic distance < 0.25 for either the epithet or the full name, we keep the original name as working name. If not stated otherwise summary statistics below are based on these standardized binomial working names.

All original names, orthographic distances, matched names and meta-information about the matching are stored in the database (**Fig. 2**). Thus, the taxonomic standardization in GIFT is fully transparent and repeatable whenever taxonomic resources are updated or extended. Moreover, the stored information can be used to filter out names that did not match, matched only to a certain degree, or that do not lead to an accepted species name, allowing for rigorous sensitivity analyses of the effects of taxonomic uncertainties on the outcome of macroecological and biogeographical analyses.

### Taxonomic backbone and phylogeny

All species working names are linked to a taxonomic backbone via their genus names. The taxonomy is based on the Angiosperm Phylogeny Group IV system for angiosperms (The Angiosperm Phylogeny Group, 2016), and on the Angiosperm Phylogeny Website version 13 (Stevens, 2013) and The Plant List 1.1 for gymnosperms, pteridophytes and bryophytes (The Plant List, 2013). Based on the taxonomic backbone, the database can be queried in two directions. First, species lists can be extracted including only species that belong to a certain taxon (e.g. only angiosperms). Second, geographic units can be chosen for which species lists cover a complete taxon of interest (e.g. all regions with Bixaceae checklists). In combination, species lists of a certain taxon can be produced for all regions where the required data is available. In addition, species-level functional traits can be aggregated at any desired taxonomic level and trait information for broad taxonomic groups can be used to derive species-level information for traits that are consistent across a larger taxonomic groups.

All seed plant species are linked to a global phylogeny with 353,185 terminal taxa (Smith & Brown, 2018) for phylogeographical analyses. Two versions of this phylogeny are included in the database in tabular form to extract checklist and trait information for particular clades and to visualize trait and taxonomic coverage across the phylogeny. In one version, species in GIFT not included in the phylogeny were added replacing all members of the genera they belong to with polytomies (Pearse *et al.*, 2015) and in the other version missing species were excluded to keep detailed phylogenetic relationships among the species covered by the phylogeny. In addition, all vascular plant species in GIFT are linked to a phylogeny with fewer terminal taxa but broader phylogenetic extent (i.e. including pteridophytes; Qian & Jin, 2016), which was used here to assess taxonomic coverage of distribution and trait data at the family level.

### Functional traits

Species in GIFT are linked to functional trait information from currently 155 original resources. Most trait resources are Floras and checklists for which annotated information on traits have been extracted, but also large trait compilations with or without spatial context are incorporated (e.g. Zotz, 2013; BGCI, 2017). The range of functional traits currently covered by GIFT reflects different aspects of plant morphology, life history, reproduction, physiology, genetics, and ecology (**Table S 1**). The focus lies on aggregated trait information at the species level, making GIFT a valuable complementary resource to initiatives that collate large amounts of trait measurements at the individual level (e.g. Kattge *et al.*, 2011b; Enquist *et al.*, 2016).

Many trait resources provide equivalent information in various languages, using different terminologies or measurement units. The first step of trait processing in GIFT is therefore the standardization of primary trait data according to pertinent trait literature (Pérez-Harguindeguy *et al.*, 2013; Garnier *et al.*, 2017) (**Fig. 3**) using defined categorical levels (categorical traits) and units of measurement (numerical traits) (**Table S 1**). To retain the maximum information provided by the original resources, many categorical traits are defined at multiple levels of detail (e.g. life form 1 vs. life form 2). In the second trait processing step, the standardized trait values are therefore subjected to a hierarchical derivation procedure (**Fig. 3**). This procedure makes use of logical links and nestedness among many functional traits based on their definitions. For example, the value “tree” in the trait growth form implies the value “woody” in the trait woodiness. As such hierarchical trait derivation increases the amount of trait information in GIFT. In addition, hierarchical derivation ensures compatibility among different levels of detail of the same trait by automatically deriving values in coarser variations from available information in more detailed ones. We organize such hierarchical relationships between traits in a directed graph that can be easily traversed to fill data gaps (**Fig. S 1**). A tabular version of the graph is stored in the database for modifications (**Table S 2**). A similar derivation approach is implemented for taxonomic groups that are uniform with respect to a particular trait (taxonomic trait derivation). In this case, the basis of derivation is not the logical hierarchy of trait values, but that of taxonomic groups. For example, the genus *Abies* consists exclusively of monoecious trees (Farjon, 2010). Thus, all species of *Abies* can be characterized with respect to growth form “tree” and reproductive mode “monoecious” based only on their taxonomic position. Subsequently, the taxonomically derived species-level traits are subjected to the hierarchical derivation as outlined above.

**Fig. 3.**
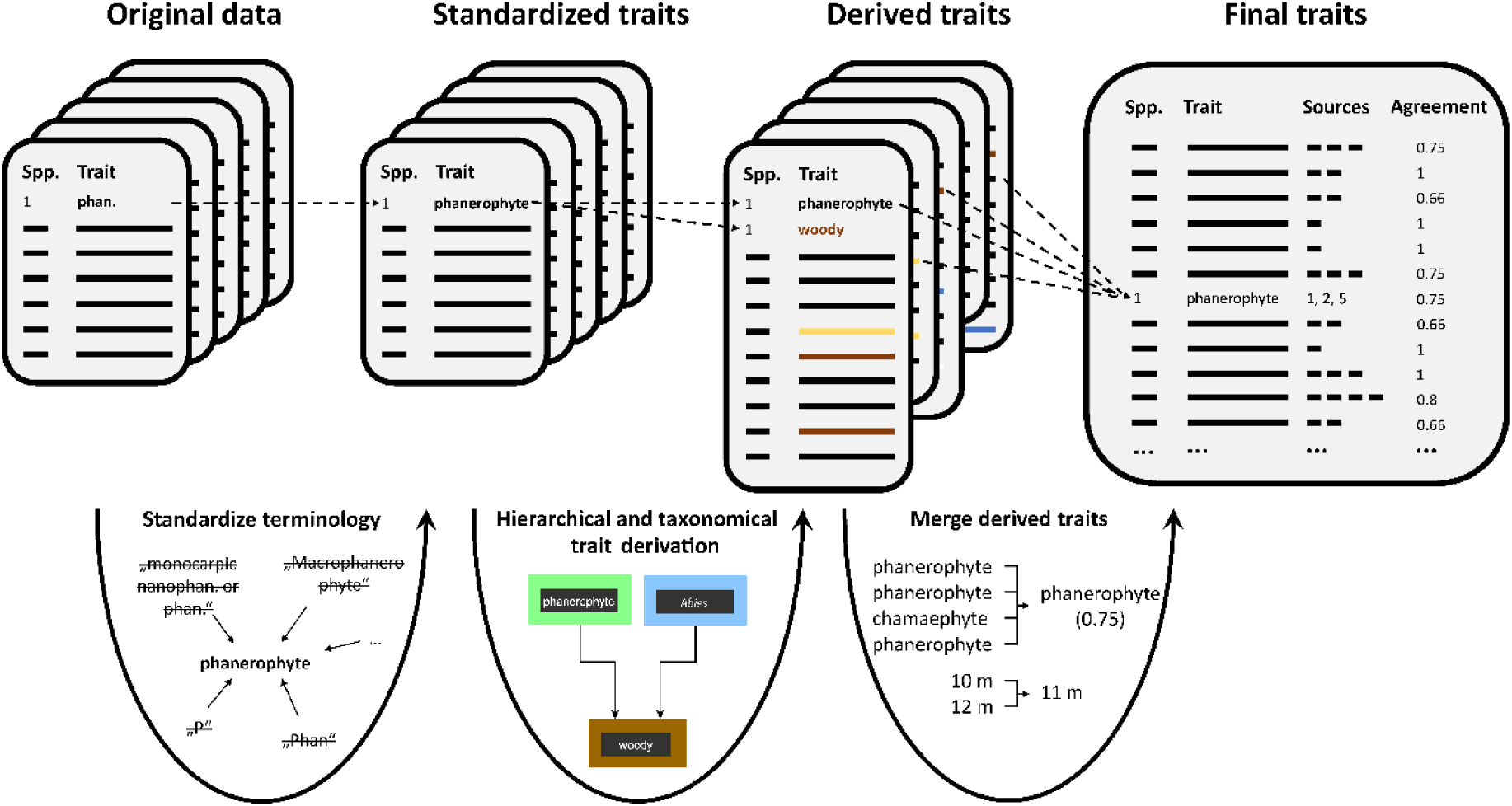
Trait processing in GIFT. Original trait records entering GIFT are subjected to three processing steps: (1) Trait values are standardized with respect to language, terminology and measurement unit. (2) Additional trait values are derived hierarchically for traits that are logically nested (**Fig. S 1**), and taxonomically for species that belong to taxonomic groups that are uniform with respect to a particular trait. (3) Derived trait values are aggregated at the species level based on the consensus among resources (categorical traits) or summary statistics are computed based on the original values (numerical traits).

The original and derived trait data in GIFT may include several values per trait-species combination. To obtain a single, standardized value per trait and species, original and derived trait values are aggregated based on the consensus among resources (categorical variables, 66 % consensus threshold) or summary statistics are computed based on the original values (numerical variables, currently: mean, minimum and maximum) (**Fig. 3**). Trait derivation and aggregation for a given species are repeated each time a new trait record enters the database, such that the final values and consensus rates are continually re-evaluated in the light of new information. Throughout the entire procedure of trait processing, information can be traced back to their original reference and unstandardized value. This provides a basis for implementing advanced gap-filling (Schrodt *et al.*, 2015) and aggregation techniques (Kattge *et al.*, 2011a) in the future.

### Geographic regions

As indicated in the original literature resources, each regional species checklist in GIFT unambiguously refers to a geographic region, e.g. an island, archipelago, a political or biogeographical unit, or a protected area. To make most out of the spatial delineation of the geographic regions, we assemble spatial polygons from the Biodiversity Information Standards Working Group (TDWG, 2007), the GADM database of Global Administrative Areas (Hijmans *et al.*, 2009), single island polygons extracted from GADM (see Weigelt *et al.*, 2013), the Global Island Database (UNEP-WCMC, 2013), the World Database on Protected Areas (UNEP-WCMC, 2014), or digitize regions manually according to the checklist literature resources. The regions vary in area by 13 orders of magnitude, ranging from small islands to large countries and botanical continents (**Fig. S 2**). Many regions may overlap with each other and small regions are frequently nested in larger ones (e.g. Yosemite National Park in California in the US). The degree of overlap is calculated automatically and recorded in the database to exclude overlapping entities or aggregate information for analyses.

For each original geographic region, a suite of 123 physical geographic, bioclimatic and socio-economic characteristics is computed for macroecological analyses based on the regions’ spatial information and additional spatial datasets (**Fig. S 3**, **Table S 3**). Specifically, this includes (1) characteristics based on the spatial polygon itself like its area, centroid coordinates and geographic extent, (2) summary statistics (15 quantiles including minimum, median and maximum, mean, standard deviation, mode, number of unique values, Shannon diversity and number of cells) derived from raster layers like digital elevation models (Danielson & Gesch, 2011), global climatologies (e.g. CHELSA; Karger *et al.*, 2017) or human population density (Doxsey-Whitfield *et al.*, 2015), and (3) miscellaneous metrics calculated from additional spatial resources like biogeographic region affinity (Takhtajan, 1986) or island isolation (Weigelt & Kreft, 2013) (**Fig. S 3**c).

For families and higher taxonomic groups, we automatically calculate the number of all species, native species, naturalized species and endemic species for all regions that are covered by checklists for the given combination of taxonomic group and floristics status. Additionally, we calculate for the same taxonomic groups and floristic subsets the percent trait coverage for all functional traits covered by the database. It is hence possible to extract and visualize for which regions and taxa what information in terms of species checklists and functional traits is available. Based on the various checklist resources available for each geographic region, we decide whether the checklist information should theoretically completely cover the whole native vascular or angiosperm flora. This, however, is only a rough and subjective estimate and given the huge amount of unexplored plant diversity especially in the tropics and only partial completeness of the according Floras, it needs further evaluation. To this end, we are currently incorporating species numbers and richness estimates from the literature (Frodin, 2001; Kreft & Jetz, 2007) into the database and develop workflows to compare them to the species numbers derived from the checklists. Regions deviating considerably from these literature values can be excluded from analyses if needed. The same way regional trait coverage can be used to exclude regions with little trait information from analyses on trait patterns.

### Versioning

This paper describes GIFT version 1.0. New data are incorporated into GIFT in chunks and each time new data are added or workflows are modified, a new version is released. Changes will be documented at http://gift.uni-goettingen.de/about. Old versions are backed up and can be restored to reproduce analyses carried out on older versions of the database.

## Current state

### Geographic coverage

Initially, GIFT started with the collection of Floras and checklists for oceanic islands and the basic workflows have been developed for various projects focusing on island plant diversity (Weigelt, 2015; Weigelt *et al.*, 2016). Island floras usually host a comparatively limited set of species and have clearly defined geographic boundaries. As such, they have attracted a lot of scientific interest in the past, leading to a high availability of island Floras and checklists. GIFT therefore offers a very comprehensive overview over the floristic composition of 1,845 of the world’s islands, which has already led to a variety of studies on island biodiversity patterns and their determinants (Weigelt *et al.*, 2013; Cabral *et al.*, 2014; e.g. Weigelt *et al.*, 2015; Weigelt *et al.*, 2016; Lenzner *et al.*, 2017). More recently, GIFT has been expanded to mainland floras, currently covering 1,048 distinct regions, to support comparative analyses of continental and insular floras (e.g. König *et al.*, 2017) and extensive studies of global plant diversity.

In total, GIFT currently includes 2,963,438 species-by-region occurrences for 315,164 standardized species names across 2,893 geographic regions covering the whole globe. For 92.7 % of the occurrences the native status and for 47.5 % endemism is reported. 2,062 have at least one checklist for all native vascular plants, together covering all floristic kingdoms and biomes and 79.1 % of the earth’s land surface excluding Antarctica (**Fig. 4a**). After removing overlapping entities to avoid pseudo-replication, up to 1,841 regions and 58.2 % land surface coverage remain when prioritizing small entities (> 100 km²) over large entities, and 1,555 regions and 73.1 % land surface coverage remain when prioritizing large entities over small entities (single islands always prioritized over island groups). Geographic coverage varies with focal taxonomic group (**Fig. 4**, **Table 1**) and floristic status (**Fig. S 4**), and is highest for native species. The largest gaps for native vascular plant floras are currently located in Tropical Africa, the Middle East, Central Eurasia, and South East Asia (**Fig. 4**). Data gaps in GIFT do not necessarily represent true knowledge gaps. Floras of the countries of the former USSR, West Africa, Madagascar, Java and India, for example, are available and are currently in the process of being incorporated.

**Fig. 4.**
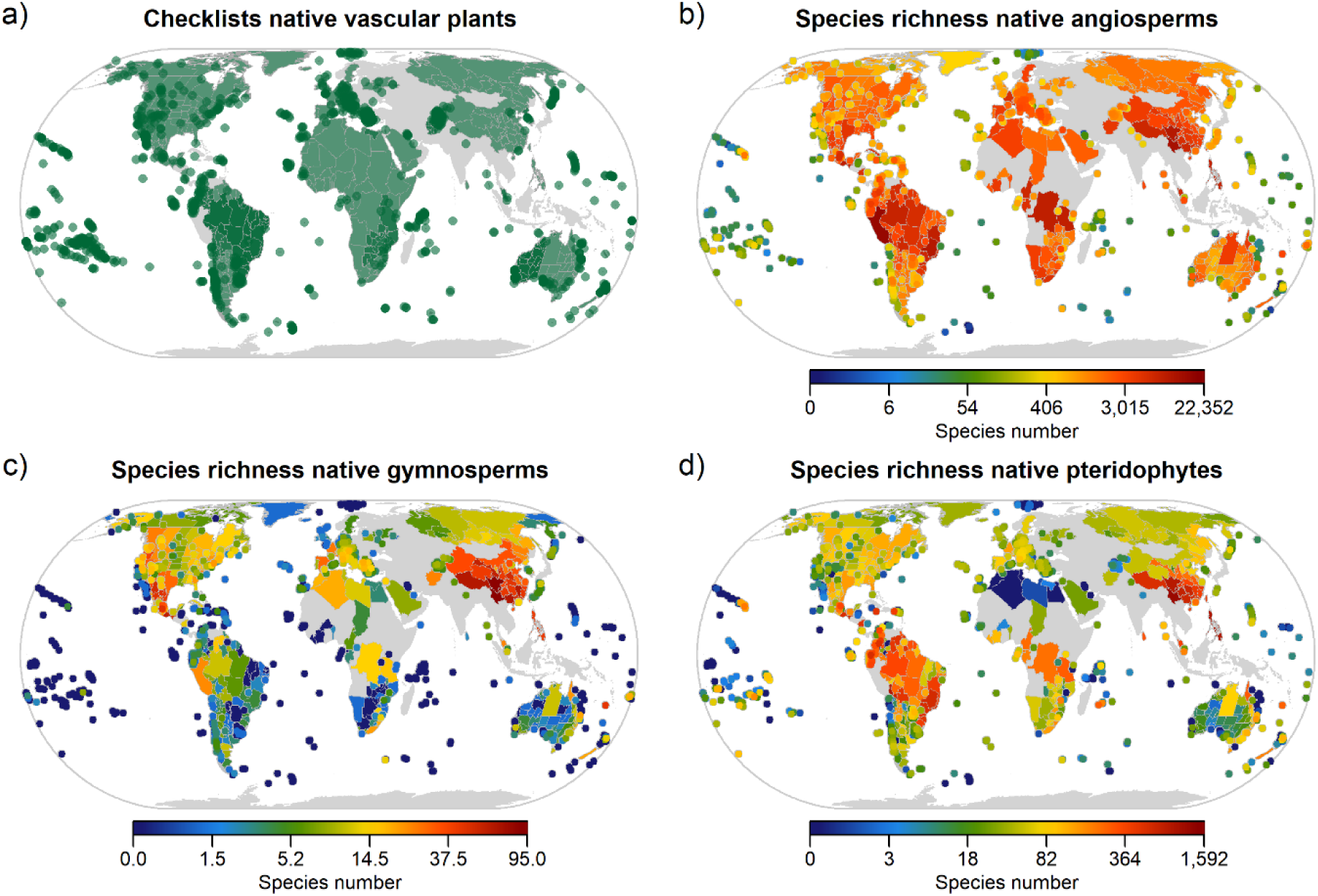
Spatial coverage of checklist data currently stored in GIFT. a) Regions with checklist data for native vascular plants. Darker green shade indicates overlapping regions. b-d) Checklist coverage and species richness of major taxonomic groups for regions with theoretically complete inventories. Polygons are plotted sequentially in order of decreasing area to show smaller regions on top of larger regions, in the case where they overlap. Regions <25,000 km² are plotted as points.

**Table 1.** Current content of GIFT for selected major plant groups in terms of number of regions with supposedly full inventories for native species, unstandardized taxonomic names, standardized species names, species with resolved taxonomy, and trait records.

| Taxonomic group | Regions | Names | Species | Species resolved | Trait records |
| --- | --- | --- | --- | --- | --- |
| Embryophyta | 53 | 717117 | 324136 | 277580 | 2307100 |
| Tracheophyta | 2062 | 714781 | 322002 | 275610 | 2306973 |
| Pteridophyta | 2079 | 24241 | 11888 | 8408 | 54772 |
| Gymnospermae | 2211 | 4031 | 1151 | 1051 | 12352 |
| Angiospermae | 2218 | 686509 | 308963 | 266151 | 2239849 |
| Orchidaceae | 2478 | 64508 | 28155 | 27029 | 192332 |
| Asteraceae | 2218 | 58492 | 27755 | 24450 | 167300 |
| Fabaceae | 2218 | 46999 | 21000 | 18416 | 145895 |
| Poaceae | 2431 | 38464 | 12368 | 11215 | 130492 |
| Rubiaceae | 2431 | 29485 | 14260 | 13545 | 96684 |
| Lamiaceae | 2431 | 18120 | 7882 | 7560 | 61708 |

Since many Floras refer to entire countries of various sizes, and some of the resources in GIFT use broad distributional classifications (e.g. WCSP, 2012; BGCI, 2017), many mainland regions in GIFT are relatively large (mean area = 170,287 km², median = 30,454 km²), especially in comparison to an average island (3,265 km²; **Fig. S 2**). However, GIFT also includes mainland regions of small sizes like protected areas and small political units, since smaller units span smaller environmental gradients, and thus provide a tighter link between taxonomic, functional and phylogenetic species composition and aggregated abiotic conditions (Pearson & Dawson, 2003).

### Taxonomic coverage

GIFT currently includes 324,136 taxonomically standardized species from all major groups of land plants (Embryophyta). 277,580 of which are resolved to accepted species names. The focus for the collection of species lists and traits lies on vascular plants (Tracheophyta, 322,002 species) and in particular on angiosperms (Angiospermae; 308,963 species; **Table 1**). On average, 79.3% of all accepted species per plant family according to TPL are covered by distribution data. Taxonomic coverage of distribution data does not show a significant phylogenetic signal (Abouheif’s Cmean = 0.03, p = 0.142; Abouheif, 1999), i.e. it exhibits no detectable bias towards certain clades (**Fig. 5**). The 324,136 species names in total derive from 717,117 unstandardized original names (after genus name correction and exclusion of hybrid names) that differ in spelling or in the availability of author names or infraspecific information. 98.2 % of all original names could be matched and standardized to an existing species name using our approach to match TPL or using the TNRS API. For 90.5 % of all names, the synonymy could be resolved. Only 3.6 % of all working names are names that were adopted unchanged from the original names because they could not be adequately matched to taxonomic resources.

**Fig. 5.**
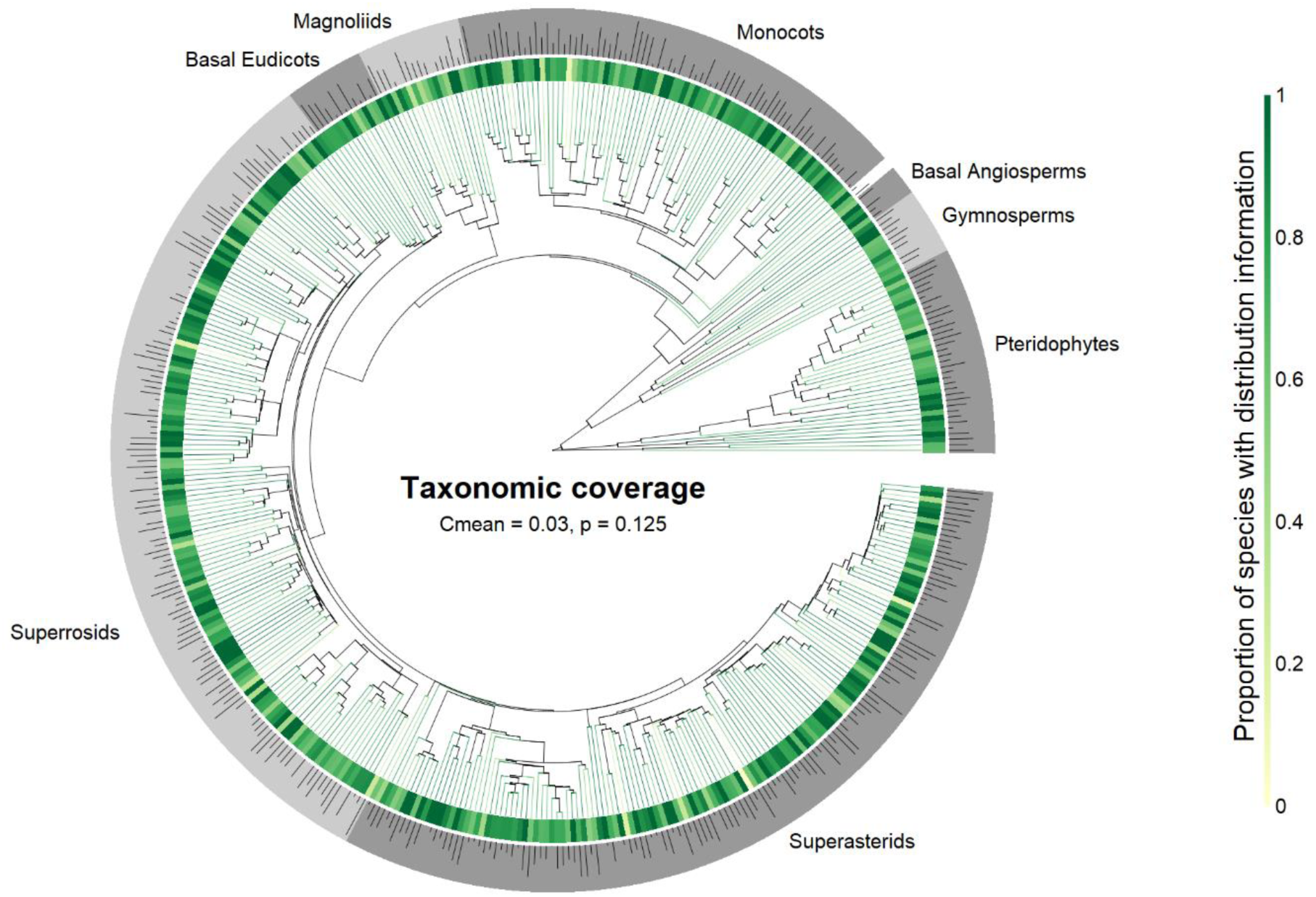
Taxonomic coverage of distribution data in GIFT at the family level. Tip color and inner ring color indicate the proportion of species with distribution information relative to all species of a given family, the grey outer ring delimits major clades of vascular plants. The height of bars in the outer ring is proportional to log10 total family species richness. Phylogenetic signal in taxonomic coverage was assessed as Abouheif’s Cmean, a measure of phylogenetic autocorrelation based on the sum of the successive squared differences between values of neighbouring tips in the phylogeny (Abouheif, 1999).

### Trait coverage

In total, there are 3,475,337 original trait records referring to 550,892 original taxon names. Hierarchical trait derivation yields an additional 1,261,718 trait records. After aggregating original and derived trait records, i.e. resolving species names and combining trait records for identical species, 2,307,100 species-trait combinations for 267,978 standardized species remain for ecological analyses (**Table 1**).

The majority of trait information in GIFT refers to morphological characteristics (**Table S 1**) such as woodiness (234,214 species) climbing habit (223,280 species), or growth form (213,372 species). Life history traits such as life form (100,607 species) or life cycle (84,206 species) are the second most common trait category. Other categorical traits are considerably rarer, e.g. photosynthetic pathway (31,534 species), dispersal syndrome (8,204 species), or pollination syndrome (4,511 species). Also quantitative traits such as maximum plant height (53,449 species), mean seed mass (23,874 species) or mean specific leaf area (2,304 species) are comparatively poorly covered. As such, GIFT represents a complimentary resource to existing trait databases which provide more records for numerical traits frequently measured in the field at the individual level (Pérez-Harguindeguy *et al.*, 2013), but have considerable lower coverage in terms of whole plant traits like, for example, growth form (84,459 species in BIEN 4.1 (Enquist *et al.*, 2016) and 99,217 species in TRY 4.1 (Kattge *et al.*, 2011b)) or life form (12,708 species in TRY 4.1).

To illustrate patterns in the geographic and taxonomic trait coverage of GIFT, we use the overall coverage across all traits as well as four exemplary traits (growth form, plant height, life form and seed mass). Geographically, most trait information per species is available in Europe and some comparatively species-poor temperate islands (**Fig. 6a**). Also, non-tropical parts of the Americas, Africa and Australia are well covered, whereas tropical regions in Africa and South-East Asia are least well covered with respect to their plant functional characteristics. However, geographic coverage varies strongly among individual functional traits. Frequent traits such as growth form are available for most species in almost every floristic region, whereas the coverage of less well-covered traits is strongly dependent on the geographic scope of the main contributing resources (**Fig. 6**). Life form *sensu* Raunkiær (1907), for example, is widely available throughout Europe, but rarely reported for species in other regions of the world (**Fig. 6d**). Likewise, plant height and seed mass exhibit uneven geographical coverage distributions, with highest coverage in Australia, South Africa and Europe (**Fig. 6c&e**).

**Fig. 6.**
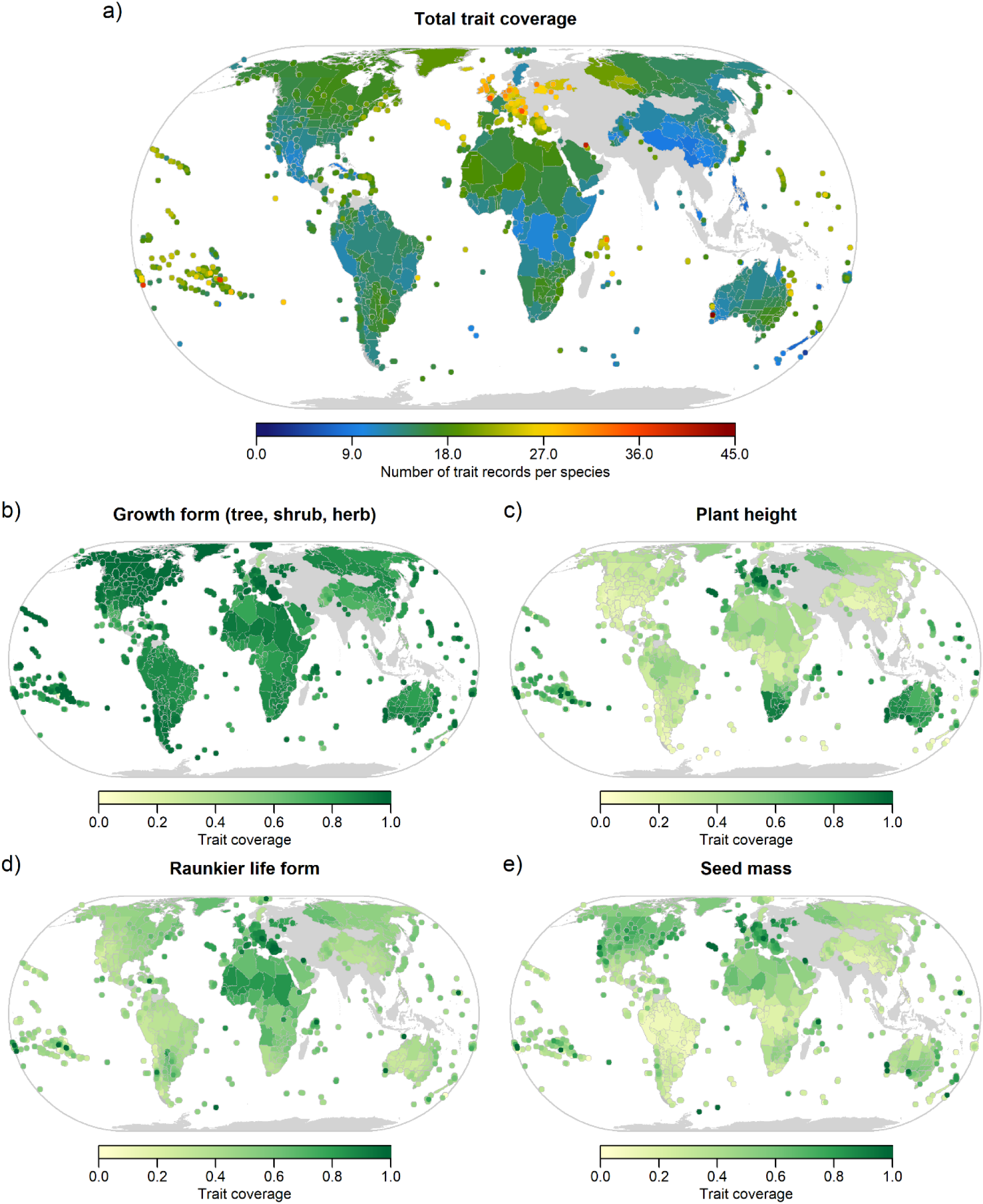
(a) Total number of trait records per native angiosperm species per region and (b-e) trait coverage per region (number of native angiosperm species with trait information/number of all native angiosperm species) for exemplary traits with characteristic geographic patterns in coverage. Polygons are plotted sequentially in order of decreasing area to show smaller regions on top of larger regions, in the case where they overlap. Regions <25,000 km² are plotted as points.

The taxonomic coverage of trait information in GIFT has small, yet significant phylogenetic signal (Cmean = 0.21, p < 0.001, **Fig. S 5**a). That is there is a mild bias towards certain taxonomic groups, e.g. monocots, in overall coverage of trait data. Examining phylogenetic signal at the level of individual traits reveals interesting patterns (**Fig. S 5**b-e). For example, plant height is very well covered for the graminid clade (leftmost group within the monocots, **Fig. S 5**c), and Raunkiær life form is particularly well covered in gymnosperms and monocots (**Fig. S 5**d).

### Web interface

A live overview over the current content of GIFT is available through a web interface at http://gift.unigoettingen.de. It provides summary statistics and allows producing customized richness and trait coverage maps for every combination of taxonomic group and floristic subset based on the species numbers and trait coverage values in the database. It is possible to see for which regions and taxa what information in terms of species checklists and functional traits is available and to browse the bibliographic references.

### Applications and outlook

Curated regional plant species composition data from Floras and plant checklists, and diverse information on species characteristics and their environment as integrated in GIFT (**Fig. 1**) allows to move global plant diversity research beyond using species richness as a proxy for biodiversity (Barthlott *et al.*, 2005; Kreft & Jetz, 2007; Kreft *et al.*, 2008). Examining the drivers of taxonomic, functional and phylogenetic diversity and turnover (Tuomisto, 2010; Qian *et al.*, 2013; Lamanna *et al.*, 2014; Weigelt *et al.*, 2015; König *et al.*, 2017) may help to disentangle the mechanisms underlying global plant diversity more directly (Graham *et al.*, 2014). Functional biogeography, for example, combines the mechanistic focus of functional ecology with the large eco-evolutionary scales of biogeography (Violle *et al.*, 2014) and thus provides a direct link between measures of organismal performance and a wide range of abiotic and biotic conditions. Although functional biogeographical approaches already provided significant insights into patterns and drivers of functional diversity (Moles *et al.*, 2014; Reichstein *et al.*, 2014; Engemann *et al.*, 2016; Butler *et al.*, 2017), the availability and representativeness of data on plant traits and distributions remains a limiting factor. Together with distribution and floristic status information available in GIFT (e.g. native, naturalized, endemic), functional traits may help to better understand the biogeographic history of plant life on Earth and its anthropogenic stressors. Analyses of endemic species and their traits, for example, can shed light on the evolution of new species and their contribution to current biogeographic patterns (Weigelt *et al.*, 2016). Naturalized alien species and their traits help to understand the role of humans in changing plant assemblages and may teach us how new habitats and regions are colonized. Knowledge on the composition of native vs. alien floras (see www.glonaf.org; van Kleunen *et al.*, 2015; Pyšek *et al.*, 2017) allows to tackle pressing questions in invasion ecology, for example what native floras are susceptible to plant invasions and how regional plant composition changes due to the naturalization of alien species (Winter *et al.*, 2009).

Apart from direct use as data source for macroecological or biogeographical research, GIFT is also a valuable resource to validate or expand other distribution or trait datasets (**Fig. 1**). Having near-global and full taxonomic coverage of distribution data (**Fig. 4**) and several functional traits (**Table S 1**), GIFT can help to assess the representativeness of macroecological datasets and to overcome data limitations to find answers to fundamental questions in functional biogeography and macroecology (FitzJohn *et al.*, 2014; e.g. Scheffer *et al.*; König *et al.*, in prep.). It may for example help to estimate whether data from resources like GBIF or TRY are sufficiently complete or representative for analyses of a given taxon, region or functional group (Meyer *et al.*, 2016; König *et al.*, in prep.). Alternatively, GIFT can also be used to infer the floristic status of plant point occurrences (e.g. to tell apart native and non-native species), to identify unlikely or dubious occurrences or to infer local species absences. The latter may be particularly useful for species distribution modelling where random pseudo-absences are commonly used when true absences are not known (Lobo *et al.*, 2010; Barbet-Massin *et al.*, 2012). Furthermore, GIFT can be used to define regional species pools of local plant communities (Karger *et al.*, 2016), for example, for identifying likely source regions of species that colonize oceanic islands (König *et al.*, 2019). Defining the regional species pool or inferring the floristic status may not only be important for macroecological studies but also for field projects at the local to regional scale.

The aggregated nature of data in GIFT, i.e. distribution data at the level of geographic regions and functional traits at the species level allow to achieve taxonomic and geographic coverage which currently is not available in databases of more highly resolved distribution and trait data (e.g. point occurrences and individual level trait measurements) (König *et al.*, in prep.). It is a mid-term goal of GIFT to reach full global coverage of vascular plant checklists. Already now, 79.1 % of the global land surface is covered and further Floras and checklists covering missing parts are currently processed. Realistically, GIFT will reach about 90% spatial coverage in 2019 and will serve as a representative resource for analyses of global plant diversity. In the meantime, regions already covered by coarse geographical units will be complemented by finer-scale data, and new literature resources will be included to update outdated checklists. Once the availability of checklists per region has further increased, workflows to spatially aggregate them will be developed. This will include the identification of conflicting information and choice of the best and most up-to-date information as well as derivation of the floristic status from small to large regions and vice-versa.

A major challenge regards the evaluation of checklist quality and completeness in GIFT (Hortal *et al.*, 2015). The species richness data sets currently being included allow for a comparison of expected and actual species numbers, but also the integration of other data like, for example, point occurrence information as provided by GBIF or vegetation plot data (Bruelheide *et al.*, 2018) may help to estimate completeness of the regional checklists in GIFT and eventually to update them. Furthermore, the lack of cosmopolitan or regionally common species in checklists, an uneven representation of expected higher taxa, or deviances from expected ecological relationships like, for example, the species area relationship or the latitudinal diversity gradient may be used to flag potentially incomplete checklists (Santos *et al.*, 2010). Also conflicting information, like species endemic in one region and native in another (currently 0.39 % of 246,583 species across 2,258 non-overlapping regions) or species not occurring in a region but present in a nested region (currently 5.73 % of 504,389 native species occurrences across 614 regions that are nested in 114 larger regions), can be used to flag potentially incomplete regions. Regions with incomplete checklists can then be excluded from analyses or survey effort can be included in statistical models and data acquisition can be prioritized for those regions (Meyer *et al.*, 2016).

In conclusion, GIFT offers a novel integrated database framework to study the geographic distribution of plant life across the globe. The integration of regional plant checklists with functional traits, phylogenetic relationships and regional environmental characteristics allows for the extraction of well-curated, high quality macroecological datasets for hypothesis testing and the validation and extension (Maitner, 2018)of alternative resources. In addition, the outlined database framework can serve as an example for other taxa with insufficiently complete information at the level of individual species and for an integration of comparable data types such as vegetation plots or surveys. The spatially nested structure of regions in GIFT allows for an ongoing inclusion of resources to improve inventory quality and spatial resolution in future database releases.

### Data accessibility

GIFT integrates various data types from different data domains such as species names, distributions, traits, taxonomy, phylogenies and environmental data. The database structure is relational and highly complex, and data in GIFT may exhibit systematic gaps, biases and uncertainties that users need to account for. Additionally, the data stem from resources with various terms of use. Data from GIFT is therefore currently available upon request. Derived diversity metrics and individual regional checklists without sharing restrictions by the data providers will be be shared without conditions. More complex and larger scale data are available within scientific collaborations. However, we are currently developing data extraction tools (comparable to those in the BIEN R package; Maitner, 2018) that enable users to retrieve data from GIFT and prepare them for macrocological and biogeographical analyses (https://github.com/BioGeoMacro/GIFT-export). Once these tools have reached a status that allows easy handling of the complex content of the database, consideration of data restrictions of the original data sources and evaluation of potential biases, GIFT will become publicly available. Already now, naturalized alien species occurrences from GIFT are included in and accessible via the global naturalized alien flora (GloNAF) database (van Kleunen *et al.*, 2019) and first trait data (e.g. plant growth form) will be published and made publicly available via the online portals of TRY (https://www.try-db.org) and GIFT (http://gift.uni-goettingen.de). The GIFT website also provides an interactive visualization of the geographic coverage of checklist and trait information in GIFT and allows the user to discover data. Here, users can select taxonomic groups and floristic subsets to map aggregated patterns of species richness or percent trait coverage for a given trait across the covered geographic regions. Furthermore, unrestricted species distributions from GIFT will be implemented in the near future for visualization on the Map of Life website (https://mol.org/).

## Acknowledgements

We are grateful to a large group of data contributors who provided either raw and unpublished or digitized Floras and checklists. A full list of all data citations and their bibliographic references is available at the GIFT website (http://gift.uni-goettingen.de). Special thanks go to the GloNAF core team (www.glonaf.org) and Anke Stein for help with finding and digitizing numerous checklists. We thank Martin Turjak for help with developing the database infrastructure and GIFT web-interface and Judith Krobbach, Dagmar Jahn, Lukas Conrad and Julian Schrader for digitizing and preparing checklists for the database import. The GIFT project was founded by HK and initially funded by start-up funding granted to HK via the Free Floater Program in the Excellence Initiative of the University of Göttingen (funded by the German Research Foundation) and is currently financed by core institutional support of the Faculty of Forest Sciences and Forest Ecology at University of Göttingen.

## Author Contributions

PW and HK conceived the GIFT database. All authors led the collection of checklist and trait data. PW and CK developed the workflows for importing and processing data in GIFT and for calculating derived variables. PW and CK performed the analyses presented here and all authors contributed to writing the manuscript.

---

**Supplementary Material**

**Fig. S 1.**
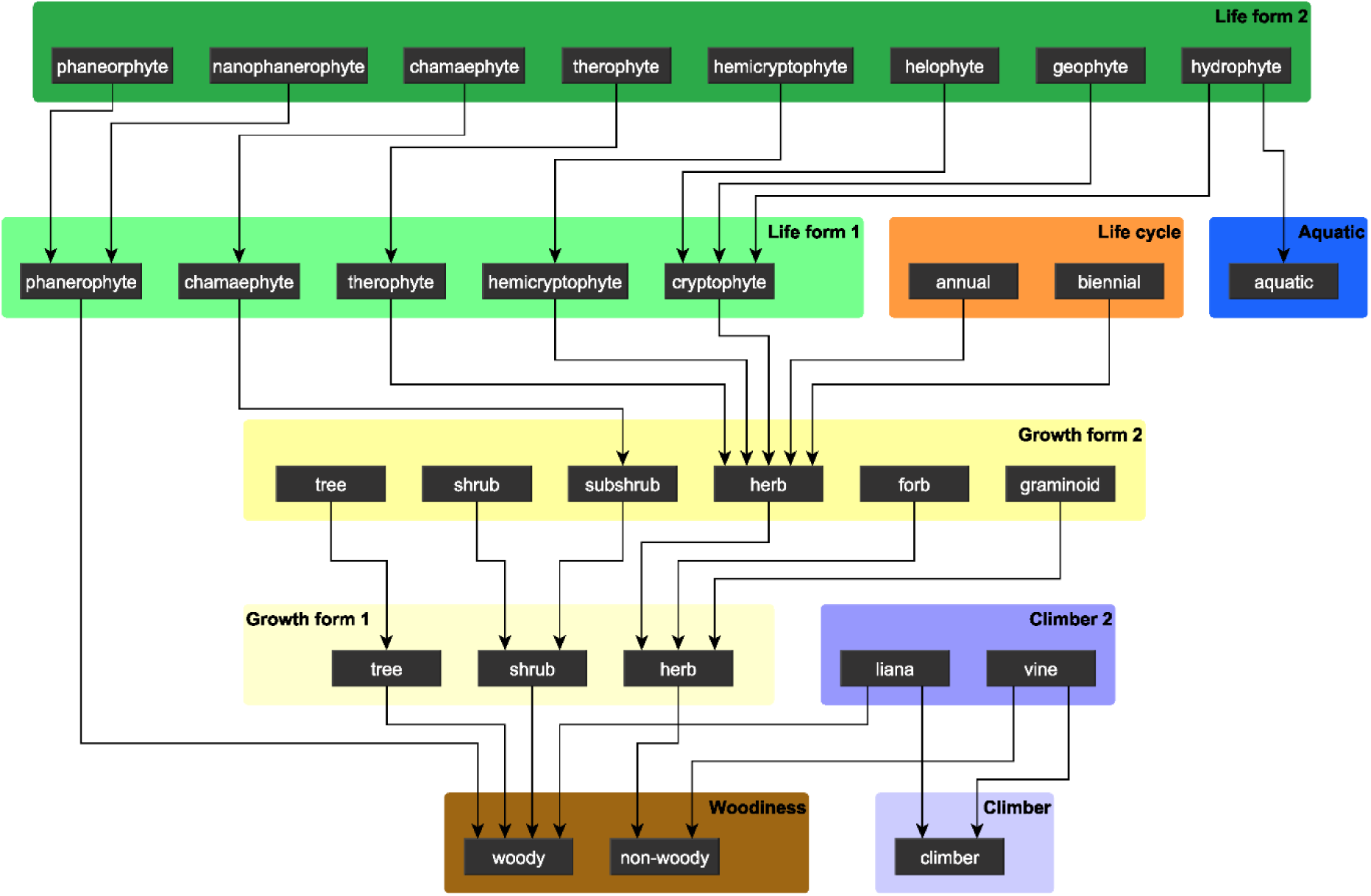
Main module of the directed graph used for hierarchical trait derivation in GIFT, defining unambiguous relationships among 30 categorical levels from five functional traits (life form, life cycle, aquatic, growth form, climber and woodiness). Some traits are represented in multiple versions (e.g. growth form) to account for varying levels of detail of original information. The full list of parent-child relationships used for trait derivation in GIFT (71 connections among 89 categorical levels) is given in **Table S 2.**

**Fig. S 2.**
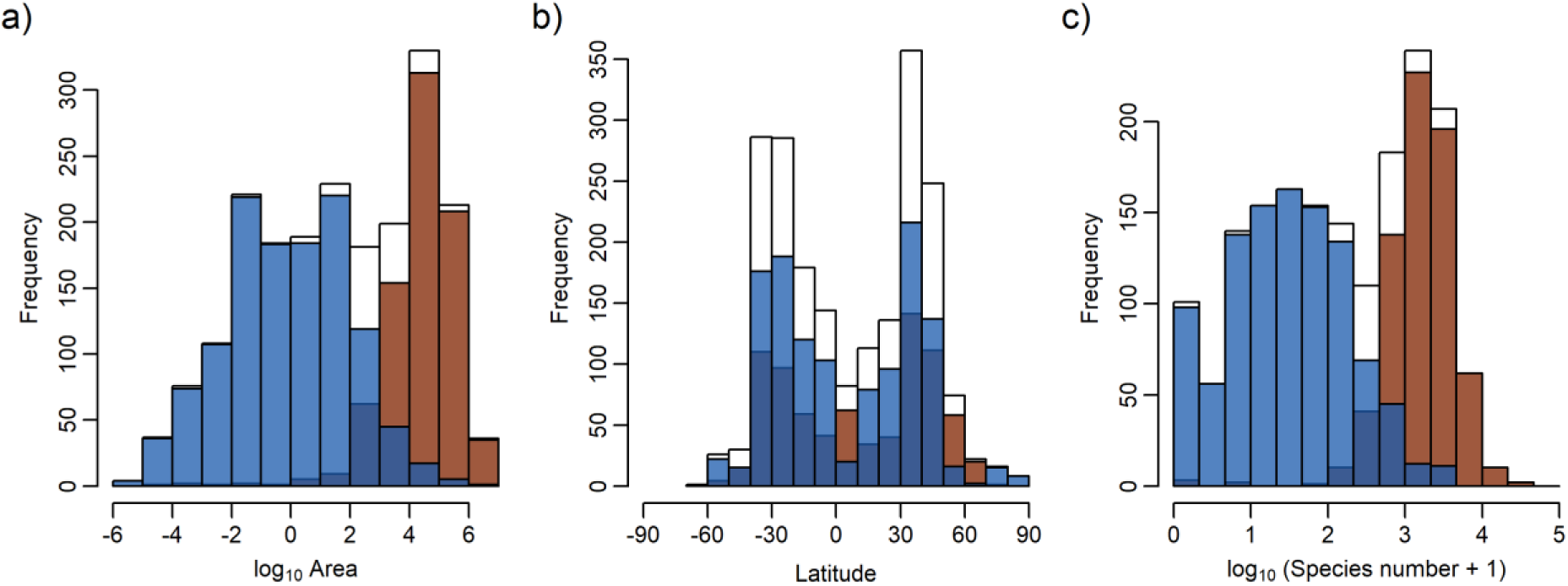
Frequency distributions of 2007 geographic regions in GIFT with information on native vascular plant species composition and spatial properties for a) region area (km²), b) latitude and c) species richness of native vascular plants. Blue bars = islands, brown = mainland regions, white = total, dark blue = overlap of mainland and island bars.

**Fig. S 3.**
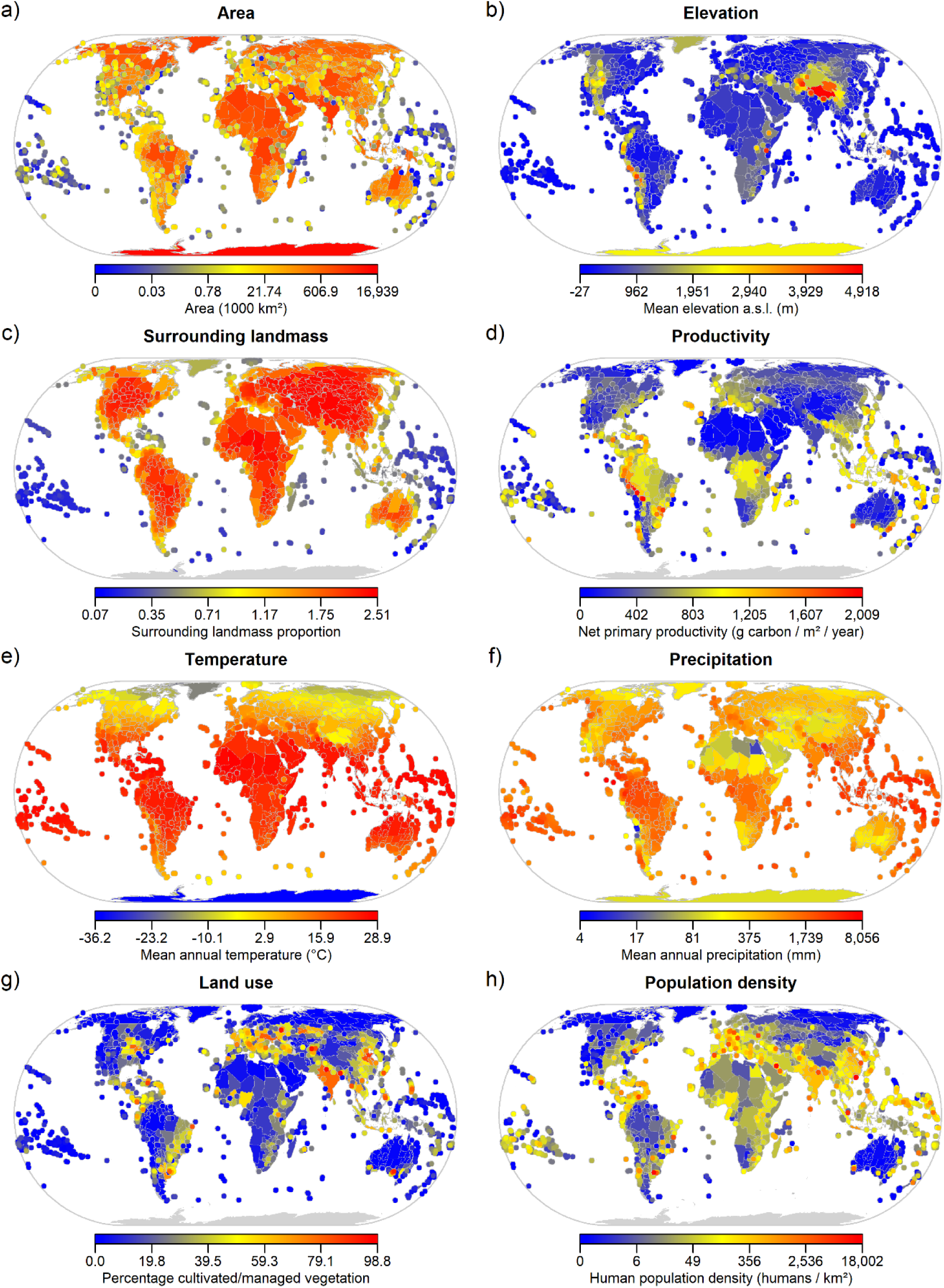
Selected environmental variables in GIFT. For a full list of geographic, environmental and socio-economic variables and source references see **Table S 3**. Polygons are plotted sequentially in order of decreasing area to show smaller regions on top of larger regions, in the case where they overlap. Regions <25,000 km² are plotted as points.

**Fig. S 4.**
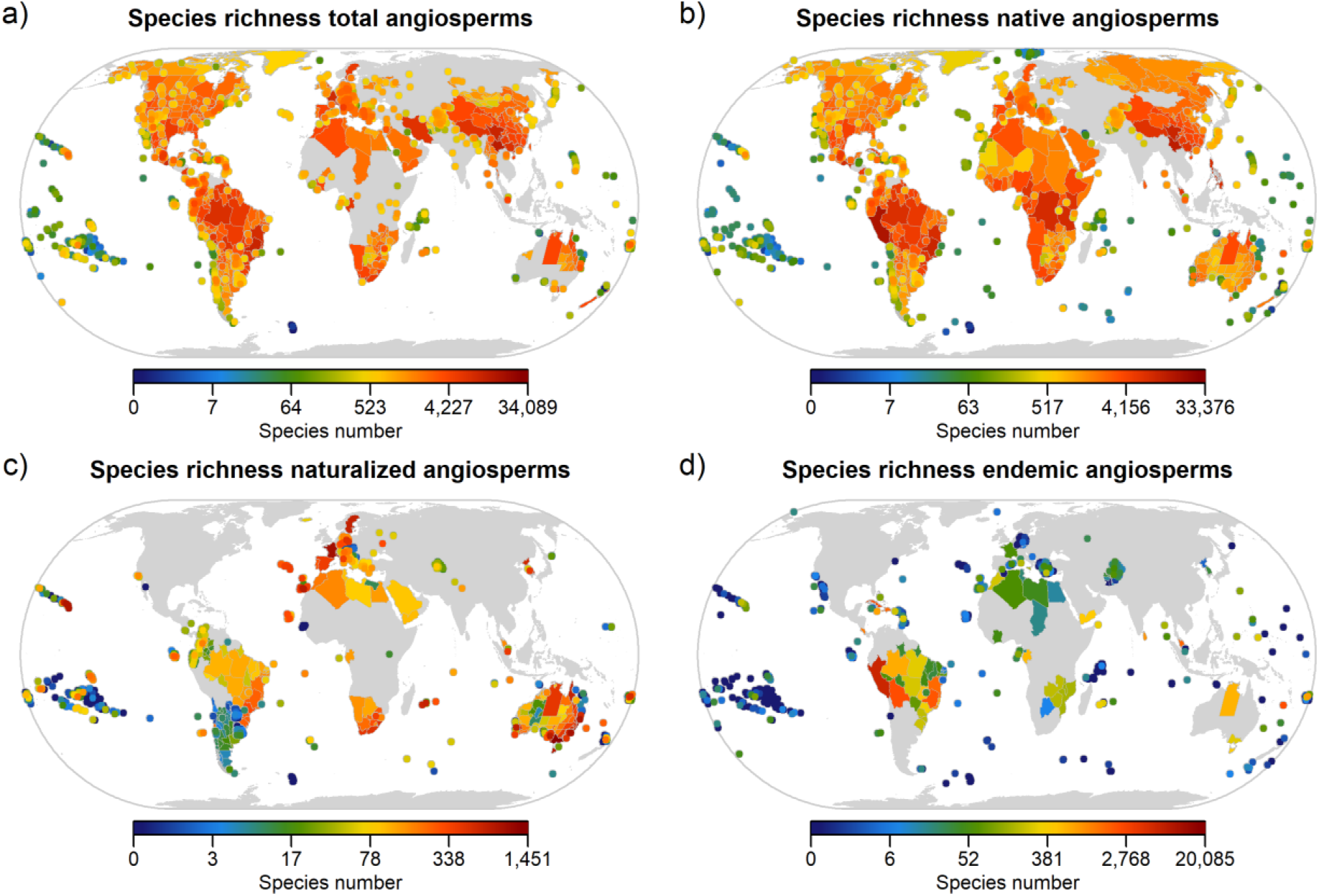
Spatial coverage of floristic subsets in GIFT. Species richness of total angiosperms (a) refers to regions with information on both native and introduced angiosperm species or species with unresolved floristic status. Most commonly resources in GIFT include information on native species (b), while information on introduced naturalized (c) and endemic species (d) is considerably rarer. Endemism information is most common for island regions. Polygons are plotted sequentially in order of decreasing area to show smaller regions on top of larger regions, in the case where they overlap. Regions <25,000 km² are plotted as points.

**Fig. S 5.**
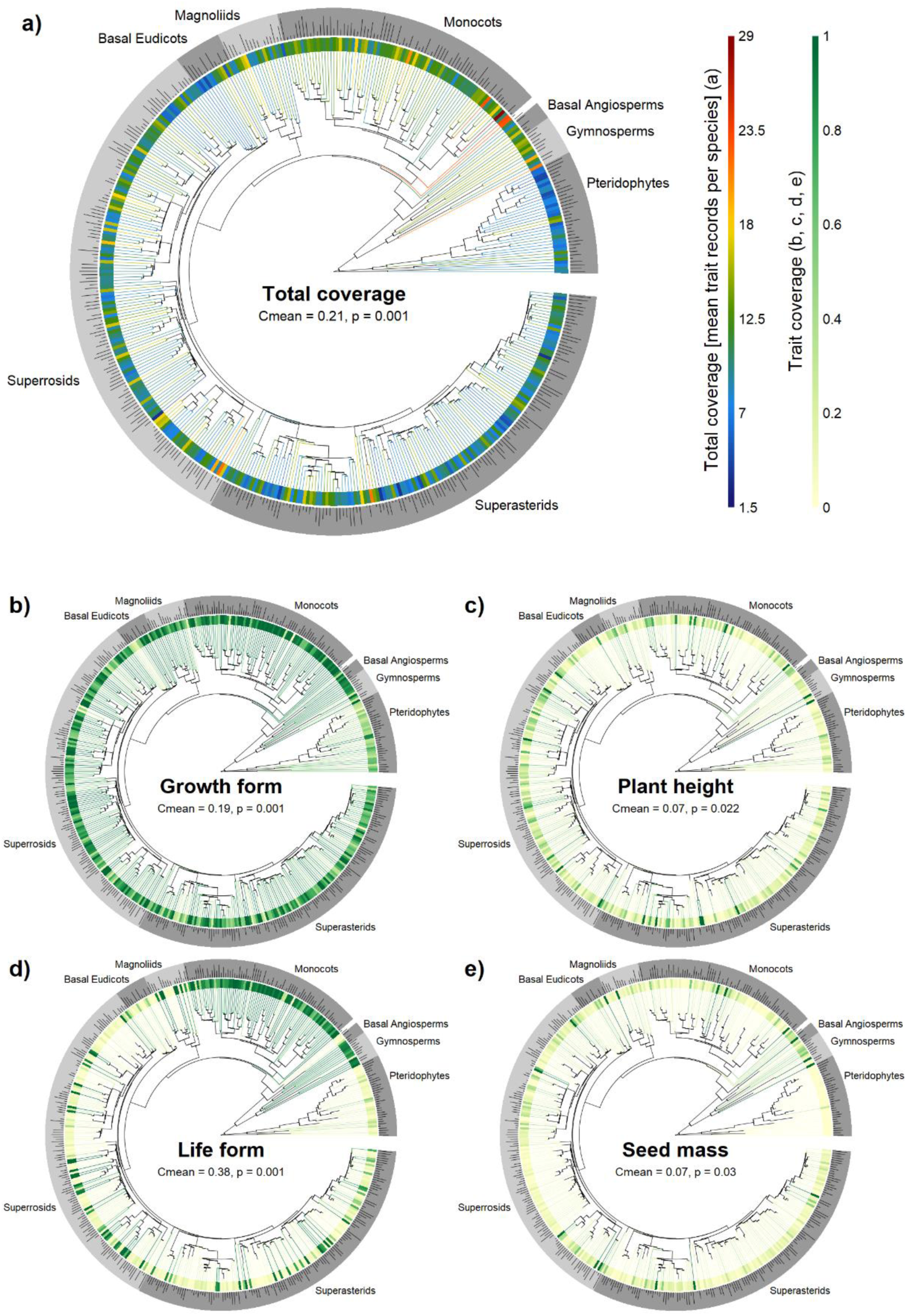
Taxonomic trait coverage of GIFT across all functional traits at the family level (a; mean trait records per species) and for four selected functional traits individually (b-e; number of species with trait information/total number of species). Tip color and inner ring color denote trait coverage per family, outer ring delimits major clades of vascular plants. The height of bars in the outer ring is proportional to log_10_ family size. Phylogenetic signal in taxonomic coverage was assessed as Abouheif’s Cmean, a measure of phylogenetic autocorrelation based on the sum of the successive squared differences between values of neighbouring tips in the phylogeny (Abouheif, 1999).

**Table S 1.**
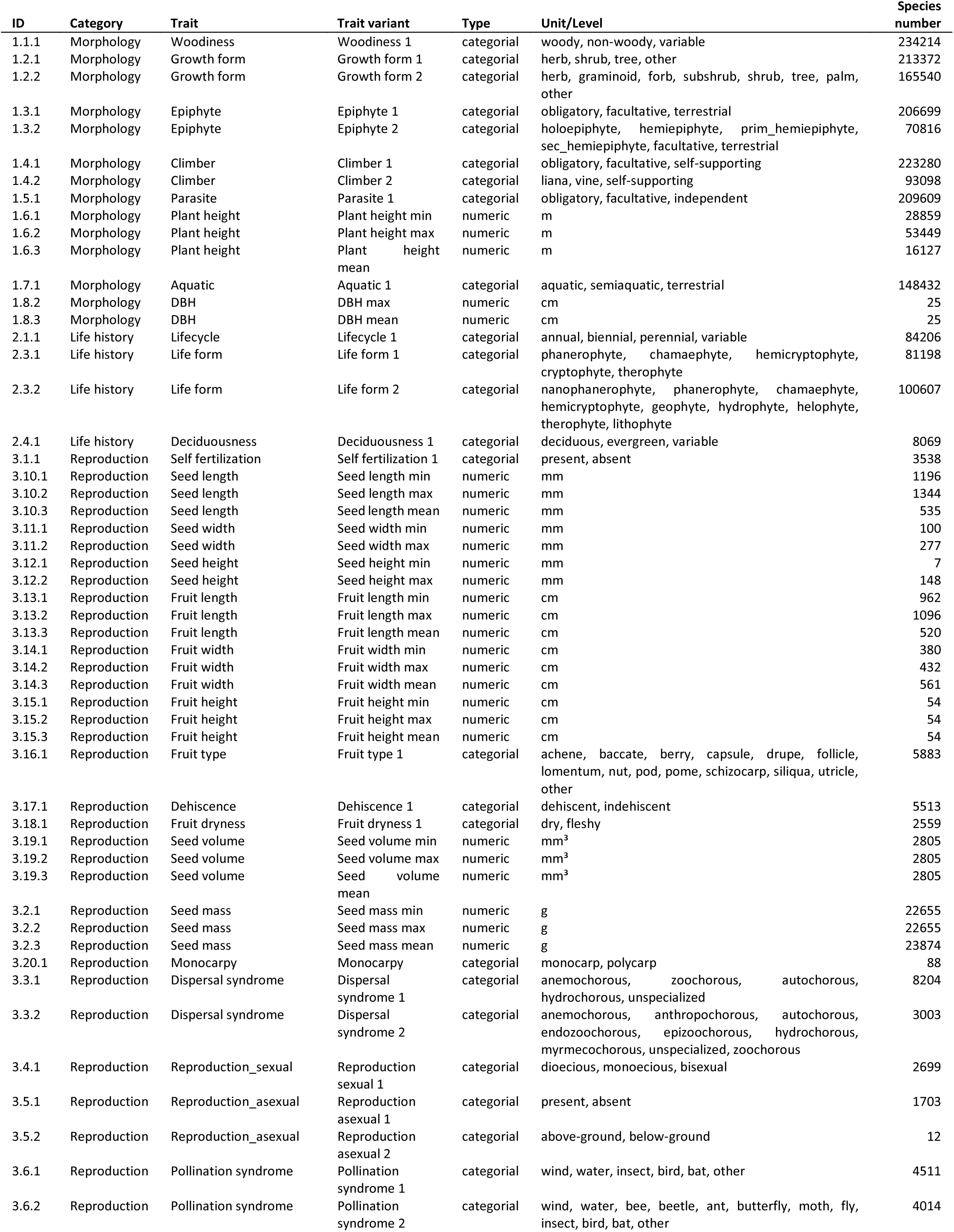
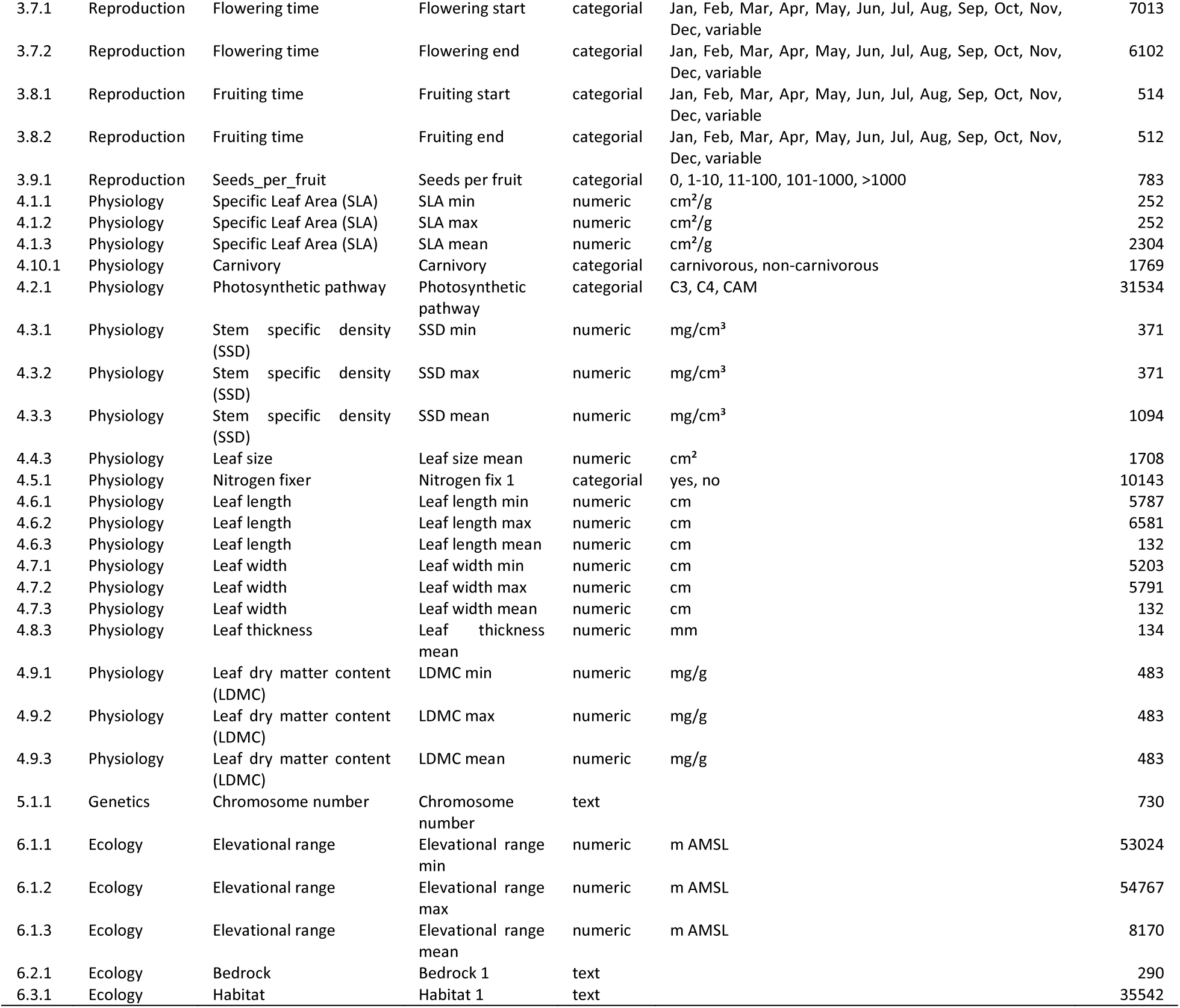
Functional traits in GIFT, their broader trait categories, type, units and factor levels respectively and the number of species covered in GIFT.

**Table S 2.**
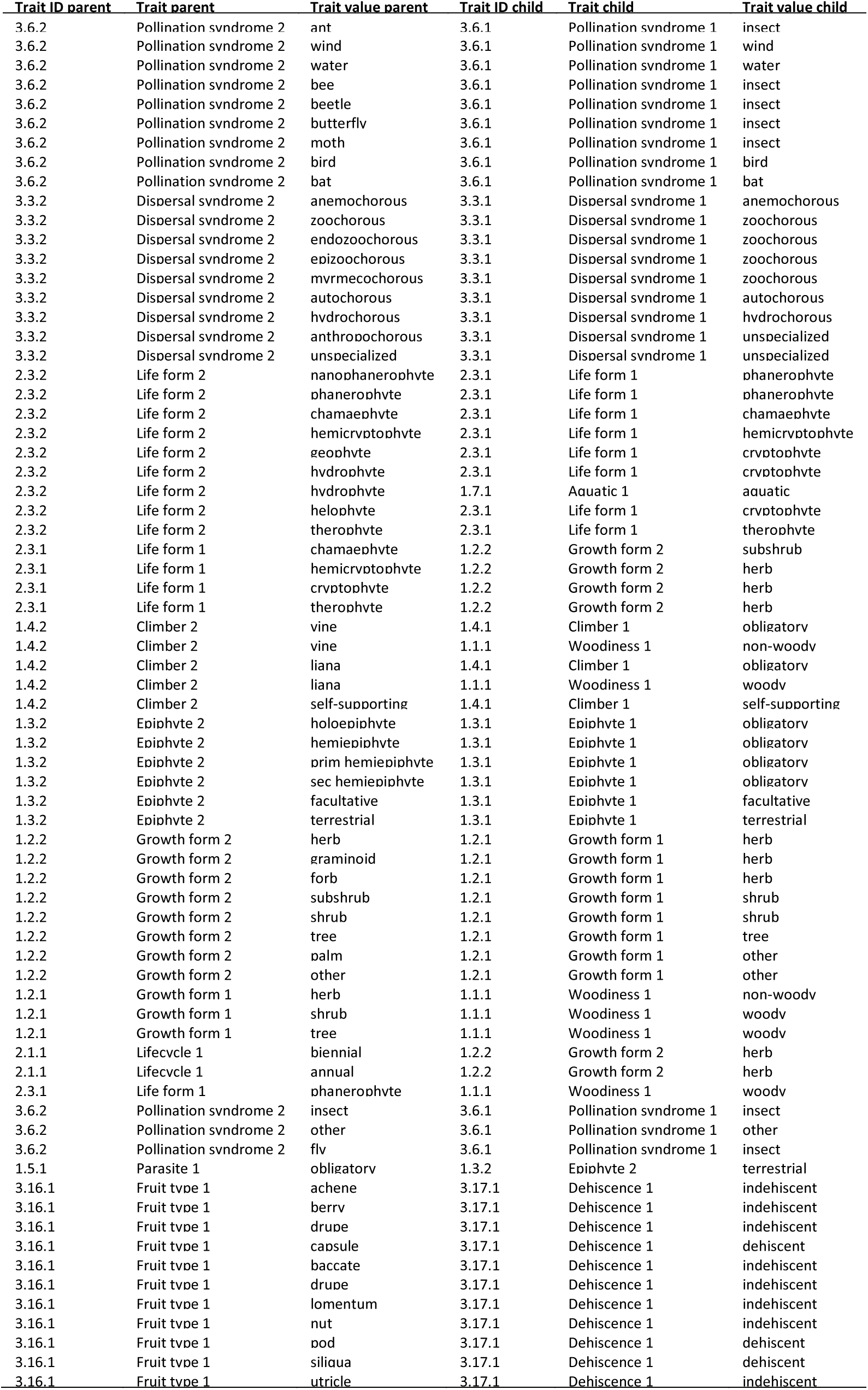
Links between parent traits and derived traits used in the hierarchical trait derivation in GIFT.

**Table S 3.** Groups of physical geographical, environmental and socio-economic variables in GIFT. Metrics of datasets of type miscellaneous are calculated based on the regions polygons and eventually additional resources cited under “References”. For resources of type raster, summary statistics (15 quantiles including minimum, median and maximum, mean, standard deviation, mode, number of unique values, Shannon diversity and number of cells) are calculated for all raster cells that fall into a region or are crossed by its border. Please revisit the original publications of the raster resources for more information.

| Dataset | Type | Reference |
| --- | --- | --- |
| Accessibility to cities | Raster | Weiss *et al.* 2018 |
| Archipelago | Miscellaneous |  |
| Area of ice cover today and during last glacial maximum | Miscellaneous | Ehlers *et al.* 2011 |
| Area of region and largest included landmass | Miscellaneous |  |
| Biome | Miscellaneous | Dinerstein *et al.* 2017 |
| Canopy height | Raster | Simard *et al.* 2011 |
| Centroid latitude and longitude coordinates | Miscellaneous |  |
| Chelsa 1.2 bioclimatic variables | Raster | Karger *et al.* 2017 |
| Consensus land cover | Raster | Tuanmu & Jetz 2014 |
| Distance to nearest mainland | Miscellaneous | Weigelt *et al.* 2013 |
| GDP from satellite | Raster | Ghosh *et al.* 2010 |
| Geographic extent of region and largest included landmass | Miscellaneous |  |
| Global aridity and potential evapotranspiration | Raster | Zomer *et al.* 2008 |
| Global cloud cover | Raster | Wilson & Jetz 2016 |
| Global gridded soil information | Raster | Hengl *et al.* 2017 |
| Global habitat heterogeneity | Raster | Tuanmu & Jetz 2015 |
| Global multi-resolution terrain elevation data | Raster | USGS 2011 |
| Global soil water balance | Raster | Trabucco & Zomer 2010 |
| Gridded population of the world 4 | Raster | Doxsey-Whitfield *et al.* 2015 |
| Human footprint | Raster | Venter *et al.* 2016 |
| Last glacial maximum mainland connection, area and number of landmass entities | Miscellaneous | Weigelt *et al.* 2016, Weatherall *et al.* 2015 |
| Last of the wild human influence | Raster | WCS & CIESIN 2005a, 2005b |
| MODIS gross and net primary productivity | Raster | Zhao & Running 2010 |
| Number of landmass entities comprised by region | Miscellaneous |  |
| Past climate change velocity | Raster | Weigelt *et al.* 2013, Hijmans *et al.* 2005 |
| Species richness at nearest mainland | Miscellaneous | Weigelt *et al.* 2013, Kreft & Jetz 2007 |
| Surrounding landmass proportion | Miscellaneous | Weigelt *et al.* 2013 |
| Takhtajan floristic region | Miscellaneous | Takhtajan 1986 |
| TDWG region | Miscellaneous | Brummitt 2001 |
| Tree density | Raster | Crowther *et al.* 2015 |
| WorldClim 1.4 last glacial maximum climate | Raster | Hijmans *et al.* 2005 |
| WorldClim 2.0 bioclimatic variables | Raster | Fick & Hijmans 2017 |

## References

Abouheif, E. (1999) A method for testing the assumption of phylogenetic independence in comparative data. Evolutionary Ecology Research, 1, 895–909.

Acevedo-Rodríguez, P. & Strong, M.T. (2007) Catalogue of the seed plants of the West Indies Website. Available at: http://botany.si.edu/antilles/WestIndies/index.htm (accessed 5 November 2014).

Barbet-Massin, M., Jiguet, F., Albert, C.H. & Thuiller, W. (2012) Selecting pseudo-absences for species distribution models: how, where and how many? Methods in Ecology and Evolution, 3, 327–338.

Barthlott, W., Mutke, J., Rafiqpoor, D., Kier, G. & Kreft, H. (2005) Global centers of vascular plant diversity. Nova Acta Leopoldina, 92, 61–83.

Barthlott, W., Burstedde, K., Geffert, J.L., Ibisch, P.L., Korotkova, N., Miebach, A., Rafiqpoor, M.D., Stein, A. & Mutke, J. (2015) Biogeography and biodiversity of cacti. Univ.-Verlag Isensee, Germany.

BGCI (2017) GlobalTreeSearch online database. Available at: www.bgci.org/global_tree_search.php (accessed 31.07.2017).

BirdLife International (2018) Taxonomic Checklist of the Birds of the World. Available at: http://www.birdlife.org (accessed 20 February 2018).

Boyle, B., Hopkins, N., Lu, Z., Garay, J.A.R., Mozzherin, D., Rees, T., Matasci, N., Narro, M.L., Piel, W.H. & Mckay, S.J. (2013) The taxonomic name resolution service: an online tool for automated standardization of plant names. BMC Bioinformatics, 14, 16.

Brach, A.R. & Song, H. (2006) eFloras: New directions for online floras exemplified by the Flora of China Project. Taxon, 55, 188–192.

Bruelheide, H., Dengler, J., Purschke, O., Lenoir, J., Jiménez-Alfaro, B., Hennekens, S.M., Botta-Dukát, Z., Chytrý, M., Field, R., Jansen, F., Kattge, J., Pillar, V.D., Schrodt, F., Mahecha, M.D., Peet, R.K., Sandel, B., van Bodegom, P., Altman, J., Alvarez-Dávila, E., Arfin Khan, M.A.S., Attorre, F., Aubin, I., Baraloto, C., Barroso, J.G., Bauters, M., Bergmeier, E., Biurrun, I., Bjorkman, A.D., Blonder, B., Čarni, A., Cayuela, L., Černý, T., Cornelissen, J.H.C., Craven, D., Dainese, M., Derroire, G., De Sanctis, M., Díaz, S., Doležal, J., Farfan-Rios, W., Feldpausch, T.R., Fenton, N.J., Garnier, E., Guerin, G.R., Gutiérrez, A.G., Haider, S., Hattab, T., Henry, G., Hérault, B., Higuchi, P., Hölzel, N., Homeier, J., Jentsch, A., Jürgens, N., Kącki, Z., Karger, D.N., Kessler, M., Kleyer, M., Knollová, I., Korolyuk, A.Y., Kühn, I., Laughlin, D.C., Lens, F., Loos, J., Louault, F., Lyubenova, M.I., Malhi, Y., Marcenò, C., Mencuccini, M., Müller, J.V., Munzinger, J., Myers-Smith, I.H., Neill, D.A., Niinemets, Ü., Orwin, K.H., Ozinga, W.A., Penuelas, J., Pérez-Haase, A., Petřík, P., Phillips, O.L., Pärtel, M., Reich, P.B., Römermann, C., Rodrigues, A.V., Sabatini, F.M., Sardans, J., Schmidt, M., Seidler, G., Silva Espejo, J.E., Silveira, M., Smyth, A., Sporbert, M., Svenning, J.-C., Tang, Z., Thomas, R., Tsiripidis, I., Vassilev, K., Violle, C., Virtanen, R., Weiher, E., Welk, E., Wesche, K., Winter, M., Wirth, C. & Jandt, U. (2018) Global trait–environment relationships of plant communities. Nature Ecology & Evolution, 2, 1906–1917.

Butler, E.E., Datta, A., Flores-Moreno, H., Chen, M., Wythers, K.R., Fazayeli, F., Banerjee, A., Atkin, O.K., Kattge, J., Amiaud, B., Blonder, B., Boenisch, G., Bond-Lamberty, B., Brown, K.A., Byun, C., Campetella, G., Cerabolini, B.E.L., Cornelissen, J.H.C., Craine, J.M., Craven, D., de Vries, F.T., Díaz, S., Domingues, T.F., Forey, E., González-Melo, A., Gross, N., Han, W., Hattingh, W.N., Hickler, T., Jansen, S., Kramer, K., Kraft, N.J.B., Kurokawa, H., Laughlin, D.C., Meir, P., Minden, V., Niinemets, Ü., Onoda, Y., Peñuelas, J., Read, Q., Sack, L., Schamp, B., Soudzilovskaia, N.A., Spasojevic, M.J., Sosinski, E., Thornton, P.E., Valladares, F., van Bodegom, P.M., Williams, M., Wirth, C. & Reich, P.B. (2017) Mapping local and global variability in plant trait distributions. Proceedings of the National Academy of Sciences, 114, E10937–E10946.

Cabral, J.S., Weigelt, P., Kissling, W.D. & Kreft, H. (2014) Biogeographic, climatic and spatial drivers differentially affect α-, β- and γ-diversities on oceanic archipelagos. Proceedings of the Royal Society Biological Sciences Series B, 281, 20133246.

Danielson, J.J. & Gesch, D.B. (2011) Global Multi-resolution Terrain Elevation Data 2010 (GMTED2010). U.S. Geological Survey, Reston, Virginia.

Doxsey-Whitfield, E., MacManus, K., Adamo, S.B., Pistolesi, L., Squires, J., Borkovska, O. & Baptista, S.R. (2015) Taking Advantage of the Improved Availability of Census Data: A First Look at the Gridded Population of the World, Version 4. Papers in Applied Geography, 1, 226–234.

Engemann, K., Sandel, B., Enquist, B.J., Jørgensen, P.M., Kraft, N., Marcuse-Kubitza, A., McGill, B., Morueta-Holme, N., Peet, R.K., Violle, C., Wiser, S. & Svenning, J.-C. (2016) Patterns and drivers of plant functional group dominance across the Western Hemisphere: a macroecological re-assessment based on a massive botanical dataset. Botanical Journal of the Linnean Society, 180, 141–160.

Enquist, B.J., Condit, R., Peet, R.K., Schildhauer, M. & Thiers, B.M. (2016) Cyberinfrastructure for an integrated botanical information network to investigate the ecological impacts of global climate change on plant biodiversity. PeerJ Preprints, 4, e2615v2.

Fang, J., Wang, Z. & Tang, Z. (ed.^eds) (2011) Atlas of woody plants in China: distribution and climate. Springer Science & Business Media, Berlin Heidelberg.

Farjon, A. (2010) A Handbook of the World's Conifers (2 vols.). Brill, Leiden, The Netherlands.

Farjon, A. & Filer, D. (2013) An atlas of the world's conifers: an analysis of their distribution, biogeography, diversity and conservation status. Brill, Leiden, Netherlands.

FitzJohn, R.G., Pennell, M.W., Zanne, A.E., Stevens, P.F., Tank, D.C., Cornwell, W.K. & Shefferson, R. (2014) How much of the world is woody? Journal of Ecology, 102, 1266–1272.

Flann, C. (2009) Global Compositae Checklist. Available at: http://compositae.landcareresearch.co.nz (accessed 3 July 2018).

Frodin, D.G. (2001) Guide to standard floras of the world, 2nd edn. Cambridge University Press, Cambridge, UK.

Garnier, E., Stahl, U., Laporte, M.-A., Kattge, J., Mougenot, I., Kühn, I., Laporte, B., Amiaud, B., Ahrestani, F.S., Bönisch, G., Bunker, D.E., Cornelissen, J.H.C., Díaz, S., Enquist, B.J., Gachet, S., Jaureguiberry, P., Kleyer, M., Lavorel, S., Maicher, L., Pérez-Harguindeguy, N., Poorter, H., Schildhauer, M., Shipley, B., Violle, C., Weiher, E., Wirth, C., Wright, I.J. & Klotz, S. (2017) Towards a thesaurus of plant characteristics: an ecological contribution. Journal of Ecology, 105, 298–309.

GBIF (2018) GBIF: The Global Biodiversity Information Facility. Available at: https://www.gbif.org (accessed 20 February 2018).

Graham, C.H., Carnaval, A.C., Cadena, C.D., Zamudio, K.R., Roberts, T.E., Parra, J.L., McCain, C.M., Bowie, R.C.K., Moritz, C., Baines, S.B., Schneider, C.J., VanDerWal, J., Rahbek, C., Kozak, K.H. & Sanders, N.J. (2014) The origin and maintenance of montane diversity: integrating evolutionary and ecological processes. Ecography, 37, 711–719.

Guralnick, R., Walls, R. & Jetz, W. (2018) Humboldt Core - toward a standardized capture of biological inventories for biodiversity monitoring, modeling and assessment. Ecography, 41, 713–725.

Hijmans, R.J., Garcia, N., Wieczorek, J., Rala, A., Maunahan, A. & Kapoor, J. (2009) GADM database of Global Administrative Areas, Version 1. Available at: https://gadm.org (accessed 16 June 2010).

Hortal, J., Bello, F.d., Diniz-Filho, J.A.F., Lewinsohn, T.M., Lobo, J.M. & Ladle, R.J. (2015) Seven Shortfalls that Beset Large-Scale Knowledge of Biodiversity. Annual Review of Ecology, Evolution, and Systematics, 46, 523–549.

IPNI (2012) The International Plant Names Index. Available at: http://www.ipni.org (accessed 20 February 2018).

IUCN (2018) IUCN Red List of Threatened Species. Available at: http://www.iucnredlist.org (accessed 20 February 2018).

Jansen, F. & Dengler, J. (2010) Plant names in vegetation databases – a neglected source of bias. Journal of Vegetation Science, 21, 1179–1186.

Jardim Botânico do Rio de Janeiro (2016) Flora do Brasil 2020 em construção. Available at: http://floradobrasil.jbrj.gov.br/ (accessed 9 May 2016).

Jetz, W., McPherson, J.M. & Guralnick, R.P. (2012) Integrating biodiversity distribution knowledge: toward a global map of life. Trends in Ecology & Evolution, 27, 151–159.

Karger, D.N., Conrad, O., Böhner, J., Kawohl, T., Kreft, H., Soria-Auza, R.W., Zimmermann, N.E., Linder, H.P. & Kessler, M. (2017) Climatologies at high resolution for the earth’s land surface areas. Scientific Data, 4, 170122.

Karger, D.N., Cord, A.F., Kessler, M., Kreft, H., Kuehn, I., Pompe, S., Sandel, B., Cabral, J.S., Smith, A.B., Svenning, J.C., Tuomisto, H., Weigelt, P. & Wesche, K. (2016) Delineating probabilistic species pools in ecology and biogeography. Global Ecology and Biogeography, 25, 489–501.

Kattge, J., Ogle, K., Bönisch, G., Díaz, S., Lavorel, S., Madin, J., Nadrowski, K., Nöllert, S., Sartor, K. & Wirth, C. (2011a) A generic structure for plant trait databases. Methods in Ecology and Evolution, 2, 202–213.

Kattge, J., Díaz, S., Lavorel, S., Prentice, I.C., Leadley, P., Bönisch, G., Garnier, E., Westoby, M., Reich, P.B., Wright, I.J., Cornelissen, J.H.C., Violle, C., Harrison, S.P., Van Bodegom, P.M., Reichstein, M., Enquist, B.J., Soudzilovskaia, N.A., Ackerly, D.D., Anand, M., Atkin, O., Bahn, M., Baker, T.R., Baldocchi, D., Bekker, R., Blanco, C.C., Blonder, B., Bond, W.J., Bradstock, R., Bunker, D.E., Casanoves, F., Cavender-Bares, J., Chambers, J.Q., Chapin III, F.S., Chave, J., Coomes, D., Cornwell, W.K., Craine, J.M., Dobrin, B.H., Duarte, L., Durka, W., Elser, J., Esser, G., Estiarte, M., Fagan, W.F., Fang, J., Fernández-Méndez, F., Fidelis, A., Finegan, B., Flores, O., Ford, H., Frank, D., Freschet, G.T., Fyllas, N.M., Gallagher, R.V., Green, W.A., Gutierrez, A.G., Hickler, T., Higgins, S.I., Hodgson, J.G., Jalili, A., Jansen, S., Joly, C.A., Kerkhoff, A.J., Kirkup, D., Kitajima, K., Kleyer, M., Klotz, S., Knops, J.M.H., Kramer, K., Kühn, I., Kurokawa, H., Laughlin, D., Lee, T.D., Leishman, M., Lens, F., Lenz, T., Lewis, S.L., Lloyd, J., Llusiá, J., Louault, F., Ma, S., Mahecha, M.D., Manning, P., Massad, T., Medlyn, B.E., Messier, J., Moles, A.T., Müller, S.C., Nadrowski, K., Naeem, S., Niinemets, Ü., Nöllert, S., Nüske, A., Ogaya, R., Oleksyn, J., Onipchenko, V.G., Onoda, Y., Ordoñez, J., Overbeck, G., Ozinga, W.A., Patiño, S., Paula, S., Pausas, J.G., Peñuelas, J., Phillips, O.L., Pillar, V., Poorter, H., Poorter, L., Poschlod, P., Prinzing, A., Proulx, R., Rammig, A., Reinsch, S., Reu, B., Sack, L., Salgado-Negret, B., Sardans, J., Shiodera, S., Shipley, B., Siefert, A., Sosinski, E., Soussana, J.-F., Swaine, E., Swenson, N., Thompson, K., Thornton, P., Waldram, M., Weiher, E., White, M., White, S., Wright, S.J., Yguel, B., Zaehle, S., Zanne, A.E. & Wirth, C. (2011b) TRY – a global database of plant traits. Global Change Biology, 17, 2905–2935.

König, C., Weigelt, P. & Kreft, H. (2017) Dissecting global turnover in vascular plants. Global Ecology and Biogeography, 26, 228–242.

König, C., Weigelt, P., Schrader, J., Taylor, A., Kattge, J. & Kreft, H. (in prep.) Global integration of plant diversity data – the significance of data resolution and domain.

König, C., Weigelt, P., Taylor, A., Stein, A., Dawson, W., Essl, F., Pergl, J., Pysek, P., van Kleunen, M., Winter, M., Chatelain, C., Wieringa, J.J., Krestov, P. & Kreft, H. (2019) Disharmony of the world's island floras. bioRxiv, 523464.

Kreft, H. & Jetz, W. (2007) Global patterns and determinants of vascular plant diversity. Proceedings of the National Academy of Sciences of the United States of America, 104, 5925–5930.

Kreft, H., Jetz, W., Mutke, J., Kier, G. & Barthlott, W. (2008) Global diversity of island floras from a macroecological perspective. Ecology Letters, 11, 116–127.

Lamanna, C., Blonder, B., Violle, C., Kraft, N.J.B., Sandel, B., Šímová, I., Donoghue, J.C., Svenning, J.-C., McGill, B.J., Boyle, B., Buzzard, V., Dolins, S., Jørgensen, P.M., Marcuse-Kubitza, A., Morueta-Holme, N., Peet, R.K., Piel, W.H., Regetz, J., Schildhauer, M., Spencer, N., Thiers, B., Wiser, S.K. & Enquist, B.J. (2014) Functional trait space and the latitudinal diversity gradient. Proceedings of the National Academy of Sciences, 111, 13745–13750.

Lenzner, B., Weigelt, P., Kreft, H., Beierkuhnlein, C. & Steinbauer, M.J. (2017) The general dynamic model of island biogeography revisited at the level of major flowering plant families. Journal of Biogeography, 44, 1029–1040.

Levenshtein, V.I. (1966) Binary codes capable of correcting deletions, insertions, and reversals. Soviet Physics Doklady, 10, 707–710.

Lobo, J.M., Jiménez-Valverde, A. & Hortal, J. (2010) The uncertain nature of absences and their importance in species distribution modelling. Ecography, 33, 103–114.

Mabberley, D.J. (2008) Mabberley's plant-book: a portable dictionary of plants, their classification and uses. Cambridge University Press, Cambridge, UK.

Maitner, B. (2018) BIEN: Tools for Accessing the Botanical Information and Ecology Network Database. R package.

Meyer, C., Weigelt, P. & Kreft, H. (2016) Multidimensional biases, gaps and uncertainties in global plant occurrence information. Ecology Letters, 19, 992–1006.

Moles, A.T., Perkins, S.E., Laffan, S.W., Flores-Moreno, H., Awasthy, M., Tindall, M.L., Sack, L., Pitman, A., Kattge, J., Aarssen, L.W., Anand, M., Bahn, M., Blonder, B., Cavender-Bares, J., Cornelissen, J.H.C., Cornwell, W.K., Díaz, S., Dickie, J.B., Freschet, G.T., Griffiths, J.G., Gutierrez, A.G., Hemmings, F.A., Hickler, T., Hitchcock, T.D., Keighery, M., Kleyer, M., Kurokawa, H., Leishman, M.R., Liu, K., Niinemets, Ü., Onipchenko, V., Onoda, Y., Penuelas, J., Pillar, V.D., Reich, P.B., Shiodera, S., Siefert, A., Sosinski, E.E., Soudzilovskaia, N.A., Swaine, E.K., Swenson, N.G., van Bodegom, P.M., Warman, L., Weiher, E., Wright, I.J., Zhang, H., Zobel, M. & Bonser, S.P. (2014) Which is a better predictor of plant traits: temperature or precipitation? Journal of Vegetation Science, 25, 1167–1180.

Morueta-Holme, N., Enquist, B.J., McGill, B.J., Boyle, B., Jørgensen, P.M., Ott, J.E., Peet, R.K., Šímová, I., Sloat, L.L., Thiers, B., Violle, C., Wiser, S.K., Dolins, S., Donoghue, J.C., Kraft, N.J.B., Regetz, J., Schildhauer, M., Spencer, N. & Svenning, J.-C. (2013) Habitat area and climate stability determine geographical variation in plant species range sizes. Ecology Letters, 16, 1446–54.

Palmer, M.W. & Richardson, J.C. (2012) Biodiversity Data in the Information Age: Do 21st Century Floras Make the Grade? Castanea, 77, 46–59.

Pearse, W.D., Cadotte, M.W., Cavender-Bares, J., Ives, A.R., Tucker, C.M., Walker, S.C. & Helmus, M.R. (2015) pez: phylogenetics for the environmental sciences. Bioinformatics, 31, 2888–2890.

Pearson, R.G. & Dawson, T.P. (2003) Predicting the impacts of climate change on the distribution of species: are bioclimate envelope models useful? Global Ecology and Biogeography, 12, 361–371.

Pérez-Harguindeguy, N., Díaz, S., Garnier, E., Lavorel, S., Poorter, H., Jaureguiberry, P., Bret-Harte, M.S., Cornwell, W.K., Craine, J.M., Gurvich, D.E., Urcelay, C., Veneklaas, E.J., Reich, P.B., Poorter, L., Wright, I.J., Ray, P., Enrico, L., Pausas, J.G., de Vos, A.C., Buchmann, N., Funes, G., Quétier, F., Hodgson, J.G., Thompson, K., Morgan, H.D., ter Steege, H., van der Heijden, M.G.A., Sack, L., Blonder, B., Poschlod, P., Vaieretti, M.V., Conti, G., Staver, A.C., Aquino, S. & Cornelissen, J.H.C. (2013) New handbook for standardised measurement of plant functional traits worldwide. Australian Journal of Botany, 61, 167–234.

Powney, G.D. & Isaac, N.J.B. (2015) Beyond maps: a review of the applications of biological records. Biological Journal of the Linnean Society, 115, 532–542.

Pyšek, P., Pergl, J., Essl, F., Lenzner, B., Dawson, W., Kreft, H., Weigelt, P., Winter, M., Kartesz, J., Nishino, M., Antonova, L.A., Barcelona, J.F., Cabezas, F.J., Cárdenas, D., Cárdenas-Toro, J., Castańo, N., Chacón, E., Chatelain, C., Dullinger, S., Ebel, A.L., Figueiredo, E., Fuentes, N., Genovesi, P., Groom, Q.J., Henderson, L., Inderjit, Kupriyanov, A., Masciadri, S., Maurel, N., Meerman, J., Morozova, O., Moser, D., Nickrent, D., Nowak, P.M., Pagad, S., Patzelt, A., Pelser, P.B., Seebens, H., Shu, W., Thomas, J., Velayos, M., Weber, E., Wieringa, J.J., Baptiste, M.P. & van Kleunen, M. (2017) Naturalized alien flora of the world: species diversity, taxonomic and phylogenetic patterns, geographic distribution and global hotspots of plant invasion. Preslia, 89, 203–274.

Qian, H. & Jin, Y. (2016) An updated megaphylogeny of plants, a tool for generating plant phylogenies and an analysis of phylogenetic community structure. Journal of Plant Ecology, 9, 233–239.

Qian, H., Swenson, N.G. & Zhang, J. (2013) Phylogenetic beta diversity of angiosperms in North America. Global Ecology and Biogeography, 22, 1152–1161.

R Core Team (2018) R: a language and environment for statistical computing. R Foundation for Statistical Computing.

Raunkiær, C.C. (1907) Planterigets livsformer og deres betydning for geografien: Med 77 figurer i teksten. I kommission hos Gyldendalske boghandel, Nordisk forlag.

RBG Kew (2008) Seed Information Database (SID), v 7.1. Available at: http://data.kew.org/sid/ (accessed 3 July 2018).

Reichstein, M., Bahn, M., Mahecha, M.D., Kattge, J. & Baldocchi, D.D. (2014) Linking plant and ecosystem functional biogeography. Proceedings of the National Academy of Sciences, 111, 13697–13702.

Richardson, D.M., Pyšek, P., Rejmánek, M., Barbour, M.G., Panetta, F.D. & West, C.J. (2000) Naturalization and invasion of alien plants: concepts and definitions. Diversity and Distributions, 6, 93–107.

Santos, A.M.C., Jones, O.R., Quicke, D.L.J. & Hortal, J. (2010) Assessing the reliability of biodiversity databases: identifying evenly inventoried island parasitoid faunas (Hymenoptera: Ichneumonoidea) worldwide. Insect Conservation and Diversity, 3, 72–82.

Scheffer, M., Vergnon, R., Cornelissen, J.H.C., Hantson, S., Holmgren, M., van Nes, E.H. & Xu, C. (2014) Why trees and shrubs but rarely trubs? Trends in Ecology & Evolution, 29, 433–434.

Schrodt, F., Kattge, J., Shan, H., Fazayeli, F., Joswig, J., Banerjee, A., Reichstein, M., Bönisch, G., Díaz, S., Dickie, J., Gillison, A., Karpatne, A., Lavorel, S., Leadley, P., Wirth, C.B., Wright, I.J., Wright, S.J. & Reich, P.B. (2015) BHPMF - a hierarchical Bayesian approach to gap-filling and trait prediction for macroecology and functional biogeography. Global Ecology and Biogeography, 24, 1510–1521.

Smith, S.A. & Brown, J.W. (2018) Constructing a broadly inclusive seed plant phylogeny. American Journal of Botany, 105, 302–314.

Stevens, P.F. (2013) Angiosperm Phylogeny Website. Version 13. Available at: http://www.mobot.org/MOBOT/research/APweb/ (accessed 1 May 2013).

Takhtajan, A. (1986) Floristic regions of the world. University of California Press, Berkeley, CA.

TDWG (2007) World geographical scheme for recording plant distributions. Available at: https://github.com/tdwg/prior-standards/tree/master/world-geographical-scheme-for-recording-plant-distributions (accessed 22 December 2017).

The Angiosperm Phylogeny Group (2016) An update of the Angiosperm Phylogeny Group classification for the orders and families of flowering plants: APG IV. Botanical Journal of the Linnean Society, 181, 1–20.

The Plant List (2013) The Plant List, Version 1.1. Available at: http://www.theplantlist.org (accessed 22 December 2017).

Tuomisto, H. (2010) A diversity of beta diversities: straightening up a concept gone awry. Part 1. Defining beta diversity as a function of alpha and gamma diversity. Ecography, 33, 2–22.

Tutin, T., Heywood, V., Burges, N., Moore, D., Valentine, D., Walters, S. & Webb, D. (ed.^eds) (1964–1980) Flora Europaea. Volume 1-5. Cambridge University Press, Cambridge, UK.

Ulloa Ulloa, C., Acevedo-Rodríguez, P., Beck, S., Belgrano, M.J., Bernal, R., Berry, P.E., Brako, L., Celis, M., Davidse, G., Forzza, R.C., Gradstein, S.R., Hokche, O., León, B., León-Yánez, S., Magill, R.E., Neill, D.A., Nee, M., Raven, P.H., Stimmel, H., Strong, M.T., Villaseñor, J.L., Zarucchi, J.L., Zuloaga, F.O. & Jørgensen, P.M. (2017) An integrated assessment of the vascular plant species of the Americas. Science, 358, 1614–1617.

UNEP-WCMC (2013) Global distribution of islands. Global Island Database (version 2). Based on Open Street Map data (© OpenStreetMap contributors). Cambridge, UK.

UNEP-WCMC (2014) Data Standards for the World Database on Protected Areas (WDPA). Cambridge, UK.

van Kleunen, M., Dawson, W., Essl, F., Pergl, J., Winter, M., Weber, E., Kreft, H., Weigelt, P., Kartesz, J., Nishino, M., Antonova, L.A., Barcelona, J.F., Cabezas, F.J., Cardenas, D., Cardenas-Toro, J., Castano, N., Chacon, E., Chatelain, C., Ebel, A.L., Figueiredo, E., Fuentes, N., Groom, Q.J., Henderson, L., Inderjit, Kupriyanov, A., Masciadri, S., Meerman, J., Morozova, O., Moser, D., Nickrent, D.L., Patzelt, A., Pelser, P.B., Baptiste, M.P., Poopath, M., Schulze, M., Seebens, H., Shu, W.S., Thomas, J., Velayos, M., Wieringa, J.J. & Pysek, P. (2015) Global exchange and accumulation of non-native plants. Nature, 525, 100–103.

van Kleunen, M., Pysek, P., Dawson, W., Essl, F., Kreft, H., Pergl, J., Weigelt, P., Stein, A., Dullinger, S., Konig, C., Lenzner, B., Maurel, N., Moser, D., Seebens, H., Kartesz, J., Nishino, M., Aleksanyan, A., Ansong, M., Antonova, L.A., Barcelona, J.F., Breckle, S.W., Brundu, G., Cabezas, F.J., Cardenas, D., Cardenas-Toro, J., Castano, N., Chacon, E., Chatelain, C., Conn, B., de Sa Dechoum, M., Dufour-Dror, J.M., Ebel, A.L., Figueiredo, E., Fragman-Sapir, O., Fuentes, N., Groom, Q.J., Henderson, L., Inderjit, Jogan, N., Krestov, P., Kupriyanov, A., Masciadri, S., Meerman, J., Morozova, O., Nickrent, D., Nowak, A., Patzelt, A., Pelser, P.B., Shu, W.S., Thomas, J., Uludag, A., Velayos, M., Verkhosina, A., Villasenor, J.L., Weber, E., Wieringa, J.J., Yazlik, A., Zeddam, A., Zykova, E. & Winter, M. (2019) The Global Naturalized Alien Flora (GloNAF) database. Ecology, 100, e02542.

Violle, C., Reich, P.B., Pacala, S.W., Enquist, B.J. & Kattge, J. (2014) The emergence and promise of functional biogeography. Proceedings of the National Academy of Sciences, 111, 13690–13696.

WCSP (2012) World Checklist of Selected Plant Families. Available at: http://apps.kew.org/wcsp/ (accessed 18 December 2014).

Weigelt, P. (2015) The macroecology of island floras. Frontiers of Biogeography, 7, fb_25073.

Weigelt, P. & Kreft, H. (2013) Quantifying island isolation – insights from global patterns of insular plant species richness. Ecography, 36, 417–429.

Weigelt, P., Jetz, W. & Kreft, H. (2013) Bioclimatic and physical characterization of the world’s islands. Proceedings of the National Academy of Sciences of the United States of America, 110, 15307–15312.

Weigelt, P., Steinbauer, M.J., Cabral, J.S. & Kreft, H. (2016) Late Quaternary climate change shapes island biodiversity. Nature, 532, 99–102.

Weigelt, P., Kissling, W.D., Kisel, Y., Fritz, S.A., Karger, D.N., Kessler, M., Lehtonen, S., Svenning, J.-C. & Kreft, H. (2015) Global patterns and drivers of phylogenetic structure in island floras. Scientific Reports, 5, 12213.

Wieczorek, J., Bloom, D., Guralnick, R., Blum, S., Döring, M., Giovanni, R., Robertson, T. & Vieglais, D. (2012) Darwin Core: An Evolving Community-Developed Biodiversity Data Standard. PLOS ONE, 7, e29715.

Wieczorek, J., Bánki, O., Blum, S., Deck, J., Döring, M., Dröge, G., Endresen, D., Goldstein, P., Leary, P., Krishtalka, L., Tuama, É.Ó., Robbins, R.J., Robertson, T. & Yilmaz, P. (2014) Meeting Report: GBIF hackathon-workshop on Darwin Core and sample data (22-24 May 2013). Standards in Genomic Sciences, 9, 585–598.

Willis, K.J. (2017) The state of the world’s plants 2017. Report. Royal Botanic Gardens, Kew, UK.

Winter, M., Schweiger, O., Klotz, S., Nentwig, W., Andriopoulos, P., Arianoutsou, M., Basnou, C., Delipetrou, P., Didžiulis, V., Hejda, M., Hulme, P.E., Lambdon, P.W., Pergl, J., Pyšek, P., Roy, D.B. & Kühn, I. (2009) Plant extinctions and introductions lead to phylogenetic and taxonomic homogenization of the European flora. Proceedings of the National Academy of Sciences, 106, 21721–21725.

Zotz, G. (2013) The systematic distribution of vascular epiphytes – a critical update. Botanical Journal of the Linnean Society, 171, 453–481.

Zuloaga, F., Morrone, O. & Belgrano, M.J. (2004) Catálogo de las Plantas Vasculares del Cono Sur. Available at: http://www.darwin.edu.ar/Proyectos/FloraArgentina/fa.htm (accessed 16.03.2015).

## References

Brummitt, R. K. (2001) World geographical scheme for recording plant distributions. Plant taxonomic database standards No. 2. (with assistance from F. Pando, S. Hollis, N. A. Brummitt and others). Edition 2. Hunt Institute for Botanical Documentation, Carnegie Mellon University, Pittsburgh, US.

Crowther, T.W., Glick, H.B., Covey, K.R., Bettigole, C., Maynard, D.S., Thomas, S.M., et al. (2015) Mapping tree density at a global scale. Nature 525, 201–205.

Dinerstein, E., Olson, D., Joshi, A., Vynne, C., Burgess, N.D., Wikramanayake, E., et al. (2017) An ecoregion-based approach to protecting half the terrestrial realm. BioScience 67, 534–545.

Doxsey-Whitfield, E., MacManus, K., Adamo, S.B., Pistolesi, L., Squires, J., Borkovska, O. & Baptista, S.R. (2015) Taking advantage of the improved availability of census data: A first look at the gridded population of the world, version 4. Papers in Applied Geography 1, 226–234.

Ehlers, J., Gibbard, P.L. & Hughes, P.D. (2011) Quaternary glaciations - extent and chronology. A closer look. Elsevier, Amsterdam, The Netherlands.

Fick, S.E. & Hijmans, R.J. (2017) WorldClim 2: new 1-km spatial resolution climate surfaces for global land areas. International Journal of Climatology 37, 4302–4315.

Ghosh, T., Powell, R.L., Elvidge, C.D., Baugh, K.E., Sutton, P.C. & Anderson, S. (2010) Shedding light on the global distribution of economic activity. The Open Geography Journal 3, 147–160.

Hengl, T., Mendes de Jesus, J., Heuvelink, G.B.M., Ruiperez Gonzalez, M., Kilibarda, M., Blagotić, A., et al. (2017) SoilGrids250m. Global gridded soil information based on machine learning. PloS one 12, e0169748. DOI: 10.1371/journal.pone.0169748.

Hijmans, R.J., Cameron, S.E., Parra, J.L., Jones, P.G. & Jarvis, A. (2005) Very high resolution interpolated climate surfaces for global land areas. International Journal of Climatology 25, 1965–1978.

Karger, D. Nikolaus, Conrad, O., Böhner, J., Kawohl, T., Kreft, H., Soria-Auza, R.W., et al. (2017) Climatologies at high resolution for the earth’s land surface areas. Scientific Data 4, 170122. DOI: 10.1038/sdata.2017.122.

Kreft, H. & Jetz, W. (2007) Global patterns and determinants of vascular plant diversity. Proceedings of the National Academy of Sciences 104, 5925–5930.

Simard, M., Pinto, N., Fisher, J.B. & Baccini, A. (2011) Mapping forest canopy height globally with spaceborne lidar. Journal of Geophysical Research 116. G04021. DOI: 10.1029/2011JG001708.

Takhtajan, A. (1986) Floristic regions of the world. University of California Press, Berkeley, US.

Trabucco, A. & Zomer, R. J. (2010) Global Soil Water Balance Geospatial. CGIAR Consortium for Spatial Information. Available at: http://www.cgiar-csi.org/data/global-high-resolution-soil-water-balance (accessed 18 April 2016).

Tuanmu, M.-N. & Jetz, W. (2014) A global 1-km consensus land-cover product for biodiversity and ecosystem modelling. Global Ecology and Biogeography 23, 1031–1045.

Tuanmu, M.-N. & Jetz, W. (2015) A global, remote sensing-based characterization of terrestrial habitat heterogeneity for biodiversity and ecosystem modelling. Global Ecology and Biogeography 24, 1329–1339.

USGS (2011) Global Multi-resolution Terrain Elevation Data 2010 (GMTED2010). U.S. Geological Survey. Virginia. Available at https://lta.cr.usgs.gov/GMTED2010 (accessed 2 May 2017).

Venter, O., Sanderson, E.W., Magrach, A., Allan, J.R., Beher, J., Jones, K., et al. (2016) Global terrestrial Human Footprint maps for 1993) and 2009. Scientific Data 3, 160067. DOI: 10.1038/sdata.2016.67.

WCS & CIESIN (2005a) Last of the Wild Project, Version 2, 2005 (LWP-2). Global Human Footprint Dataset (Geographic). NASA Socioeconomic Data and Applications Center (SEDAC), Palisades, US. DOI: 10.7927/H4M61H5F.

WCS & CIESIN (2005b) Last of the Wild Project, Version 2, 2005 (LWP-2). Global Human Influence Index (HII) Dataset (Geographic). NASA Socioeconomic Data and Applications Center (SEDAC), Palisades, US. DOI: 10.7927/H4BP00QC.

Weatherall, P., Marks, K.M., Jakobsson, M., Schmitt, T., Tani, S., Arndt, J.E., et al. (2015) A new digital bathymetric model of the world’s oceans. Earth and Space Science 2, 331–345. DOI: 10.1002/2015EA000107.

Weigelt, P., Jetz, W. & Kreft, H. (2013) Bioclimatic and physical characterization of the world’s islands. Proceedings of the National Academy of Sciences 110, 15307–15312. DOI: 10.1073/pnas.1306309110.

Weigelt, P., Steinbauer, M.J., Cabral, J.S. & Kreft, H. (2016) Late Quaternary climate change shapes island biodiversity. Nature 532, 99–102.

Weiss, D.J., Nelson, A., Gibson, H.S., Temperley, W., Peedell, S., Lieber, A., et al. (2018) A global map of travel time to cities to assess inequalities in accessibility in 2015. Nature 553, 333. DOI: 10.1038/nature25181.

Wilson, A.M. & Jetz, W. (2016) Remotely sensed high-resolution global cloud dynamics for predicting ecosystem and biodiversity distributions. PLOS Biology 14, e1002415. DOI: 10.1371/journal.pbio.1002415.

Zhao, M. & Running, S.W. (2010) Drought-induced reduction in global terrestrial net primary production from 2000) through 2009. Science 329, 940–943.

Zomer, R.J., Trabucco, A., Bossio, D.A. & Verchot, L.V. (2008) Climate change mitigation. A spatial analysis of global land suitability for clean development mechanism afforestation and reforestation. Agriculture, Ecosystems & Environment 126, 67–80.

## Appendix S 1: Literature resources used to assemble the species checklist and trait data in GIFT

3D Environmental (2008) Vegetation Communities and Regional Ecosystems of The Torres Strait Islands, Queensland, Australia.

3D Environmental (2013a) Profile for management of the habitats and related ecological and cultural resource values of Badu Island.

3D Environmental (2013b) Profile for management of the habitats and related ecological and cultural resource values of Boigu Island.

3D Environmental (2013c) Profile for management of the habitats and related ecological and cultural resource values of Dauan Island.

3D Environmental (2013d) Profile for management of the habitats and related ecological and cultural resource values of Saibai Island.

3D Environmental (2013e) Profile for management of the habitats and related ecological and cultural resource values of Warraber Island.

Abbadi, G.A. & El-Sheikh, M.A. (2002) Vegetation analysis of Failaka Island (Kuwait). Journal of Arid Environments, 50, 153–165.

Abbott, I. & Black, R. (1980) Changes in species composition of floras on islets near Perth, Western Australia. Journal of Biogeography, 7, 399–410.

Abdel Khalik, K., El-Sheikh, M. & El-Aidarous, A. (2013) Floristic diversity and vegetation analysis of Wadi Al-Noman, Mecca, Saudi Arabia. Turkish Journal of Botany, 37, 894–907.

Abderrahman, A. & El Kadmiri, A.A. (2005) Richesse et diversité floristique de la subéraie de la Mamora (Maroc). Acta Botanica Malacitana, 30, 127–138.

Abe, T. (2006) Threatened pollination systems in native flora of the Ogasawara (Bonin) Islands. Annals of Botany (London), 98, 317.

Academica Sinica (ed) (1998) Proc Int Symp on Rare, Threatened, and Endangered Floras of Asia and the Pacific. Institute of Botany, Taiwan.

Acevedo-Rodríguez, P. & Strong, M.T. (2007) Catalogue of the seed plants of the West Indies Website. http://botany.si.edu/antilles/WestIndies/catalog.htm Accessed 1 March 11.

Adou Yao, C.Y. (2005) Diversité floristique et végétation dans le Parc National de Taï, Côte d'Ivoire. Tropenbos Côte d'Ivoire, Abidjan.

Al Khulaidi, A.W.A. Flora of Yemen. Sustainable Environmental Management Program, Republic of Yemen.

Al-Abbadi, G.A. & El-Sheikh, M.A.E. (2017) Vegetation ecology and diversity of six Kuwait Islands: Factors influence on species composition and richness. Rendiconti Lincei, 28 (1), 117–131.

Al-Eisawi, D.M. & Al-Khader, I.A. (2007) Checklist of Plants of the Hashemite Kingdom of Jordan. http://ww2.odu.edu/~lmusselm/plant/jordan/ Accessed 12 January 15.

Alves, R.J.V. (1998) Ilha da Trindade & Arquipélago Martin Vaz: Um ensaio geobotânico. Serviço de Documentação da Marinha, Rio de Janeiro, Brasil.

Alves, Ruy, José Válka & Kolbek, J. (2009) Summit vascular flora of Serra de São José, Minas Gerais, Brazil. Check list, 5 (1), 35–73.

Amerson Jr, A.B. (1975) Species Richness on the nondisturbed Northwestern Hawaiian Islands. Ecology, 56, 435–444.

Arakaki, M. & Cano, A. (2003) Composición florística de la cuenca del río Ilo-Moquegua y Lomas de Ilo, Moquegua, Peru. Revista Peruana de Biología, 10, 5–19.

Arechavaleta, M., Rodríguez, S., Zurita, N. & García, A. (2009) Lista de especies silvestres de Canarias. Hongos, plantas y animales terrestres. Consejería de Medio Ambiente y Ordenación Territorial, Gobierno de Canarias, Santa Cruz de Tenerife, Spain.

Arechavaleta, M., Zurita, N., Marrero, M.C. & Martín, J.L. (2005) Lista preliminar de especies silvestres de Cabo Verde (hongos, plantas y animales terrestres). Consejería de Medio Ambiente y Ordenación Territorial, Gobierno de Canarias, Santa Cruz de Tenerife, Spain.

Ashmole, P. & Ashmole, M. (2000) St Helena and Ascension Island: A natural history. Anthony Nelson Ltd, Oswestry, Shropshire, UK.

Assédé, E.P.S., Adomou, A.C. & Sinsin, B. (2012) Magnoliophyta, Biosphere Reserve of Pendjari, Atacora Province, Benin. Check list, 8, 642–661.

Assyov, B., Petrova, A., Dimitrov, D. & Vassilev, R. (2012) Conspectus of the Bulgarian vascular flora, Sofia.

Athens, J.S., Blinn, D.W. & Ward, J.V. (2007) Vegetation history of Laysan Island, Northwestern Hawaiian Islands. Pacific Science, 61, 17–37.

Atkinson, I.A.E. (1972) Vegetation and flora of Sail Rock, Hen and Chickens Islands. New Zealand journal of botany, 10, 545–558.

Azbukina, Z.M., Bardunov, L.V., Bezdeleva, T.A., Bogacheva, A.V., Bulakh, E.M., Vasilyeva, L.N., Govorova, O.K., Egorova, L.N., Zhabyko, E.V., Nikulina, T.V., Rodnikova, I.M., Skirina, I.F., Tarankov, V.I., Fedina, L.A. & Cherdantseva, V.Y. (2006) Flora, vegetation and mycobiota of the reserve «Ussuriysky». Dalnauka, Vladivostok.

Badshah, L., Hussain, F. & Sher, Z. (2013) Floristic inventory, ecological characteristics and biological spectrum of rangeland, District Tank, Pakistan. Pak. J. Bot, 45 (4), 1159–1168.

Baker, M.L. & Duretto, M.F. (2011) A census of the vascular plants of Tasmania. Tasmanian Herbarium, Tasmanian Museum and Art Gallery, Hobart, Australia.

Bakis, Y., Babac, M.T. & Uslu, E. (2015) Tübives: Turkish Plants Data Service. http://www.tubives.com/ Accessed 13 February 15.

Bancheva, S. & Vassilev, K. (2006) Vascular flora of the Beli Lom Nature Reserve in Northeast Bulgaria. Phytologia Balcanica, 12, 377–386.

Barker, W.R., Barker, R.M., Jessop, J.P. & Vonow, H.P. (2005) Census of South Australian vascular plants. Journal of the Adelaide Botanic Gardens Supplement, 1, 1–396.

Batalha, M.A. & Mantovani, W. (2001) Floristic composition of the Cerrado in the Pé-De-Gigante Reserve (Santa Rita do Passa Quatro, Southeastern Brazil). Acta botanica Brasilica, 15, 289–304.

Batalha, M.A. & Martins, F.R. (2004) Reproductive phenology of the cerrado plant community in Emas National Park (central Brazil). Australian Journal of Botany, 52, 149–161.

Batianoff, G.N., Naylor, G.C., Dillewaard, H.A. & Neldner, V.J. (2009) Plant strategies, dispersal and origins of flora at the northern Coral Sea Islands Territory, Australia. Cunninghamia, 11, 97–106.

Batianoff, G.N., Neldner, V.J., Naylor, G.C. & Olds, J.A. (2012) Mapping and evaluating Capricornia Cays vegetation and regional ecosystems.

Belhacene, L. (2010) Catalogue 2010 des plantes vasculaires du département de la Haute-Garonne. Supplément à Isaatis, 10, 1–145.

Benito, B.M., Lorite, J., Pérez-Pérez, R., Gómez-Aparicio, L., Peñas, J. & Robertson, M. (2014) Forecasting plant range collapse in a mediterranean hotspot: When dispersal uncertainties matter. Diversity and Distributions, 20, 72–83.

Bernal, R., Gradstein, S.R. & Celis, M. (2015) Catálogo de plantas y líquenes de Colombia. http://catalogoplantasdecolombia.unal.edu.co/ accessed 15 January 16.

BGCI (2017) GlobalTreeSearch online database. www.bgci.org/globaltree_search.php Accessed 14 August 17.

Bingham, M.G., Willemen, A., Wursten, B.T., Ballings, P. & Hyde, M.A. (2016) Flora of Zambia. http://www.zambiaflora.com/ Accessed 14 November 16.

BioScripts (2014) Flora Vascular. http://www.floravascular.com/ Accessed 25 May 14.

Borges, P.A.V., Abreu, C., Aguiar, A.M.F., Carvalho, P., Jardim, R., Melo, I., Oliveira, P., Sérgio, C., Serrano, A.R.M. & Vieira, P. (2008) Listagem dos fungos, flora e fauna terrestres dos arquipélagos da Madeira e Selvagens. Direcção Regional do Ambiente da Madeira and Universidade dos Açores, Funchal and Angra do Heroísmo, Portugal.

Borges, P.A.V., Costa, A., Cunha, R., Gabriel, R., Gonçalves, V., Martins, A.F., Melo, I., Parente, M., Raposeiro, P., Rodrigues, P., Santos, R.S., Silva, L., Vieira, P. & Vieira, V. (2010) A list of the terrestrial and marine biota from the Azores. Princípia, Cascais.

Botanical Garden Tel Aviv Israel Flora. Tel Aviv University.

Bou Dagher-Kharrat, M. (2015) Lebanon Flora. http://www.lebanon-flora.org/ Accessed 15 January 15.

Boulos, L. & Al-Dosari, M. (1994) Checklist of the flora of Kuwait. Journal of the University of Kuwait (Science), 21, 203–2017.

Bowdoin Scientific Station (2011) Vascular plants of Kent Island. https://www.bowdoin.edu/kent-island/species/plants.shtml Accessed 14 September 11.

BRAHMS online (2014) Plants of Gabon. http://herbaria.plants.ox.ac.uk/bol/gabon Accessed 29 January 15.

Breckle, S.-W., Hedge, I.C. & Rafiqpoor, M.D. (2013) Vascular plants of Afghanistan: An augmented checklist. Scientia Bonnensis, Bonn.

Brennan, K. (1996) An annotated checklist of the vascular plants of the Alligator Rivers Region, Northern Territory, Australia. Supervising Scientist, Barton, Australia.

Bridgewater, S.G.M., Harris, D.J., Whitefoord, C., Monro, A.K., Penn, M.G., Sutton, D.A., Sayer, B., Adams, B., Balick, M.J., Atha, D.H., Solomon, J. & Holst, B.K. (2006) A Preliminary Checklist of the vascular plants of the Chiquibul Forest, Belize. Edinburgh Journal of Botany, 63, 269–321.

Brofas, G., Karetsos, G., Panitsa, M. & Theocharopoulos, M. (2001) The flora and vegetation of Gyali Island, SE Aegean, Greece. Willdenowia, 31, 51–70.

Broughton, D.A. & McAdam, J.H. (2005) A checklist of the native vascular flora of the Falkland Islands (Islas Malvinas): new information on the species present, their ecology, status and distribution. The Journal of the Torrey Botanical Society, 132, 115–148.

Brown, E.A. (1979) Vegetation and flora of Ponui Island, Hauraki Gulf, New Zealand. TANE, 25, 5–16.

Brundu, G. & Camarda, I. (2013) The Flora of Chad: a checklist and brief analysis. Phytokeys, 23, 1–17.

Buchwald, E., Wind, P., Bruun, H.H., Møller, P.F., Ejrnæs, R. & Svart, H.E. (2013) Hvilke planter er hjemmehørende i Danmark? Jydsk Naturhistorisk Forening, 118, 72–96.

Bundesamt für Naturschutz (2016) Floraweb. www.floraweb.de Accessed 15 September 16.

Burke Museum of Natural History and Culture (2012) Russian Far East Flora Database. http://biology.burke.washington.edu/herbarium/okhotskia/search.php Accessed 1 October 12.

Burton, R.M. (1991) A check-list and evaluation of the flora of Nisyros (Dodecanese, Greece). Willdenowia, 20, 15–38.

Butler, B.J., Barclay, J.S. & Fisher, J.P. (1999) Plant communities and flora of Robins Island (Long Island), New York. Journal of the Torrey Botanical Society, 126, 63–76.

Buzzard, V., Hulshof, C.M., Birt, T., Violle, C., Enquist, B.J. & Larjavaara, M. (2016) Re-growing a tropical dry forest: Functional plant trait composition and community assembly during succession. Functional Ecology, 30, 1006–1013.

Byrd, G.V. (1984) Vascular vegetation of Buldir Island, Aleutian Islands, Alaska, compared to another Aleutian Island. Arctic, 37, 37–48.

Cameron, E.K. (1982) Vascular plants of an unclassified islet, Cape Brett Peninsula, northern New Zealand. TANE (28), 213–220.

Cameron, E.K. (1984) Vascular plants of the three largest Chickens (Marotere) islands: Lady Alice, Whatupuke, Coppermine: North-East New Zealand. TANE, 30, 53–75.

Cameron, E.K. (1985) Vascular flora and vegetation of Rimariki and associated islands, Mimiwhangata, north-east New Zealand. TANE, 31, 47–74.

Cameron, E.K. (1990) Vascular plants of the main northern Mokohinau Islands, North-East New Zealand. TANE, 32, 113–130.

Cameron, E.K. (1991) Paratahi Island - Karekare, West Auckland. Auckland Botanical Society Journal, 46, 84–85.

Cameron, E.K., Lange, P.J. de, McCallum, J., Taylor, G.A. & Bellingham, P.J. (2007) Vascular flora and some fauna for a chain of six Vascular flora and some fauna for a chain of six Hauraki Gulf islands east and southeast of Waiheke Island. Auckland Botanical Society Journal, 62, 136–156.

Cameron, E.K. & Taylor, G.A. (1997) Flora and fauna of Sentinel Rock, Mangawhai Heads, Northern New Zealand. TANE, 36, 15– 25.

CARMABI (2009) Dutch Caribbean Biodiversity Explorer. http://www.dcbiodata.net/explorer/home accessed 24 June 11.

Cascante-Marín, A. & Estrada-Chavarría, A. (2012) Las plantas vasculares de El Rodeo, Costa Rica. Brenesia, 77, 71–128.

Case, T.J., Cody, M.L. & Ezcurra, E. (2002) A new island biogeography of the Sea of Cortés. Oxford University Press, New York, NY.

Catarino, L., Martins, E.S., Basto, M.F. & Diniz, M.A. (2008) An annotated checklist of the vascular flora of Guinea-Bissau (West Africa). Blumea-Biodiversity, Evolution and Biogeography of Plants, 53, 1–222.

Chandra, R., Prusty, B Anjan Kumar & Azeez, P.A. (2011) A revised Checklist of the Flora of Keoladeo National Park, a World Heritage Site in India. Environmental Research Journal, 5, 331–348.

Chang, C.-S., Kim, H. & Chang, K. Provisional Checklist of the Vascular Plants for the Korea Peninsular Flora (KPF): Version 1.0, Korea.

Charters, M. (2007) Flora of Bermuda. http://www.calflora.net/floraofbermuda/ Accessed 30 July 11.

Chatelain, C., Assi, L.A., Spichiger, R. & Gautier, L. (2011) Cartes de distribution des plantes de Côte d'Ivoire. Conservatoire et Jardin botaniques de la Ville de Genève, Chambésy.

Chawla, A., Parkash, O., Sharma, V., Rajkumar, S., Lal, B., Gopichand, Singh, R.D. & Thukral, A.K. (2012) Vascular plants, Kinnaur, Himachal Pradesh, India. Check list, 8, 321–348.

Cheffings, C.M. & Farrell, L. (eds) (2005) The vascular plant red data list for Great Britain. Joint Nature Conservation Committee, Peterborough.

Chen, S.-C. & Moles, A.T. (2015) A mammoth mouthful? A test of the idea that larger animals ingest larger seeds. Global Ecology and Biogeography, 24, 1269–1280.

Chernyaeva, A.M. (1973) Flora of Onekotan Island. Bulletin of Main Botanical Garden, 87, 21–29.

Chiapella, J. & Ezcurra, C. (1999) La flora del parque provincial Tromen, provincia de Neuquén, Argentina. Multequina, 8, 51–60.

Chinese Virtual Herbarium (2016) The Flora of China v. 5.0. http://www.cvh.org.cn/ Accessed 15 January 16.

Chong, K.Y., Tan, T.W.H. & Corlett, R.T. (2009) A checklist of the total vascular plant flora of Singapore: Native, Naturalised and Cultivated Species. Raffles Museum of Biodiversity Research, Singapore.

Christmas Island National Park (2002) Third Christmas Island national park management plan. Parks Australia North, Christmas Island, Australia.

Christodoulakis, D. (1996) The flora of Ikaria (Greece, E. Aegean Islands). Phyton, 36, 63–91.

Christophersen, E. (1931) Vascular plants of Johnston and Wake Islands. Occasional Papers of the Bernice Pauahi Bishop Museum of Polynesian, 9, 1–20.

Clapp, R.B. & Sibley, F.C. (1971) The vascular flora and terrestrial vertebrates of Vostok Island, South-Central Pacific. Atoll Research Bulletin, 144, 1–10.

Clark, J.L., Neill, D.A. & Asanza, M. (2006) Floristic checklist of the Mache-Chindul mountains of Northwestern Ecuador. Contributions from the United States National Herbarium, 54, 1–180.

Cochard, R. & Bloesch, U. (2007) Electronic plant species database of the Saadani National Park, coastal Tanzania. http://www.wildlife-baldus.com/saadani.html Accessed 14 November 16.

CONABIO (2016) Sistema Nacional de Información sobre Biodiversidad. https://www.gob.mx/conabio Accessed 11 April 16.

Conti, F., Abbate, G., Alessandrini, A. & Blasi, C. (2005) Annotated Checklist of the Italian Vascular Flora. Palombi Editori, Roma, Italy.

Conti, F. & Bartolucci, F. (2015) The Vascular Flora of the National Park of Abruzzo, Lazio and Molise (Central Italy). Springer International Publishing, Cham.

Convey, P., Lewis Smith, R.I., Hodgson, D.A. & Peat, H.J. (2000) The flora of the South Sandwich Islands, with particular reference to the influence of geothermal heating. Journal of Biogeography, 27, 1279–1295.

Costion, C. & Lorence, D. (2012) The endemic plants of Micronesia: A geographical checklist and commentary. Micronesica, 43, 51–100.

Court, D.J. & Hardacre, A. K., Lynch, P.A. (1973) The vegetation of the aldermen islands: a reappraisal. TANE, 19, 41–60.

Cronk, Q.C.B. (1989) The past and present vegetation of St Helena. Journal of Biogeography, 16, 47–64.

Cuc Phuong National Park (2011) Flora of the Cuc Phuong National Park, Vietnam.

Da Vela, M., Frignani, F., Bonari, G. & Angiolini, C. (2013) La flora vascolare della diserva naturale "La Pietra" (Toscana meridionale). Micologia e vegetazione mediterranea, 28, 135–160.

Danihelka, J., Chrtek, J. & Kaplan, Z. (2012) Checklist of vascular plants of the Czech Republic. Preslia, 84, 647–811.

D'Arcy, W.G. (1971) The island of Anegada and its flora. Atoll Research Bulletin, 139, 1–21.

Datar, M.N. & Lakshminarasimhan, P. (2013) Check List of Wild Angiosperms of Bhagwan Mahavir (Molem) National Park, Goa, India. Check list, 9, 186–207.

Dauby, G., Leal, M. & Stevart, T. (2008) Vascular plant checklist of the coastal National Park of Pongara, Gabon. Systematics and geography of plants, 78, 155–216.

Dauby, G., Zaiss, R., Blach-Overgaard, A., Catarino, L., Damen, T., Deblauwe, V., Dessein, S., Dransfield, J., Droissart, V., Duarte, M.C., Engledow, H., Fadeur, G., Figueira, R., Gereau, R.E., Hardy, O.J., Harris, D.J., Heij, J. de, Janssens, S., Klomberg, Y., Ley, A.C., MacKinder, B.A., Meerts, P., van de Poel, J.L., Sonké, B., Sosef, M.S.M., Stévart, T., Stoffelen, P., Svenning, J.-C., Sepulchre, P., van der Burgt, X., Wieringa, J.J. & Couvreur, T.L.P. (2016) RAINBIO: A mega-database of tropical African vascular plants distributions. PhytoKeys, 74, 1–18.

de la Luz, León, Rebman, J., Domínguez-León, M. & Domínguez-Cadena, R. (2008) The vascular flora and floristic relationships of the sierra de la Giganta in Baja California Sur, México. Revista mexicana de biodiversidad, 79, 29–65.

de Medeiros, Marcelo Brilhante, Walter, B.M.T., Da Silva, Glocimar Pereira, Gomes, B.M., Lima, Isabela Lustz Portela, Silva, S.R., Moser, P., Oliveira, W.L. & Cavalcanti, T.B. (2012) Vascular flora of the Tocantins river middle basin, Brazil. Check list, 8, 852–885.

de V. Barbosa, Maria Regina, Thomas, W.M.W., Zárate, E.L.d.P., de Lima, R.B., de Fátima Agra, M., de Lima, I.B., Pessoa, M.d.C.R., Lourenço, A.R.L., Delgado Júnior, G.C., de Pontes, R.A., Chagas, E.C.O., Viana, J.L., Gadelha Neto, P.d.C., Araújo, C.M.L.R., Araújo, A.d.A.M., de Freita, G.B., Lima, J.R., Silva, F.O., Vieira, L.d.A.F., Pereira, L.d.A., Costa, R.M.T., Duré, R.C. & Sá, M.d.G.V.d. (2011) Checklist of the vascular plants of the Guaribas Biological Reserve, Paraíba, Brazil. Revista Nordestina de Biologia, 20, 79–106.

Desmet, P. & Brouillet, L. (2013) Database of Vascular Plants of Canada (VASCAN): a community contributed taxonomic checklist of all vascular plants of Canada, Saint Pierre and Miquelon, and Greenland. Phytokeys, 25, 55–67.

Dimopoulos, P., Raus, T., Bergmeier, E., Constantinidis, T., Iatrou, G., Kokkini, S., Strid, A. & Tzanoudakis, D. (2013) Vascular plants of Greece: An annotated checklist. Botanischer Garten und Botanisches Museum Berlin-Dahlem, Freie Universität Berlin; Hellenic Botanical Society, Berlin, Athens.

Directorate of Wrangel Island Reserve (2003) Natural System of Wrangel Island Reserve. The World Heritage Committee, Paris, France.

Dobignard, A. & Chatelain, C. (2010-2013) Synonymic and bibliographic index of North African plants (Vol. 1-5). http://www.ville-ge.ch/musinfo/bd/cjb/africa/index.php?langue=an.

Domínguez, E., Marticorena, C., Elvebakk, A. & Pauchard, A. (2004) Catálogo de la flora vascular del Parque Nacional Pali Aike. XII Región, Chile. Gayana Botánica, 61, 67–72.

Doroftei, M., Oprea, A., & Ştefan, N. & Sârbu, I. (2011) Vascular wild flora of Danube Delta Biosphere Reserve. Sci. Annals of Danube Delta Institute, 17, 15–52.

Dostálek, J. & Kučera, J. (2011) Flóra a vegetace národní přírodní rezervace Bukačka v Orlických horách. Acta Musei Reginaehradecensis, 33, 15–36.

Dowhan, J.J. & Rozsa, R. (1989) Flora of Fire Island, Suffolk County, New York. Bulletin of the Torrey Botanical Club, 116, 265–282.

Drummond, R.B. & Mapaure, I. (1994) List of Flowering Plants and Ferns (of Chirinda Forest). http://www.zimbabweflora.co.zw/speciesdata/checklist-display.php?checklist_code=4 Accessed 20 January 15.

Du Puy, D.J. (1993) Christmas Island: species lists. http://www.anbg.gov.au/abrs/online-resources/flora/ Accessed 6 April 11.

Easley, M.C. & Judd, W.S. (1993) Vascular Flora of Little Talbot Island, Duval County, Florida. Castanea, 58, 162–177.

Edwards, J.A. (1980) An experimental introduction of vascular plants from South Georgia to the maritime Antarctic. British Antarctic Survey Bulletin, 49, 73–80.

Egea, J. de, Peña-Chocarro, M., Espada, C. & Knapp, S. (2012) Checklist of vascular plants of the Department of Ñeembucú, Paraguay. Phytokeys, 9, 15–179.

Egorova, E.M. (1964) Flora of Shiashkotan Island. Bulletin of the Main Botanical Garden, 54, 114–120.

Ejaz-ul-Islam Dar, M., Cochard, R., Shrestha, R.P. & Ahmad, S. (2012) Floristic composition of Machiara National Park, District Muzaffarabad Azad Kashmir, Pakistan. International Journal of Biosciences, 2, 28–45.

Eleftheriadou, E. & Raus, T. (1996) The vascular flora of the nature reserve Frakto Virgin Forest of Nomos Dramas (E Makedonia, Greece). Willdenowia, 25, 455–485.

El-Ghani, M.M. & Abdel-Khalik, K.N. (2005) Floristic Diversity and Phytogeography of the Gebel Elba National Park, South-East Egypt. Turkish Journal of Botany, 30, 121–136.

Engemann, K., Sandel, B., Boyle, B.L., Enquist, B.J., Jørgensen, P.M., Kattge, J., McGill, B.J., Morueta-Holme, N., Peet, R.K., Spencer, N.J., Violle, C., Wiser, S.K. & Svenning, J.-C. (2016) A plant growth form dataset for the New World. Ecology, 97 (11), 3243.

Esler, A.E. (1978) Botanical features of the Mokohinau Islands. TANE, 24, 187–197.

Espinosa-Jiménez, J.A., Pérez-Farrera, M.Á. & Martínez-Camilo, R. (2011) Inventario florístico del Parque Nacional Cañón del Sumidero, Chiapas, México. Boletín de la Sociedad Botánica de México, 89, 37–82.

Evarts-Bunders, P., Evarte-Bundere, G., Bāra, J. & Nitcis, M. (2013) The flora of vascular plants in nature reserve „Eglone”. Acta Biologica Universitatis Daugavpiliensis, 13 (2), 21–38.

Exell, A.W. (1944) Catalogue of the vascular plants of S. Tome (with Principe and Annobon). Trustees of the British Museum, London, UK.

Fern, K. & Fern, A. (2015) Useful Tropical Plants Database Accessed 25 July 15.

Ferreira, A.L., Coutinho, B.R., Pinheiro, H.T. & Thomaz, L.D. (2007) Composição florística e formações vegetais da Ilha dos Franceses, Espírito Santo. Bol. Mus. Biol. Mello Leitao, 22, 25–44.

Figueiredo, E., Paiva, J., Stevart, T., Oliveira, F. & Smith, G.F. (2011) Annotated catalogue of the flowering plants of São Tomé and Príncipe. Bothalia, 41, 41–82.

Figueiredo, E. & Smith, G.F. (2008) Plants of Angola / Plantas de Angola. http://floras.cenapad.unicamp.br/floradeangola.

Figueroa-C., Y. & Galeano, G. (2007) Lista comentada de las plantas vasculares del enclave seco interandino de La Tatacoa (Huila, Colombia). Caldasia, 29, 263–281.

Fischer, E., Rembold, K., Althof, A., Obholzer, J., Malombe, I., Mwachala, G., Onyango, J.C., Dumbo, B. & Theisen, I. (2010) Annotated checklist of the vascular plants of Kakamega Forest, Western Province, Kenya. Journal of East African Natural History, 99, 129–226.

Fischer, M.A., Adler, W. & Oswald, K. (2008) Exkursionsflora für Österreich, Liechtenstein und Südtirol: Bestimmungsbuch für alle in der Republik Österreich, im Fürstentum Liechtenstein und in der Autonomen Provinz Bozen, 3., verb. u. erw. Aufl. der "Exkursionsflora von Österreich" edn. Ö Landesmuseum, Linz.

Flood, P.G. & Heatwole, H. (1986) Coral Cay instability and species-turnover of plants at Swain Reefs, Southern Great Barrier Reef, Australia. Journal of Coastal Research, 2, 479–496.

Florence, J., Chevillotte, H., Ollier, C. & Meyer, J.-Y. (2007) Base de données botaniques Nadeaud de l'Herbier de la Polynésie française (PAP). http://www.herbier-tahiti.pf/ Accessed 1 July 11.

Forester, L.J. & Anderson, P.J. (1995) Vascular plants, vegetation and wildlife of Matapia Island, Far North, New Zealand. TANE, 35, 39–50.

Fosberg, F.R. (1949) Flora of Johnston Island, Central Pacific. Pacific Science, 3, 338–339.

Fosberg, F.R. (1985) Vegetation of Bikini Atoll. Atoll Research Bulletin, 315, 1–28.

Fosberg, F.R., Otobed, D., Sachet, M.H., Oliver, R.L., Powell, D.A. & Canfield, J.E. (1980) Vascular plants of Palau with vernacular names. Smithsonian Institution, Department of Botany, Washington DC.

Fosberg, F.R., Renvoize, S.A. & Townsend, C.C. (1980) The flora of Aldabra and neighbouring islands. HMSO, London, UK.

Fosberg, F.R. & Sachet, M.H. (1987) Flora of Maupiti, Society Islands. Atoll Research Bulletin, 294, 1–70.

Fosberg, F.R., Stoddart, D.R., Sachet, M.-H. & Spellman, D.L. (1982) Plants of the Belize cays. Atoll Research Bulletin, 258, 1–77.

Franklin, J., Keppel, G. & Whistler, W.A. (2008) The vegetation and flora of Lakeba, Nayau and Aiwa islands, central Lau Group, Fiji. Micronesica, 40, 169–225.

Frenot, Y., Gloaguen, J.C., Masse, L. & Lebouvier, M. (2001) Human activities, ecosystem disturbance and plant invasions in subantarctic Crozet, Kerguelen and Amsterdam Islands. Biological Conservation, 101, 33–50.

Fuentes Parada, N. (2016) ALARM project: Chilean vascular plants. unpublished.

Funk, V.A., Hollowell, T., Berry, P., Kelloff, C. & Alexander, S.N. (2007) Checklist of the plants of the Guiana Shield (Venezuela: Amazonas, Bolivar, Delta Amacuro; Guyana, Surinam, French Guiana). Department of Botany, National Museum of Natural History, Washington, DC.

Gabrielsen, G.W., Brekke, B., Alsos, I.G. & Hansen, J.R. (1997) Natur-og kulturmiljøet på Jan Mayen. Norsk Polarinstitutt, Oslo, Norway.

Gafurova, М.М. (2014) Vascular plants of Chuvash Republic: Flora of the Volga River Basin V. III. The Russian Academy of Sciences.

Gage, S., Joneson, S.L., Barkalov, V.Y., Eremenko, N.A. & Takahashi, H. (2006) A newly compiled checklist of the vascular plants of the Habomais, the Little Kurils. Bulletin of the Hokkaido University Museum, 3, 67–91.

Gamit, S.B., Maurya, R.R., Qureshimatva, U.M. & Solanki, H.A. (2015) Check list of flowering plants in Tapi District, Gujarat, India. International Journal of advanced Research, 3, 1104–1123.

Gerlach, J. (2003) The biodiversity of the granitic islands of Seychelles. Phelsuma, 11 (Supplement A), 1–47.

Gnoumou, A., Ouedraogo, O., Schmidt, M. & Thiombiano, A. (2015) Floristic diversity of classified forest and partial faunal reserve of Comoé-Léraba, southwest Burkina Faso. Check list, 11, 1557.

Gray, A., Pelembe, T. & Stroud, S. (2005) The conservation of the endemic vascular flora of Ascension Island and threats from alien species. Oryx, 39, 449–453.

Green, P.S. (1994a) Lord Howe Island: species lists. http://www.environment.gov.au/biodiversity/abrs/online-resources/flora/49/index.html accessed 6 April 11.

Green, P.S. (1994b) Norfolk Island: Species lists. http://www.environment.gov.au/biodiversity/abrs/online-resources/flora/49/index.html accessed 6 April 11.

Greene, S.W. & Walton, D.W.H. (1975) An annotated check list of the sub-Antarctic and Antarctic vascular flora. Polar Record, 17, 473–484.

Grozeva, N.H. (2005) The flora of Atanasovsko lake natural reserve. Proceedings of the Balkan Scientific Conference of Biology (eds B. Gruev, M. Nikolova & A. Donev), 381–396.

Harris, D.J. (2002) The vascular plants of the Dzanga-Sangha Reserve, Central African Republic. Royal Botanic Garden Edinburgh, Edinburgh, Scotland, UK.

Hauenstein, E., Muñoz-Pedreros, A., Yánez, J., Sánchez, P., Möller, P., Guiñez, B. & Gil, C. (2009) Flora y vegetación de la Reserva Nacional Lago Peñuelas, Reserva de la Biósfera, Región de Valparaíso, Chile. Bosque (Valdivia), 30, 159–179.

Hawkins, B.A., Rueda, M., Rangel, T.F., Field, R., Diniz-Filho, J.A.F. & Linder, P. (2013) Community phylogenetics at the biogeographical scale: cold tolerance, niche conservatism and the structure of North American forests. Journal of Biogeography, 41, 23–38.

Heads, M. (2006) Seed plants of Fiji: an ecological analysis. Biological Journal of the Linnean Society, 89, 407–431.

Heatwole, H., Levins, R. & Byer, M.D. (1981) Biogeography of the Puerto Rican Bank. Atoll Research Bulletin, 251, 1–55.

Heatwole, H. & Levins R (1973) Biogeography of the Puerto Rican Bank: Species-Turnover on a Small Cay, Cayo Ahogado. Ecology, 54, 1042–1055.

Heberling, J.M., Jo, I., Kozhevnikov, A., Lee, H. & Fridley, J.D. (2017) Biotic interchange in the Anthropocene: Strong asymmetry in East Asian and eastern North American plant invasions. Global Ecology and Biogeography, 26, 447–458.

Hill, M.J. (ed) (2002) Biodiversity surveys and conservation potential of inner Seychelles islands. Smithsonian Institution, Washington, DC.

Hill, S.R. (1986) An annotated checklist of the vascular flora of Assateague Island (Maryland and Virginia). Castanea, 51, 265–305.

Hnatiuk, R.J. (1993) Subantarctic Islands: species lists. http://www.environment.gov.au/biodiversity/abrs/online-resources/flora/50/index.html accessed 7 April 11.

Hoang, V.S. (2009) Uses and conservation of plant diversity in Ben En National Park, Vietnam. Ph.D., National Herbarium of the Netherlands, Leiden University Branch.

Hoke, P., Demey, R. & Peal, A. (2007) A Rapid Biological Assessment of North Lorma, Gola and Grebo National Forests, Liberia. Conservation International, Arlington, USA.

Hyde, M.A., Wursten, B.T., Ballings, P. & Coates Palgrave, M. (2016a) Flora of Botswana. http://www.botswanaflora.com/ Accessed 14 November 16.

Hyde, M.A., Wursten, B.T., Ballings, P. & Coates Palgrave, M. (2016b) Flora of Caprivi. http://www.capriviflora.com/ Accessed 14 November 16.

Hyde, M.A., Wursten, B.T., Ballings, P. & Coates Palgrave, M. (2016c) Flora of Malawi. http://www.malawiflora.com/index.php Accessed 14 November 16.

Hyde, M.A., Wursten, B.T., Ballings, P. & Coates Palgrave, M. (2016d) Flora of Mozambique. http://www.mozambiqueflora.com/ Accessed 14 November 16.

Hyde, M.A., Wursten, B.T., Ballings, P. & Coates Palgrave, M. (2016e) Flora of Zimbabwe Accessed 14 November 16.

ICIMOD (2015) Hindu Kush-Himalayan (HKH) Conservation Portal. http://www.icimod.org/hkhconservationportal/Initiatives.aspx Accessed 21 April 15.

Imada, C.T. (ed) (2012) Hawaiian Native and Naturalized Vascular Plants Checklist, (December 2012 update) edn. Hawaii Biological Survey, Bishop Museum, Honolulu, Hawai‘i.

INBIO (2000) Lista de planta de Costa Rica: With updates by Eduardo Chacón. http://www.inbio.ac.cr/papers/manual_plantas/index.html Accessed 5 February 16.

Iwatsuki, K., Yamazaki, T., Boufford, D.E. & Ohba, H. (1995) Flora of Japan, Vol. 1 - Pteridophyta and Gymnospermae. Kodansha, Tokyo, Japan.

Jackes, B.R. (2010) Plants of Magnetic Island, 3rd edn. James Cook University, Townsville, Australia.

Jahn, R. & Schönfelder, P. (1995) Exkursionsflora für Kreta. Ulmer (Eugen), Stuttgart, Germany.

Jaramillo Díaz, P. & Guézou, A. (2011) CDF checklist of Galapagos vascular plants. http://www.darwinfoundation.org/datazone/checklists/vascular-plants/ Accessed 2 February 11.

Jardim Botânico do Rio de Janeiro (2016) Flora do Brasil 2020 em construção. http://floradobrasil.jbrj.gov.br/ Accessed 9 May 16.

Jasprica, N., Dolina, K. & Milović, M. (2015) Plant taxa and communities on three islets in south Croatia, NE Mediterranean. Natura Croatica, 24, 191–213.

Jiménez, J.E., Juárez, P. & Díaz, A. (2016) Checklist of the vascular flora of Reserva Biológica San Luis, Costa Rica. Check list, 12, 1859.

Johnson, P.N. & Campbell, D.J. (1975) Vascular plants of the Auckland Islands. New Zealand journal of botany, 13, 665–720.

Johnston, I.M. (1931) The flora of the Revillagigedo Islands. Proceedings of the California Academy of Sciences, 20, 9–104.

Jordano, P. (2008) FRUBASE. http://ebd10.ebd.csic.es/mywork/frubase/frubase.html Accessed 26.06.15.

Junak, S., Philbrick, R., Chaney, S. & Clark, R. (1997) A checklist of vascular plants of Channel Islands National Park, 2nd edn. Southwest Parks and Monuments Association, Tucson, Arizona.

Kabuye, C.H.S., Mungai, G.M. & Mutangah, J.G. (1986) Flora of Kora National Reserve. Kora: An Ecological Inventory of the Kora National Reserve, Kenya: Kora Research Project 1982-85 a Joint Venture Between the National Museums of Kenya and the Royal Geographical Society (eds M. Coe & N.M. Collins), pp. 57–104. Royal Geographical Society.

Kamari, G., Phitos, D., Snogerup, B. & Snogerup, S. (1988) Flora and vegetation of Yioura, N Sporades, Greece. Willdenowia, 17, 59–85.

Karlsson, T. (2011) Lista över Östergötlands kärlväxter. http://ostgotaflora.se/handledning.html accessed 17 May 16.

Karlsson, T. & Agestam, M. (2014) Checklist of Nordic vascular plants: Sweden Checklist. http://www.euphrasia.nu/checklista/index.eng.html Accessed 11 February 16.

Kartesz, J.T. (2014) The Biota of North America Program (BONAP): North American Plant Atlas. http://www.bonap.org/napa.html.

Kattge, J., Diaz, S., Lavorel, S., Prentice, I.C., Leadley, P., Bönisch, G., Garnier, E., Westoby, M., Reich, P.B. & Wright, I.J. (2011) TRY – a global database of plant traits. Global Change Biology, 17 (9), 2905–2935.

Kelloff, C.L. & Funk, V.A. (1998) Preliminary checklist of the plants of Kaieteur National Park, Guyana. National Museum of Natural History, Smithsonian Institution, Washington.

Kelly, L. (2006) The vascular flora of Huggins Island, Onslow County, North Carolina. Castanea, 71 (4), 295–311.

Kemenes, A. (2003) Distribuição espacial da flora terrestre fanerogâmica do Parque Nacional Marinho de Abrolhos, BA. Revista Brasil. Bot, 26, 141–150.

Kenneally, K.F. (1993) Ashmore Reef and Cartier Island: Species lists. http://www.environment.gov.au/biodiversity/abrs/online-resources/flora/50/index.html accessed 6 April 11.

Keppel, G. (2005) Summary report on forests of the Mataqali Nadicake Kilaka, Kubulau district, Bua, Vanua Levu.

Keppel, G., Gillespie, T.W., Ormerod, P. & Fricker, G.A. (2016) Habitat diversity predicts orchid diversity in the tropical south-west Pacific. Journal of Biogeography, 43, 2332–2342.

Kerguelen, M. (2005) Base de Données Nomenclaturales de la Flore de France. http://www.tela-botanica.org/page:284# Accessed 13 January 12.

Khanina, L., Zaugolnova, L., Smirnova, O., Shovkun, M. & Glukhova, E. (2016) Flora of vascular plants in the Central European Russia. http://www.impb.ru/eco/index.php?l=en Accessed 6 September 17.

Kim, C.-S., Koh, J.-G., Moon, M.-O., Song, G.-P., Hyun, H.-J., Song, K.-M. & Kim, M.-H. (2007) Flora and life form spectrum of Hallasan natural reserve, Korea. Journal of the Environmental Sciences, 16, 1257–1269.

Kim, H.-J., Ji, S.-J., Jung, S.-Y., Park, S.H., Lee, S.-G., Lee, C.-W. & Chang, K.S. (2015) Flora of Vascular Plants in Deokjeokdo (Ongjingun) and Its Adjacent Regions, Korea. Korean Journal of Plant Resources, 28, 487–510.

Kingston, N., Waldren, S. & Bradley, U. (2003) The phytogeographical affinities of the Pitcairn Islands – a model for south-eastern Polynesia? Journal of Biogeography, 30 (9), 1311–1328.

Kirchner, F., Picot, F., Merceron, E. & Gigot, G. (2010) Flore vasculaire de La Réunion. Conservatoire Botanique National de Mascarin, Réunion, France.

Kleyer, M., Bekker, R.M., Knevel, I.C., Bakker, J.P., Thompson, K., Sonnenschein, M., Poschlod, P., van Groenendael, J.M., Klimeš, L., Klimešová, J., Klotz, S., Rusch, G.M., Hermy, M., Adriaens, D., Boedeltje, G., Bossuyt, B., Dannemann, A., Endels, P., Götzenberger, L., Hodgson, J.G., Jackel, A.-K., Kühn, I., Kunzmann, D., Ozinga, W.A., Römermann, C., Stadler, M., Schlegelmilch, J., Steendam, H.J., Tackenberg, O., Wilmann, B., Cornelissen, J.H.C., Eriksson, O., Garnier, E. & Peco, B. (2008) The LEDA Traitbase: a database of life-history traits of the Northwest European flora. Journal of Ecology, 96, 1266–1274.

Klimeš, L. & Dickoré, B. (2015) Flora of Ladak (NW Himalaya): A preliminary check-list. http://www.butbn.cas.cz/klimes/desert.html Accessed 12 January 15.

Koltzenberg, M. (2002) Flora von Kamtschatka: Vorläufige Checkliste.

Koltzenburg, M. (2011) Checkliste der Gefässpflanzen Irlands. http://www.saxifraga.de/eire/irl_artenlisten_gp.html Accessed 4 April 11.

Kraaij, T. (2011) The flora of the Bontebok National Park in regional perspective. South African Journal of Botany, 77, 455–473.

Kraft, T.S., Wright, S.J., Turner, I., Lucas, P.W., Oufiero, C.E., Supardi Noor, M.N., Sun, I.-F., Dominy, N.J. & Whitney, K. (2015) Seed size and the evolution of leaf defences. Journal of Ecology, 103, 1057–1068.

Krestov, P.V. (2016) Flora of Siberia.

Kristinsson, H. (2008) Checklist of the vascular plants of Iceland. Náttúrufræðistofnun Íslands, Reykjavík, Iceland.

Kumar, A., Bajpai, O., Mishra, A.K., Sahu, N., Behera, S.K., Bargali, S.S. & Chaudhary, L.B. (2015) A checklist of the flowering plants of Katerniaghat Wildlife Sanctuary, Uttar Pradesh, India. Journal of Threatened Taxa, 7, 7309–7408.

Kupriyanov, A.N. & Kupriyanov, O.A. The study of the flora (by the example of Kemerovo region), Kemerovo.

Kuzmenkova, S.M. (2015) Plants of Belarus. http://hbc.bas-net.by/plantae/eng/default.php Accessed 15 February 16.

Laliga, L.S., Benavent,, J.E.O., Conca, A., Signes, J.X.S. & Nebot, J.R. (2012) Catálogo de la flora del Parque Natural de la Sierra de Mariola (Alicante-Valencia). Flora Montiberica, 51, 97–125.

Lange, P.J. de (1994) The Flora of Gannet Island (Karewa), Tasman Sea, Western North Island. Wellington Botanical Society Bulletin, 46, 63–69.

Lange, P.J. de (2015) The Flora and Vegetation of L’Esperance Rock,Southern Kermadec Islands. Bulletin of the Auckland Museum, 20, 231–242.

Lange, P.J. de & Cameron, E.K. (1999) The vascular flora of Aorangi Island, Poor Knights Islands, northern New Zealand. New Zealand journal of botany, 37, 433–468.

Lange, P.J. de, Cameron, E.K. & Taylor, G.A. (1995) Flora and fauna of Tatapihi (Groper) Island, Mokohinau Islands. TANE, 35, 69–94.

Lange, P.J. de, Heenan, P.B. & Rolfe, J.R. (2011) Checklist of vascular plants recorded from Chatham Islands. Department of Conservation, Wellington, New Zealand.

Lazkov, G.A. & Sultanova, B.A. (2011) Checklist of vascular plants of Kyrgyzstan. Botanical Museum, Finnish Museum of Natural History, Helsinki.

Le Houerou, H.N. (2004) Plant diversity in Marmarica (Libya & Egypt): a catalogue of the vascular plants reported with their biology, distribution, frequency, usage, economic potential, habitat and main ecological features, with an extensive bibliography. Candollea, 59, 259–308.

Lee, R.-Y., Jang, H.-D., Kim, Y.-Y., Yang, S.-G., Choi, H.-J., Ji, S.-J. & Oh, B.-U. (2014) Flora of vascular plants in the Chilgapsan Provincial Park, Korea. Journal of Asia-Pacific Biodiversity, 7, 237–247.

Lee, S.-M., Lee, H.-Y., Lee, Y.-M., Park, S.-H., Lee, B.-C. & Lim, W.-H. (2008) Flora of Gyeongju National Park, Korea. Journal of Korean Nature, 1, 21–38.

Lester-Garland, L.V. (1903) A flora of the islands of Jersey: with a list of the plants of the Channel Islands in general, and remarks upon their distribution and geographical affinities. West, Newman & Co, London, UK.

Letcher, S.G., Lasky, J.R., Chazdon, R.L., Norden, N., Wright, S.J., Meave, J.A., Pérez-García, E.A., Muñoz, R., Romero-Pérez, E., Andrade, A., Andrade, J.L., Balvanera, P., Becknell, J.M., Bentos, T.V., Bhaskar, R., Bongers, F., Boukili, V., Brancalion, P.H.S., César, R.G., Clark, D.A., Clark, D.B., Craven, D., DeFrancesco, A., Dupuy, J.M., Finegan, B., González-Jiménez, E., Hall, J.S., Harms, K.E., Hernández-Stefanoni, J.L., Hietz, P., Kennard, D., Killeen, T.J., Laurance, S.G., Lebrija-Trejos, E.E., Lohbeck, M., Martínez-Ramos, M., Massoca, P.E.S., Mesquita, R.C.G., Mora, F., Muscarella, R., Paz, H., Pineda-García, F., Powers, J.S., Quesada-Monge, R., Rodrigues, R.R., Sandor, M.E., Sanaphre-Villanueva, L., Schüller, E., Swenson, N.G., Tauro, A., Uriarte, M., van Breugel, M., Vargas-Ramírez, O., Viani, R.A.G., Wendt, A.L., Williamson, G.B. & Zhou, S. (2015) Environmental gradients and the evolution of successional habitat specialization: A test case with 14 Neotropical forest sites. Journal of Ecology, 103, 1276–1290.

Levin, G.A. & Moran, R. (1989) The vascular flora of Socorro, Mexico. Memoirs of the San Diego Society of Natural History, 16, 1–71.

Limbu, D., Koirala, M. & Shang, Z. (2013) A Checklist of Angiospermic Flora of Tinjure-Milke-Jaljale, Eastern Nepal. Nepal Journal of Science and Technology, 13, 87–96.

Linhart, Y.B. (1980) Local biogeography of plants on a Caribbean atoll. Journal of Biogeography, 7, 159–171.

Lipkin, R. (2005) Aniakchak National Monument and Preserve, vascular plant inventory: final technical report. National Park Service, Southwest Alaska Network Inventory & Monitroing Program, Anchorage, USA.

Lopez-Martinez, J.O., Sanaphre-Villanueva, L., Dupuy, J.M., Hernandez-Stefanoni, J.L., Meave, J.A. & Gallardo-Cruz, J.A. (2013) Beta-Diversity of functional groups of woody plants in a tropical dry forest in Yucatan. PloS one, 8, e73660.

Lord, J.M. (2015) Patterns in floral traits and plant breeding systems on Southern Ocean Islands. AoB Plants, 7, plv095.

Lorite, J. (2016) An updated checklist of the vascular flora of Sierra Nevade (SE Spain). Phytotaxa, 261, 1–57.

Luke, Q. (2005) Annotated Checklist of the Plants of the Shimba Hills, Kwale District, Kenya. Journal of East African Natural History, 94, 5–120.

Luna-Jorquera, G., Fernández, C.E. & Rivadeneira, M.M. (2012) Determinants of the diversity of plants, birds and mammals of coastal islands of the Humboldt current systems: Implications for conservation. Biodiversity and Conservation, 21, 13–32.

Lynch, P.A., Ferguson, E.J. & Hynes, P. (1972) The vegetation of Red Mercury Island: Part I:The plant communities and a vascular plant species list. TANE, 18, 21–29.

Marquand, E.D. (1901) Flora of Guernsey and the lesser Channel Islands: namely Alderney, Sark, Herm, Jethou, and the adjacent islets. Dulau & Co, London, UK.

Marticorena, C., Squeo, F.A., Arancio, G. & Muñoz, M. (2008) Catálogo de la flora vascular de la IV Región de Coquimbo. Libro rojo de la flora nativa y de los sitios prioritarios para su conservación: Región de Atacama (eds F.A. Squeo, G. Arancio & J.R. Gutiérrez), pp. 105–142. Ediciones Universidad de La Serena La Serena.

Marticorena, C., Stuessy, T.F. & Baeza, C.M. (1998) Catalogue of the vascular flora of the Robinson Crusoe or Juan Fernández islands, Chile. Gayana Botánica, 55, 187–211.

Martínez-Laborde, J.B. (1998) A new report on the vascular flora of the island of Alborán (Spain). Flora Mediterrania, 8, 37–39.

Masharabu, T. Flore et végétation du Parc National de la Ruvubu au Burundi: diversité, structure et implications pour la conservation.

McClatchey, W., Thaman, R. & Vodonaivalu, S. (2000) A preliminary checklist of the flora of Rotuma with Rotuman names. Pacific Science, 54, 345–363.

McCrea, J. (2003) Inventory of the land conservation values of the Houtman Abrolhos Islands. Department of Fisheries, Government of Western Australia, Perth, Australia.

Medina, R., Reina-E, M., Herrera, E., Ávila, F.A., Chaparro, O. & Cortés-B., R. (2010) Catálogo preliminar da flora vascular dos bosques subandinos da cuchilla El Fara (Santander-Colômbia). Colombia Forestal, 13, 55–85.

Medjahdi, B., Ibn Tattou, M., Barkat, D. & Benabedli, K. (2009) La flore vasculaire des Monts des Trara (Nord Ouest Algérien). Acta Botanica Malacitana, 34, 57–75.

Memariani, F., Joharchi, M.R., Ejtehadi, H. & Emadzadeh, K. (2009) Contributions to the flora and vegetation of Binalood Mountain range, NE Iran: Floristic and chorological studies in Fereizi region. Ferdowsi University International Journal of Biological Sciences, 1, 1–17.

Miller, A.G. & Morris, M. (2004) Ethnoflora of the Soqotra Archipelago. Royal Botanic Garden, Edinburgh, UK.

Ministry of Environment (2012) The national red list 2012 of Sri Lanka: Conservation status of the fauna and flora, Colombo, Sri Lanka.

Miranda Freitas, A.M. de (2007) A Flora Fanerogâmica Atual do Arquipélago de Fernando de Noronha - Brasil. Universidade Estadual, Feira de Santana, Brasil.

Missouri Botanical Garden (2009) Shaw Nature Reserve. http://www.missouribotanicalgarden.org/visit/family-of-attractions/shaw-nature-reserve.aspx.

Missouri Botanical Garden (2015) Flora of the Jatun Sacha Biological Station: Preliminary Checklist. http://www.mobot.org/MOBOT/research/ecuador/jatun/checklist.shtml Accessed 15 January 15.

Moran, R. (1996) The flora of Guadalupe Island, Mexico. California Academy of Sciences, San Francisco, CA.

Morat, P., Jaffré, T., Tronchet, F., Munziger, J., Pillon, Y., Veillon, J.-M., Chalopin, M., Birnbaum, P., Rigault, F., Dagostini, G., Tinerl, J. & Lowry II, P.P. (2012) The taxonomic reference base Florical and characteristics of the native vascular flora of New Caledonia. Adansonia, 34, 179–221.

Nakamura, K., Suwa, R., Denda, T. & Yokota, M. (2009) Geohistorical and current environmental influences on floristic differentiation in the Ryukyu Archipelago, Japan. Journal of Biogeography, 36, 919–928.

Nassif, F. & Tanji, A. (2013) Floristic analysis of Marmoucha’s plant diversity (Middle Atlas, Morocco). LAZAROA, 34, 117–140.

Nationalpark Eifel (2015) Artenliste Farne und Blütenpflanzen. http://www.nationalpark-eifel.de/go/artenliste.html Accessed 11 March 15.

Nee, M. (2015) Flora de la Región del Parque Nacional Amboró, Bolivia. http://www.nybg.org/botany/nee/ambo/List.html Accessed 12 January 15.

Negoita, L., Fridley, J.D., Lomolino, M.V., Mittelhauser, G., Craine, J.M. & Weiher, E. (2016) Isolation-driven functional assembly of plant communities on islands. Ecography, 39, 1066–1077.

New Zealand Plant Conservation Network (2011) New Zealand's Flora. http://nzpcn.org.nz/page.asp?flora Accessed 18 March 11.

Newhook, F.J., Dickson, E.M. & Bennett, K.J. (1971) A botanical survey of some offshore islands of the Coromandel Peninsular. TANE, 17, 97–117.

Niering, W.A. (1963) Terrestrial ecology of Kapingamarangi Atoll, Caroline Islands. Ecological Monographs, 33, 131–160.

Nikolić, T. (2016) Flora Croatica Database. http://hirc.botanic.hr/fcd Accessed 12 February 16.

Noroozi, J. (2014) Checklist of the Flora of Iran based on the Flora Iranica (ed: Rechinger KH) and an update based on recent species records.

Norton, J., Majid, S.A., Allan, D., Al Safran, M., Böer, B. & Richer, R.A. (2009) An illustrated checklist of the flora of Qatar. Browndown Publications Gosport, Gosport, UK.

Notov, A.A. (2010) National park “Zavidovo”: vascular plants, bryophyte, lichens, Moscow.

Nowak, A. (2018) Vascular plants of Tajikistan.

NPS (2015) NPSpecies: Information on Species in National Parks. https://irma.nps.gov/NPSpecies/ Accessed 9 April 15.

Oggero, A.J. & Arana, M.D. (2012) Inventario de las plantas vasculares del sur de la zona serrana de Córdoba, Argentina. Hoehnea, 39, 171–199.

Ogle, C.C. (2002) Checklist of vascular plants of Sugar Loaf Islands. Wanganui plant list, 61, 1–8.

Oliveira-Filho, A.T. (2014) NeoTropTree: Flora arbórea da Região Neotropical: Um banco de dados envolvendo biogeografia, diversidade e conservação. http://www.icb.ufmg.br/treeatlan/ Accessed 12 February 16.

Ornduff, R. & Vasey, M.C. (1995) The vegetation and flora of the Marin Islands, California. Madrono, 42, 358–365.

Owiunji, I., Nkuutu, D., Kujirakwinja, D., Liengola, I., Plumptre, A.J., Nsanzurwimo, A., Fawcett, K., Gray, M. & McNeilage, A. The biodiversity of the Virunga Volcanoes.

Pal, D., Kumar, A. & Dutt, B. (2014) Floristic diversity of Theog Forest Division, Himachal Pradesh, Western Himalaya. Check list, 10, 1083–1103.

Pandey, R.P. & Diwakar, P.G. (2008) An integrated checklist flora of Andaman and Nicobar Islands, India. J. Econ. Taxon. Bot, 32, 403–500.

Pandža, M. (2002) Flora of the small islands of Murter. Natura Croatica, 11, 77–101.

Pandža, M. (2003) Flora of the island of Zirje and the small islands around it (eastern Adriatic coast, Croatia). Acta Botica Croatia, 62, 115–139.

Pandža, M. (2010) Flora parka prirode Papuk (Slavonija, Hrvatska). Šumarski list, 134, 25–43.

Pandža, M. & Milović, M. (2015a) Flora of the islets near Pakoštane (Dalmatia, Croatia). Natura Croatica, 24, 19–35.

Pandža, M. & Milović, M. (2015b) Flora of the Veliki Lagan and Mali Lagan islets (Dugi Otok island, Croatia). Natura Croatica, 24, 215–222.

Pandža, M., Milović, M., Kripina & Tafra, D. (2011) Vascular flora of the Vrgada islets (Zadar Archipelago, Eastern Adriatic). Natura Croatica, 20, 97–116.

Pandža, M. & Skvorc, Z. (2002) The flora of some uninhabited Sibenik Archipelago islands (Dalmatia, Croatia). Natura Croatica, 11, 367–385.

Panitsa, M., Bazos, I., Dimopoulos, P., Zervou, S., Yannitsaros, A. & Tzanoudakis, D. (2004) Contribution to the Study of the Flora and Vegetation of the Kithira Island Group: Offshore Islets of Kithira (S Aegean, Greece). Willdenowia, 34, 101–115.

Parks Canada (2015) Biotics Web Explorer. http://www.pc.gc.ca/apps/bos/BOSIntro_e.asp accessed 13 April 15.

Parris, B.S. & Latiff, A. (1997) Towards a pteridophyte flora of Malaysia: A provisional checklist of taxa. Malayan Nature Journal,50, 235–280.

Paula, S., Arianoutsou, M., Kazanis, D., Tavsanoglu, Ç., Lloret, F., Buhk, C., Ojeda, F., Luna, B., Moreno, J.M. & Rodrigo, A. (2009) Fire-related traits for plant species of the Mediterranean Basin. Ecology, 90, 1420.

Pawlaczyk, J. & Pawlaczyk, P. (2000) Vascular plants of Drawa National park and its neighbourhood. http://www.eko.org.pl/lkp/dpn/chckl_rosliny_sh.html Accessed 23 April 15.

Pedashenko, H. & Vassilev, K. (2014) Flora of Ponor Special Protection Area (Natura 2000), western Bulgaria. Acta zoologica bulgarica, 5, 33–60.

Pelser, P.B., Barcelona, J.F. & Nickrent, D.L. (2011) Co's Digital Flora of the Philippines. https://www.philippineplants.org/

Peña Nieto, E., Cárdenas Monroy, G., Zariñana Oronoz, C., Ocaña Nava, L., Robles Martínez, D., Peña Lomelí, A., Martínez Romero, J., Llerena Villalpando, F.A., Rugerio Galván, A.C. & Juárez Espejel, V. (2010) Inventarios Florísticos y Faunísticos de la Cuenca Alta del Río Lerma: Plan Maestro para la Restauración Ambiental de la Cuenca Alta del Río Lerma. Secretaría del Medio Ambiente, Agosto, Mexico.

Peña-Chocarro, M.d.C. (2010) Updated checklist of vascular plants of the Mbaracayú Forest Nature Reserve (Reserva Natural del Bosque Mbaracayú), Paraguay. Magnolia Press, Auckland, N.Z.

Pôle Flore Habitats (2015) Catalogue de la flore vasculaire de Rhône-Alpes. http://www.pifh.fr/pifhcms/index.php accessed 12 January 15.

Price, J.P. & Wagner, W.L. (2011) A phylogenetic basis for species–area relationships among three Pacific Island floras. American Journal of Botany, 98, 449–459.

Proctor, G.R. (1980) Checklist of the plants of Little Cayman. Geography and ecology of Little Cayman. Atoll Research Bulletin, 241, 71–80.

Proctor, G.R. (1985) Ferns of Jamaica: a guide to the Pteridophytes. British Museum (Natural History), London, UK.

Proctor, G.R. (1989) Ferns of Puerto Rico and the Virgin Islands. Memoirs of the New York Botanical Garden, New York, NY.

Programma de Conservación y Manejo (2006) Parque Nacional Cumbres de Monterrey, Mexico.

Pulu Keeling National Park (2004) Second Pulu Keeling National Park Management Plan.

Queensland Government (2014) Census of the Queensland flora 2014. https://data.qld.gov.au/dataset/census-of-the-queensland-flora-2014 accessed 5 February 15.

Rahman, A.H.M.M. (2013) Angiospermic Flora of Rajshahi District, Bangladesh. American Journal of Life Sciences, 1, 105.

Rahman, M.S., Hossain, G.M., Khan, S.A. & Uddin, S.N. (2015) An annotated Checklist of the Vascular Plants of Sundarban Mangrove Forest of Bangladesh. Bangladesh Journal of Plant Taxonomy, 22, 17–41.

Rakov, N.S., Saksonov S.V., Senator S.А. & Vasjukov V.M. (2014) Vascular plants of Ulyanovsk Region. Russian Academy of Sciences, Togliatti.

Raulerson, L. (2006) Checklist of plants of the Mariana Islands. University of Guam Herbarium Contribution, 37, 1–69.

Releford, J.S., Stevens, J., Bridges, K.W. & McClatchey, W.C. (2009) Flora of Rongelap and Ailinginae Atolls, Republic of the Marshall Islands. Atoll Research Bulletin, 572, 1–13.

Renvoize, S.A. (1975) A floristic analysis of the western Indian Ocean coral islands. Kew Bulletin, 30, 133–152.

Revushkin, A. (2014) Key to the flora of the Tomsk Region [Opredelitel rasteniy Tomskoy oblasty]. with minor corrections by Alexandr Ebel. Tomsk State University Press, Tomsk.

Robinson, A.C., Canty, P.D. & Fotheringham, D. (2008) Investigator group expedition 2006: flora and vegetation. Transactions of the Royal Society of South Australia, 132, 173–220.

Robinson, A.C., Canty, P.D., Wace, N.M. & Barker, R.M. (2003) The encounter 2002 expedition to the isles of St Francis, South Australia: flora and vegetation. Transactions of the Royal Society of South Australia, 127, 107–128.

Rossetto, E.F.S. & Vieira, A.O.S. (2013) Vascular Flora of the Mata dos Godoy State Park, Londrina, Paraná, Brazil. Check list, 9, 1020–1034.

Roux, J.P. (2009) Synopsis of the Lycopodiophyta and Pteridophyta of Africa, Madagascar and neighbouring islands. South African National Biodiversity Institute, Cape Town, South Africa.

Royal Botanic Gardens and Domain Trust (2017) PlantNET - The NSW Plant Information Network System. http://plantnet.rbgsyd.nsw.gov.au Accessed 16 September 16.

Royal Botanic Gardens Kew (2016) Seed information database (SID): v7.1. http://data.kew.org/sid/ Accessed 22 March 13.

Rundel, P.W., Dillon, M.O. & Palma, B. (1996) Flora and Vegetation of Pan de Azúcur National Park in the Atacama desert of Northern Chile. Gayana Bot, 53, 295–315.

Sachet, M.-H. (1962) Flora and vegetation of Clipperton Island. Proceedings of the California Academy of Sciences, 31, 249–307.

Sáez, L. & Rosselló, J.A. (2001) Llibre vermell de la flora vascular de les Illes Balears. Direcció General de Biodiversitat, Conselleria de Medi Ambient, Govern de les Illes Balears, Palma de Mallorca, Spain.

SANBI (2014) Plants of Southern Africa: An online checklist. http://posa.sanbi.org Accessed 9 March 15.

Sandbakk, B.E., Alsos, I.G., Arnesen, G. & Elven, R. (1996) The flora of Svalbard. http://svalbardflora.no/ Accessed 16 March 11.

Schaefer, H., Hardy, O.J., Silva, L., Barraclough, T.G. & Savolainen, V. (2011) Testing Darwin's naturalization hypothesis in the Azores. Ecology Letters, 14, 389–396.

Schönfelder, P. & Schönfelder, I. (1997) Die Kosmos-Kanarenflora: Über 850 Arten der Kanarenflora und 48 tropische Ziergehölze. Franckh-Kosmos, Stuttgart.

Scouppe, M. (2011) Composition floristique et diversité de la végétation de la zone Est du Parc National de Taï (Côte d’Ivoire). Master, Université de Genève.

Searle, J. & Madden, S. (2006) Flora assessment of South Stradbroke Island. Gold Coast City Council, Gold Coast City, Australia.

SEINet (2015) SEINet Flora Projects. http://swbiodiversity.org/seinet/projects/index.php Accessed 2 April 15.

Selvi, F. (2010) A critical checklist of the vascular flora of Tuscan Maremma (Grosseto province, Italy). Fl. Medit, 20, 47–139.

Senterre, B., Chew, M.Y. & Chung, R.C.K. (2015) Flora and vegetation of Pulau Babi Tengah, Johor, Peninsular Malaysia. Check list,11, 1714.

Seregin, A.P. (2013) New Flora of the Maeshchera National Park (Vladimir Oblast, Russia): Checklist, distribution atlas, peculiarities, and distributional changes in species over the last decade (2002–2012). ASTRA, Tula.

Sfair, J.C., Arroyo-Rodríguez, V., Santos, B.A. & Tabarelli, M. (2016) Taxonomic and functional divergence of tree assemblages in a fragmented tropical forest. Ecological Applications, 26, 1816–1826.

Shaheen, H., Qureshi, R., Akram, A., Gulfraz, M. & Potter, D. (2014) A preliminary floristic checklist of Thal Desert Punjab, Pakistan. Pakistan Journal of Botany, 46, 13–18.

Shaw, J.D., Spear, D., Greve, M. & Chown, S.L. (2010) Taxonomic homogenization and differentiation across Southern Ocean Islands differ among insects and vascular plants. Journal of Biogeography, 37, 217–228.

Shherbina, S. (2009) Flora of Vascular Plants of Central Siberian State Biosperic Reserve and neighboring territories. Turczaninowia, 12, 71–241.

Short, P.S., Albrecht, D.E., Cowie, I.D., Lewis, D.L. & Stuckey, B.M. (2011) Checklist of the vascular plants of the Northern Territory. Department of Natural Resources, Environment, The Arts and Sport, Darwin.

Silaeva, T.B., Chugunov, G.G., Kiryukhin, I.V., Ageeva, A.M., Vargot, E.V., Grishutkina, G.A. & Khapugin, A.A. Flora of the national park "Smolny". Mosses and vascular plants: annotated list of species [In Russian]. Commission of RAS for the Conservation of Biological Diversity [Комиссия РАН по сохранению биологического разнообразия].

Silantyeva, M. (2013) Synopsis of the flora of Altayskyi Krai [Konspekt flory Altayskogo kraya]. with minor corrections by Alexandr Ebel. Altay State University Press, Barnaul.

Singh, A. (2015) Observations on the vascular flora of Banaras Hindu University Main Campus, India. International Journal of Modern Biology and Medicine, 6, 48–87.

Skelin, M., Ljubičić, I., Skelin, I., Vitasović Kosić, I. & Bogdanović, S. (2014) The flora of Zečevo (Hvar Archipelago, Croatia). Agriculturae Conspectus Scientificus, 79, 85–91.

Slik, F.J.W., Arroyo-Rodríguez, V., Aiba, S.-I., Alvarez-Loayza, P., Alves, L.F., Ashton, P., Balvanera, P., Bastian, M.L., Bellingham, P.J., van den Berg, Eduardo, Bernacci, L., da Conceição Bispo, Polyanna, Blanc, L., Böhning-Gaese, K., Boeckx, P., Bongers, F., Boyle, B., Bradford, M., Brearley, F.Q., Breuer-Ndoundou Hockemba, M., Bunyavejchewin, S., Calderado Leal Matos, Darley, Castillo-Santiago, M., Catharino, Eduardo L M, Chai, S.-L., Chen, Y., Colwell, R.K., Robin, C.L., Clark, C., Clark, D.B., Clark, D.A., Culmsee, H., Damas, K., Dattaraja, H.S., Dauby, G., Davidar, P., DeWalt, S.J., Doucet, J.-L., Duque, A., Durigan, G., Eichhorn, K.A.O., Eisenlohr, P.V., Eler, E., Ewango, C., Farwig, N., Feeley, K.J., Ferreira, L., Field, R., de Oliveira Filho, Ary T, Fletcher, C., Forshed, O., Franco, G., Fredriksson, G., Gillespie, T., Gillet, J.-F., Amarnath, G., Griffith, D.M., Grogan, J., Gunatilleke, N., Harris, D.J., Harrison, R., Hector, A., Homeier, J., Imai, N., Itoh, A., Jansen, P.A., Joly, C.A., de Jong, Bernardus H J, Kartawinata, K., Kearsley, E., Kelly, D.L., Kenfack, D., Kessler, M., Kitayama, K., Kooyman, R., Larney, E., Laumonier, Y., Laurance, S., Laurance, W.F., Lawes, M.J., Amaral, Ieda Leao do, Letcher, S.G., Lindsell, J., Lu, X., Mansor, A., Marjokorpi, A., Martin, E.H., Meilby, H., Melo, Felipe P L, Metcalfe, D.J., Medjibe, V.P., Metzger, J.P., Millet, J., Mohandass, D., Montero, J.C., de Morisson Valeriano, Márcio, Mugerwa, B., Nagamasu, H., Nilus, R., Ochoa-Gaona, S., Onrizal, Page, N., Parolin, P., Parren, M., Parthasarathy, N., Paudel, E., Permana, A., Piedade, Maria T F, Pitman, N.C.A., Poorter, L., Poulsen, A.D., Poulsen, J., Powers, J., Prasad, R.C., Puyravaud, J.-P., Razafimahaimodison, J.-C., Reitsma, J., Dos Santos, João Roberto, Roberto Spironello, W., Romero-Saltos, H., Rovero, F., Rozak, A.H., Ruokolainen, K., Rutishauser, E., Saiter, F., Saner, P., Santos, B.A., Santos, F., Sarker, S.K., Satdichanh, M., Schmitt, C.B., Schöngart, J., Schulze, M., Suganuma, M.S., Sheil, D., da Silva Pinheiro, Eduardo, Sist, P., Stevart, T., Sukumar, R., Sun, I.-F., Sunderand, T., Suresh, H.S., Suzuki, E., Tabarelli, M., Tang, J., Targhetta, N., Theilade, I., Thomas, D.W., Tchouto, P., Hurtado, J., Valencia, R., van Valkenburg, Johan L C H, van Do, T., Vasquez, R., Verbeeck, H., Adekunle, V., Vieira, S.A., Webb, C.O., Whitfeld, T., Wich, S.A., Williams, J., Wittmann, F., Wöll, H., Yang, X., Adou Yao, C Yves, Yap, S.L., Yoneda, T., Zahawi, R.A., Zakaria, R., Zang, R., de Assis, Rafael L, Garcia Luize, B. & Venticinque, E.M. (2015) An estimate of the number of tropical tree species. Proceedings of the National Academy of Sciences of the United States of America, 112, 7472–7477.

Sloover, J.R. de & Liégeois, S. (1997) Eagle Island Flora and Vegetation (Queensland, Australia). Bulletin du Jardin Botanique National de Belgique, 66, 347–383.

de Sloover, JR & Liégeois, S. (1997) Eagle Island flora and vegetation (Queensland, Australia). Nationale Plantentuin van België, 66, 347–383.

SLUFG (2015) Rote Liste und Artenliste Sachsen: Farn-und Samenpflanzen. Sächsisches Landesamt für Umwelt, Landwirtschaft und Geologie, Dresden.

Smith, A.C. (1979-1996) Flora Vitiensis nova: a new Flora of Fiji (spermatophytes only). Pacific Tropical Botanical Garden (Lawaii, Hawaii).

Smithsonian Institution (2003) A checklist of Trees, Shrubs, Herbs, and Climbers of Myanmar. Contributions from the United States National Herbarium, 45, 1–590.

St John, H. (1988) Census of the flora of the Gambier Islands, Polynesia. Pacific Plant Studies 43, Honolulu, Hawaii.

Stace, C.A., Ellis, R.G., Kent, D.H. & McCosh, D.J. (2003) Vice-county Census Catalogue of the vascular plants of Great Britain, the Isle of Man and the Channel Islands. Botanical Society of the British Isles, London, UK.

Stalmans, M. (2006) Tinley's plant species list for the Greater Gorongosa ecosystem, Moçambique. Unpublished report by International Conservation Services to the Carr Foundation and the Ministry of Tourism.

Stalter, R. & Lamont, E.E. (2005) The Historical and Extant Flora of Great Gull Island, New York. Journal of the Torrey Botanical Society, 132, 628–634.

Stalter, R. & Lamont, E.E. (2006) The historical and extant flora of Sable Island, Nova Scotia, Canada. Journal of the Torrey Botanical Society, 133, 362–374.

Stoddart, D.R. & Fosberg, F.R. (1994) Flora of the Phoenix Islands, central Pacific. Atoll Research Bulletin, 393, 1–60.

Stone, B.C. (1959) The flora of Namonuito and the Hall Islands. Pacific Science, 13, 88–104.

Strahm, W.A. The conservation and restoration of the flora of Mauritius and Rodrigues, Reading, University of.

Sykes, W.R. (1970) Contributions to the flora of Niue. Bulletin. Department of Scientific and Industrial Research, New Zealand, 200, 321.

Sykes, W.R., West, C.J., Beever, J.E. & Fife, A.J. (2000) Kermadec Islands flora-special edition: a compilation of modern material about the flora of the Kermadec Islands. Manaaki Whenua Press, Landcare Research.

Takahashi, H., Barkalov, V.Y., Gage, S., Joneson, S., Ilushko, M. & Zhuravlev, Y.N. (2002) A floristic study of the vascular plants of Raikoke, Kuril Islands. Acta Phytotaxonomica et Geobotanica, 53, 17–33.

Takahashi, H., Barkalov, V.Y., Gage, S., Semsrott, B., Ilushko, M. & Zhuravlev, Y.N. (1999) A preliminary checklist of the vascular plants of Chirinkotan, Kuril Islands. Journal of Phytogeography and Taxonomy, 47, 131–137.

Takahashi, H., Barkalov, V.Y., Gage, S., Semsrott, B., Ilushko, M. & Zhuravlev, Y.N. (2006) A floristic study of the vascular plants of Kharimkotan, Kuril Islands. Bulletin of the Hokkaido University Museum, 3, 41–66.

Takahashi, H., Barkalov, V.Y., Gage, S. & Zhuravlev, Y.N. (1997) A preliminary study of the flora of Chirpoi, Kuril Islands. Acta Phytotaxonomica et Geobotanica, 48, 31–42.

Tamis, W.L.M., van der Meijden, R., Runhaar, J., Bekker, R.M., Ozinga, W.A., Odé, B. & Hoste, I. (2004) Standard List of the Flora of the Netherlands 2003. Gorteria, 30, 101–195.

Tatewaki, M. (1957) Geobotanical studies on the Kurile Islands. Acta Horti Gotoburgensis, 21, 43–123.

Taylor, G.A. (1991) Flora and fauna of Plate (Motunau) Island, Bay of Plenty. TANE, 33, 113–120.

Taylor, G.A. (1995) Flora and Fauna of Needle Rock, eastern Coromandel. TANE, 35, 51–56.

Taylor, G.A. & Cameron, E.K. (1990) Kauwahaia Island, Te Henga, West Auckland. Auckland Botanical Society Journal, 45, 71–77.

Taylor, G.A. & Lovegrove, T.G. (1997) Flora and vegetation of Stanley (Atiu) Island, Mercury Islands. TANE, 36, 85–111.

Taylor, G.A., Lovegrove, T.G., Miskelly, CM, McFadden, I & Whitaker, A.H. (1987) An ecological survey of small islands in the Mercury Group. TANE, 32, 151–167.

Taylor, R. (2006) Straight through from London: the Antipodes and Bounty Islands, New Zealand. Heritage Expeditions New Zealand, Christchurch, New Zealand.

Tela Botanica (2015) Base de données des Trachéophytes de France métropolitaine (bdtfx). http://www.tela-botanica.org/bdtfx.

Telford, I.R.H. (1993a) Cocos (Keeling) Islands: Species lists. http://www.environment.gov.au/biodiversity/abrs/online-resources/flora/50/ accessed 7 April 11.

Telford, I.R.H. (1993b) Coral Sea Islands Territory: Species lists. http://www.environment.gov.au/biodiversity/abrs/online-resources/flora/50/ accessed 7 April 11.

Ter Steege, H., Vaessen, R.W., Cardenas-Lopez, D., Sabatier, D., Antonelli, A., Oliveira, S.M. de, Pitman, N.C.A., Jorgensen, P.M. & Salomao, R.P. (2016) The discovery of the Amazonian tree flora with an updated checklist of all known tree taxa. Scientific reports, 6, 29549.

Thaman, R.R., Fosberg, F.R., Manner, H.I. & Hassall, D.C. (1994) The flora of Nauru. Atoll Research Bulletin, 392, 1–233.

Thaman, R.R., Keppel, G., Watling, D., Thaman, B., Gaunavinaka, T., Naikatini, A., Bolaqace, N., Sekinoco, E. & Masere, M. (2005) Nasoata mangrove island, the PABITRA coastal study site for Viti Levu, Fiji Islands. Pacific Science, 59, 193–204.

Thiombiano, A., Schmidt, M., Dressler, S., Ouédraogo, A. & Hahn, K. (2012) Catalogue des plantes vasculaires du Burkina Faso. Boissiera: mémoires des Conservatoire et Jardin botaniques de la Ville de Genève, 65, 1–391.

Thomas, J. (2011) Plant diversity of Saudi Arabia: Flora checklist. http://plantdiversityofsaudiarabia.info/biodiversity-saudi-arabia/flora/Checklist/Cheklist.htm accessed 14 January 16.

Tonkov, S., Pavlova, D., Atanassova, J., Nedelcheva, A. & Marinova, E. (2004) Floristis catalogue of the nature reserve Rilomanastirska Gora (Central Rila Mountains). I. The locality Kirilova. University of Sofia.

Tressens, S.G., Keller, H.A. & Revilla, V. (2008) Las plantas vasculares de la reserva de uso múltiple Guaraní, Misiones (Argentina). Boletín de la Sociedad Argentina de Botánica, 43, 273–293.

Trigas, P. (2015a) Plants of the Aegean: Aegean islands. unpublished.

Tropicos (2015a) Catálogo de las Plantas Vasculares de Bolivia. Tropicos. Missouri Botanical Garden, St. Louis.

Tropicos (2015b) Catalogue of the Vascular Plants of Ecuador. http://www.tropicos.org/Project/CE Accessed 22 October 15.

Tropicos (2015c) Catalogue of the Vascular Plants of the Department of Antioquia (Colombia). Tropicos. Missouri Botanical Garden, St. Louis.

Tropicos (2015d) Flora de Nicaragua. Tropicos. Missouri Botanical Garden, St. Louis.

Tropicos (2015e) Listado de la Flora del Parque Nacional Madidi, Bolivia. Tropicos. Missouri Botanical Garden, St. Louis.

Tropicos (2015f) Panama Checklist. Tropicos. Missouri Botanical Garden, St. Louis.

Tropicos (2015g) Paraguay Checklist. Tropicos. Missouri Botanical Garden, St. Louis.

Tropicos (2015h) Peru Checklist. Tropicos. Missouri Botanical Garden, St. Louis.

Trusty, J.L., Kesler, H.C. & Delgado, G.H. (2006) Vascular flora of Isla del Coco, Costa Rica. Proceedings of the California Academy of Sciences, 57, 247–355.

Turner, I.M. (1995) A catalogue of the vascular plants of Malaya. Gardens' Bulletin, Singapore.

Tutul, E., Uddin, M.Z., Rahman, M.O. & Hassan, M.A. (2009) Angiospermic flora of Runctia sal forest, Bangladesh. I. Liliopsida (Monocots). Bangladesh Journal of Plant Taxonomy, 16, 83–90.

UIB (2007) Herbario virtual del Mediterráneo Occidental. http://herbarivirtual.uib.es/cas-med/ Accessed 7 August 12.

University of Greifswald (2010) FloraGREIF - Virtual Flora of Mongolia. http://greif.uni-greifswald.de/floragreif/ Accessed 2 February 16.

University of Kent (2012) Cook Islands Biodiversity and Ethnobiology Database. http://cookislands.pacificbiodiversity.net/cibed/dbs/search.html Accessed 12 April 12.

USDA & NRCS (2015) The PLANTS Database. http://plants.usda.gov Accessed 24 April 15.

van Vreeswyk, A.M.E., Payne, A.L., Leighton, K.A. & Hennig, P. (2004) An inventory and condition survey of the Pilbara region, Western Australia. Department of Agriculture.

Vander Velde, N. (2003) The vascular plants of Majuro atoll, Republic of the Marshall Islands. Atoll Research Bulletin, 503, 1–141.

Vanderplank, S.E. (2010) The Vascular Flora of Greater San Quintín, Baja California, Mexico. CGU Theses & Dissertations.

Veklich, T.N. (2009) Flora of the Norsky Nature Reserve (Amur region). Russian Academy of Sciences, Far East Division, Blagoveshensk.

Velarde, E., Wilder, B.T., Felcer, R.S. & Ezcurra, E. (2014) Floristic diversity and dynamics of Isla Rasa, Gulf of California - A globally important seabird island. Botanical Sciences, 92, 89–101.

Velarde, E.P., Guzmán, R.C. & Koch, S.D. (2008) Plantas vasculares y vegetación de la parte alta del Arroyo Agua Fría, municipio de Minatitlán, Colima, México. Acta Botanica Mexicana, 84, 25–72.

Velayos, M., Barberá, P., Cabezas, F.J., La Estrella, M.d., Fero, M. & Aedo, C. (2014) Checklist of the Vascular Plants of Annobón (Equatorial Guinea). Phytotaxa, 171, 1–78.

Velayos Rodríguez, M. (2016) Flora de Guinea Equatorial. http://www.floradeguinea.com/ Accessed 14 February 16.

VicFlora (2016) Flora of Victoria. http://vicflora.rbg.vic.gov.au Accessed 30 November 16.

Viciani, D., Gonnelli, V., Sirotti, M. & Agostini, N. (2010) An annotated check-list of the vascular flora of the “Parco Nazionale delle Foreste Casentinesi, Monte Falterona e Campigna”(Northern Apennines Central Italy). Webbia, 65, 3–131.

Vogt, C. (2011) Composición de la Flora Vascular del Chaco Boreal, Paraguay. I. Pteridophyta y Monocotiledoneae. Steviana, 3, 13–47.

Wace, N.M. (1961) The vegetation of Gough Island. Ecological Monographs, 31, 337–367.

Wace, N.M. & Dickson, J.H. (1965) The terrestrial botany of the Tristan da Cunha Islands. Philosophical Transactions of the Royal Society of London B Biological Sciences, 249 (759), 273–360.

Wagner, W.L., Herbst, D.R. & Lorence, D.H. (2005) Flora of the Hawaiian Islands website. http://botany.si.edu/pacificislandbiodiversity/hawaiianflora/ Accessed 16 October 10.

Wagner, W.L. & Lorence, D.H. (2002) Flora of the Marquesas Islands website. http://botany.si.edu/pacificislandbiodiversity/marquesasflora/index.htm Accessed 28 November 14.

Walker, T.A. (1991) Pisonia Islands of the Great Barrier Reef: Part III. Changes in the vascular Flora of Lady Musgrave Island. Atoll Research Bulletin, 350, 31–41.

Walker, T.A., Chaloupka, M.Y. & King, B.R. (1991) Pisonia Islands of the Great Barrier Reef: Part II. The vascular Floras of Bushy and Redbill Islands. Atoll Research Bulletin, 350, 24–30.

WCSP (2014) World Checklist of Selected Plant Families. http://apps.kew.org/wcsp/home.do Accessed 1 December 14.

Webster, G.L. & Rhode, R.M. (2001) Plant diversity of an Andean cloud forest: inventory of the vascular plants of Maquipucuna, Ecuador. Publications in Botany, 82, 1–228.

Wellington Botanical Society (2008) Native vascular plants of Great Barrier Island. Wellington Botanical Society, Wellington, New Zealand.

Wester, L. (1985) Checklist of the vascular plants of the northern Line Islands. Atoll Research Bulletin, 187, 1–38.

Western Australian Herbarium (2017) FloraBase - the Western Australian Flora. https://florabase.dpaw.wa.gov.au/ Accessed 16 September 16.

Whistler, W.A. Botanical survey of Diego Garcia, Chagos Archipelago, British Indian Ocean Territory. Appendix E1.

Whistler, W.A. (1983) Vegetation and flora of the Aleipata Islands, Western Samoa. Pacific Science, 37, 227–249.

Whistler, W.A. (1998) A study of the rare plants of American Samoa. US Fish and Wildlife Service, Honolulu, Hawaii.

Whistler, W.A. (2012) Botanical survey of the Ringgold Islands, Fiji. Allertonia, 11, 1–28.

Wieringa, J.J. (2016) Flora of Gabon. unpublished.

Woodroffe, C.D. (1986) Vascular plant speciesarea relationships on Nui Atoll, Tuvalu, Central Pacific: a reassessment of the small island effect. Australien Journal of Ecology, 11, 21–31.

Wright, A.E. (1976) The vegetation of Great Mercury Island. TANE, 22, 23–49.

Yakubov, V.V. (2010) Illustrated Flora of the Kronotsky Reserve (Kamchatka): Vascular plants. Institute of Biology and Soil Science, Vladivostok.

Yineger, H., Kelbessa, E., Bekele, T. & Lulekal, E. (2008) Floristic composition and structure of the dry afromontane forest at Bale Mountains National Park, Ethiopia. SINET: Ethiopian Journal of Science, 31, 103–120.

Yuncker, T.G. (1959) Plants of Tonga. Bishop Museum Bulletin, 220, 1–283.

Zanne, A.E., Lopez-Gonzalez, G., Coomes, D.A., Ilic, J., Jansen, S., Lewis, S.L., Miller, R.B., Swenson, N.G., Wiemann, M.C. & Chave, J. (2009) Data from: Towards a worldwide wood economics spectrum. Dryad Digital Repository. https://doi.org/10.5061/dryad.234

Zanne, A.E., Tank, D.C., Cornwell, W.K., Eastman, J.M., Smith, S.A., FitzJohn, R.G., McGlinn, D.J., O'Meara, B.C., Moles, A.T., Reich, P.B., Royer, D.L., Soltis, D.E., Stevens, P.F., Westoby, M., Wright, I.J., Aarssen, L., Bertin, R.I., Calaminus, A., Govaerts, R., Hemmings, F., Leishman, M.R., Oleksyn, J., Soltis, P.S., Swenson, N.G., Warman, L. & Beaulieu, J.M. (2014) Three keys to the radiation of angiosperms into freezing environments. Nature, 506, 89–92.

ZDSF & SKEW (2014) Info Flora: Artenliste Schweiz 5×5 km. https://www.infoflora.ch/de/daten-beziehen/artenliste-5×5-km.html Accessed 13 February 15.

Zhang, S.-B., Slik, J.W.F., Zhang, J.-L. & Cao, K.-F. (2011) Spatial patterns of wood traits in China are controlled by phylogeny and the environment. Global Ecology and Biogeography, 20, 241–250.

Zhang, S.T., Zhen Du, G. & Chen, J.K. (2004) Seed size in relation to phylogeny, growth form and longevity in a subalpine meadow on the east of the Tibetan plateau. Folia Geobotanica, 39, 129–142.

Zotz, G. (2013) The systematic distribution of vascular epiphytes - a critical update. Bot. J. Linn. Soc., 171, 453–481.

Zuloaga, F.O., Morrone, O. & Belgrano, M. (2014) Catálogo de las Plantas Vasculares del Cono Sur. http://www.darwin.edu.ar/Proyectos/FloraArgentina/fa.htm Accessed 16 March 15.

Zvyagintseva, K.O. (2015) An annotated checklist of the urban flora of Kharkiv. Kharkiv National University, Kharkiv, Ukraine.

Абрамова, Л.А. & Волкова, П.А. (2011) Сосудистые растения Байкальского заповедника: (Аннотированный список видов). Флора и фауна заповедников, 117, 1–112.

Антипова, E.M. (2012) Флора внутриконтинентальных островных лесостепей Средней Сибири. гос. пед. ун-т им. В.П. Астафьева, Красноярск.

Артемов, И.А. (2012) Определитель растений Катунского биосферного заповедника. Russian Academy of Sciences, БАРНАУЛ.

Гаджиев, В.Д. & Юсифов, Э.Ф. (2003) Флора и растительность Кызылагачского заповедника и их биоразнообразие. Национальная Академия Наук Азербайджана, Баку.

Евстигнеев, О.И. & Федотов, Ю.П. (2007) Флора сосудистых растений заповедника "Брянский лес". Гос. природ. биосфер. заповедник Брян. лес, Брянск.

Куликов, П.В. & Кирсанова, О.Ф. (2012) Сосудистые растения заповедника "Денежкин камень": (Аннотированный список видов). Флора и фауна заповедников, 119, 1–140.

Лактионов, А.П., Пилипенко, В.Н., Глаголев, С.Б. & Лактионова, Н.А. (2008) Сосудистые растения заповедника «Богдинско-Баскунчакский» (Аннотированный список видов): Флора и фауна заповедников. Флора и фауна заповедников, 113, 1–113.

Миркина, Б.М. (2010) Флора и растительность Национального парка «Башкирия». Russian Academy of Sciences.

Морозова, О.В., Царевская, Н.Г. & Белоновская, Е.А. (2010) Сосудистые растения национального парка «Валдайский». Флора и фауна заповедников, 7, 1–98.

Хапугин, A.A. (2013) Сосудистые растения Ромодановского района Республики Мордовия (конспект флоры), Saransk.

